# Allometry of cell types in planarians by single cell transcriptomics

**DOI:** 10.1101/2023.11.01.565140

**Authors:** Elena Emili, Alberto Pérez-Posada, Maria D. Christodoulou, Jordi Solana

**Affiliations:** Department of Biological and Medical Sciences, Oxford Brookes University, Oxford, UK; Department of Statistics, University of Oxford, Oxford, UK

## Abstract

Allometry explores the relationship between an organism’s body size and its various components, offering insights into ecology, physiology, metabolism, and disease. The cell is the basic unit of biological systems, and yet, the study of cell type allometry remains relatively unexplored. Single-cell RNA sequencing (scRNA-seq) provides a promising tool for investigating cell type allometry. Planarians, capable of growing and degrowing following allometric scaling rules, serve as an excellent model for such studies. We used scRNA-seq to examine cell type allometry in asexual planarians of different sizes, revealing that they consist of the same basic cell types but in varying proportions. Notably, the gut basal cells are the most responsive to changes in size, suggesting a role in energy storage. We capture the gene regulatory programs of distinct cell types in response to size. This research sheds light on the molecular and cellular aspects of cell type allometry in planarians and underscores the utility of scRNA-seq in such investigations.

## Introduction

Allometry studies how body parts, structures, or processes in an organism relate to its overall body size (*1*). It is important for understanding ecology, physiology, metabolism, disease, and other aspects of organismal biology. Scaling laws often govern allometry, such as the Kleiber law that relates metabolism to body size using a 3/4 power function (*2*).

While allometry has been classically studied using whole organism parts, the study of allometry at the cell type level is still vastly unexplored. Cells are the fundamental units of life, serving as building blocks of multicellular organisms. They can be classified in different types, based on their morphology, function, or gene expression patterns (*3*). However, the exact definition of a cell type is still a topic of debate and controversy among researchers (*4–10*). Despite the importance of cell types in understanding biological systems, the study of how cell types vary with size (i.e. cell type allometry) is still a largely unexplored field. Advances have been made using morphological cell type classifications and markers (*11*) but these are typically limited to one or few cell types. Whether cell types vary with size, and how, is still a question, for most cell types, in most organisms.

The study of cell types has been enhanced by single cell methods (*12–14*), particularly single cell RNA sequencing (scRNA-seq) (*15–17*), which measures the expression of hundreds to thousands of individual transcripts in thousands of cells. Each cell type is characterised by specific expression of effector and regulatory genes, allowing clustering algorithms to reveal the presence of cell populations. However, it is still unclear how these populations correspond to cell types as opposed to other cell states such as stress, differentiation or developmental states. Despite this, scRNA-seq enables quick and simple classification of cells into largely interpretable populations, such as neurons, muscle cells, and epidermal cells.

Single cell analysis techniques have potential for studying cell type allometry, but several challenges remain to be addressed. Cell dissociation techniques can cause stress responses and biases (*18–20*), leading to cell death and differential survival. Including different samples can also lead to batch effects. However, fixative cell dissociation approaches like ACME (*21*) can address the first concern, while combinatorial single cell transcriptomic approaches like SPLiT-seq (*21, 22*) allow for sample multiplexing and convenient multi-sample experiments. Combining ACME and SPLiT-seq could therefore enhance the study of cell type allometry.

Planarians are an interesting model for studying cell type allometry because apart from growing when fed, they can degrow when starved (*23–25*). This process follows allometric scaling rules (*11, 26, 27*). They exhibit a wide range of body sizes, with sexual populations typically being larger than asexual ones. Romero and Baguñà examined cell dissociations from animals of varying sizes to study 13 basic cell types based on their morphology (*26*). They showed that body size increases mostly by increases in cell numbers, with quantitative changes in the proportions of cell types. For example, larger planarians have a decrease in neuron proportion and an increase in fixed parenchymal cells. Thommen and coworkers established that the metabolic rate in planarians also follows the Kleiber law (*27*). Interestingly, larger planarians have more cells, but their cellular size scales with mass following a 3/4 power relationship. This indicates that larger planarians have many more cells, but that at least some of these cells are also larger. It remains unknown if this applies to all cell types or if it is largely driven by one or few cell types. Planarians are an excellent model for single cell biology (*21, 28–35*), but the allometry of their cell types has not been investigated yet with these methods.

Here, we aimed to investigate the allometry of cell types in asexual planarians using scRNA-seq across body size categories. To minimise batch effects and enable an integrated analysis, we used SPLiT-seq to multiplex planarians of different sizes in a single experiment. Our findings reveal that asexual planarians of different sizes are made of the same cell types in different proportions. Specifically, smaller planarians exhibit a higher proportion of neurons and fewer parenchymal cells. Unlike previous studies, scRNA-seq has greater resolution on which types of neurons and parenchymal cells vary the most. Notably, further to cell proportions, scRNA-seq allows us to access cell type specific genetic programs. Our data shows that epidermal cells and the recently discovered basal cells are the ones that respond more dynamically to organism size at the transcriptomic level. Overall, our results show that scRNA-seq using ACME and SPLiT-seq is a powerful method to study cell type allometry that can be applied to virtually any organism. Our data reveals the basic principles of cell type scaling at the cellular and molecular level.

### A single cell approach for studying cell type allometry

We aimed at generating one scRNA-seq experiment to multiplex asexual planarians of different sizes (Figure 1A). We first selected asexual planarians from our cultures and classified them in sizes by visual inspection. To corroborate our inspection, we took low resolution images using a general field camera and measured the area of each individual using Fiji (*36*) (Figure 1B). This was done to avoid the stress of monitoring the animals under a bright light prior to single cell transcriptomics. We undertook the image capture immediately prior to cell dissociation, to avoid that animals vary in size further after capture and to avoid fission events that would modify the sizes. We used ACME (*21*) to dissociate the different sized animals in 4 independent cell dissociations. We initially aimed at classifying 4 body sizes, but found that the two higher size classes overlapped significantly and decided to consider them as one class. We therefore considered 3 classes of body size, termed Large (L), Medium (M) and Small (S). Measuring the area of the individual animals revealed a good size separation in 3 classes (Figure 1C, Supplementary File 1). The mean size of our classes was 8.04, 4.02 and 1.24 mm2 for L, M and S planarians respectively. The larger planarian measured 12.98 mm2 and the smaller measured 0.30mm2, encompassing a ∼43-fold difference in area. In the average populations the fold difference between L and S planarians was 6.45. To achieve comparable cell numbers we used 35, 50 and 100 animals of L, M and S sizes respectively (Figure 1D).

**Figure 1.**
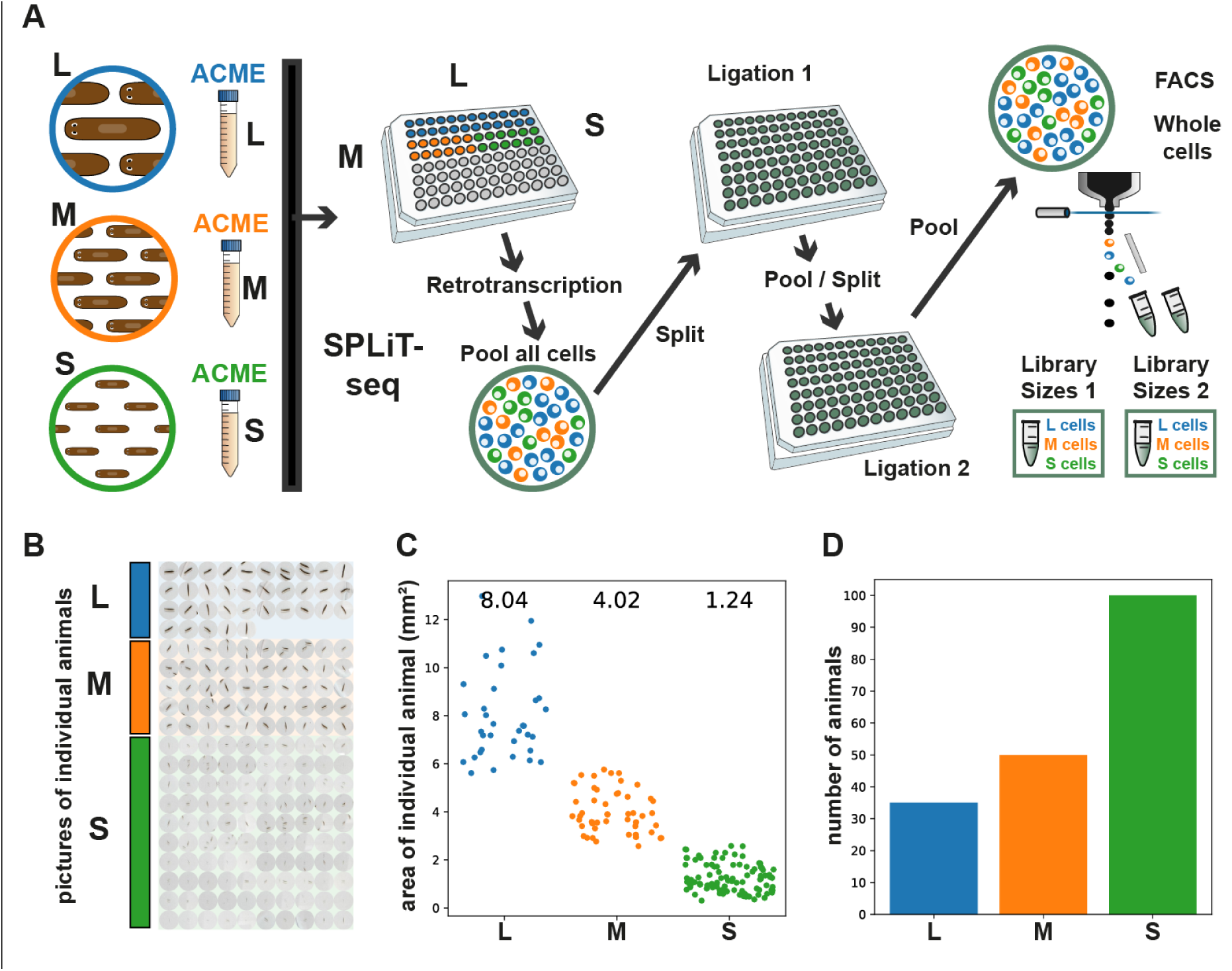
Animal selection, measurement and single cell transcriptomic approach. A: Experimental workflow. Small, Medium, and Large planarians were ACME-dissociated in separate tubes and subjected to multiplexed combinatorial barcoding for sequencing (see Methods). B: Low-resolution images of selected animals taken with a standard field camera. C: Area of individual animal per size class. D: Number of animals per size class.

### A cell type atlas of planarian body sizes

We then subjected these cell dissociations to scRNA-seq, multiplexing the different classes using RT barcodes in SPLiT-seq’s first combinatorial barcoding round (Figure 1A). This step allows demultiplexing the cells coming from each sample in the data analysis steps. After this first round, cells are pooled and every subsequent barcoding, sorting and library preparation step is performed in pools of cells containing all samples, therefore minimising batch effects. Furthermore, SPLiT-seq is very robust against ambient RNA, as the successive pooling and centrifugation steps select the material encapsulated in cells, while the supernatants (with any soluble ambient RNA released from the cell suspension) are eliminated in each round. We implemented FACS sorting after the 3^rd^ round of combinatorial barcoding, a step that allows us to sort barcoded whole cells into the lysis buffer, prior to the 4^th^ round of barcoding. This step drastically reduces the presence of broken cells and particles of cellular debris in the experiment. We obtained two sub-libraries which were subjected to the 4^th^ round of barcoding during library preparation. In these conditions, the cost of a single SPLiT-seq experiment is around ∼£750 in enzymes, FACS sorting time and library preparation reagents, making it an affordable scRNA-seq approach. We sequenced the library pool using Illumina NovaSeq technology, to obtain 715 million paired-end reads.

We then processed these reads through our SPLiT-seq analysis pipeline (*21, 37*). The final dataset contained 28,738 cells, consistent with the ∼19Kx2 cells observed in each of the 2 FACS sorted samples. Leiden clustering revealed the presence of ∼40-73 cell clusters, depending on clustering resolution. We selected resolution 3 (Figure 2A), which resolved 64 cell populations, largely corresponding to planarian well known cell types. We grouped these cell type identities in broad groups by hierarchical clustering (Figure 2B). We annotated cell clusters using previous literature (Figure 2C, Supplementary Files 2-3, Supplementary Figure 1) and cross referencing published markers with markers obtained from this experiment (Supplementary Files 4-6). Our dataset resolves 3 types of differentiated muscle cells, 3 epidermal and gastrodermal differentiation states (Supplementary Figure 1), and up to 17 neuronal cell populations. This resolution reveals clusters containing relatively rare cell types such as the glia (Figure 2D, Supplementary Figure 1, 0.19 %), the psd+ cells (Figure 2D, 0.26%), but fails to resolve the photoreceptors which are clustered together with other cholinergic neurons. We annotated cluster 13 as the basal cells (Figure 2A-D), which had been clustered together with phagocytes and goblet cells in previous studies (*21, 28, 30*). Neoblasts were clustered in 3 major clusters, including one cluster that we called “committed neoblasts” as it expressed markers of differentiation to gastrodermal and epidermal types (Supplementary Figure 1). Finally, a number of small cell types (10 clusters, 315 cells, 1.1 %, Figure 2A-B, Supplementary Files 4-6) had unspecific markers or no significant markers were interpreted as putative doublets, and were left unannotated in the final dataset. SPLiT-seq resulted in a low UMI and gene per cell content (Supplementary Figure 2A-D), but allowed us to distinguish 55 annotated cell clusters. The average mean counts and genes per cluster vary from cluster to cluster (Supplementary Figure 2A-D). To elucidate whether this variation is technical, or reflects biological properties of the cells instead, we studied the relationship between counts and cells at the cell and cluster level. The number of genes detected in each cell largely correlates with the number of counts obtained, as expected (Supplementary Figure 2E). However, at the cluster level averages, some clusters deviate from the correlation: secretory cell types have more counts than other cell clusters with similar gene counts, consistent with the idea that they express a few genes at higher levels (Supplementary Figure 2F). The cluster with more gene counts and UMI counts is nanos+ germ cell progenitors, consistent with previous publications (*21*). These observations indicate that the variations in cluster counts and genes are biological signals. Altogether, these results show that our ACME + SPLiT-seq approach reliably detects dozens of cell clusters.

**Figure 2.**
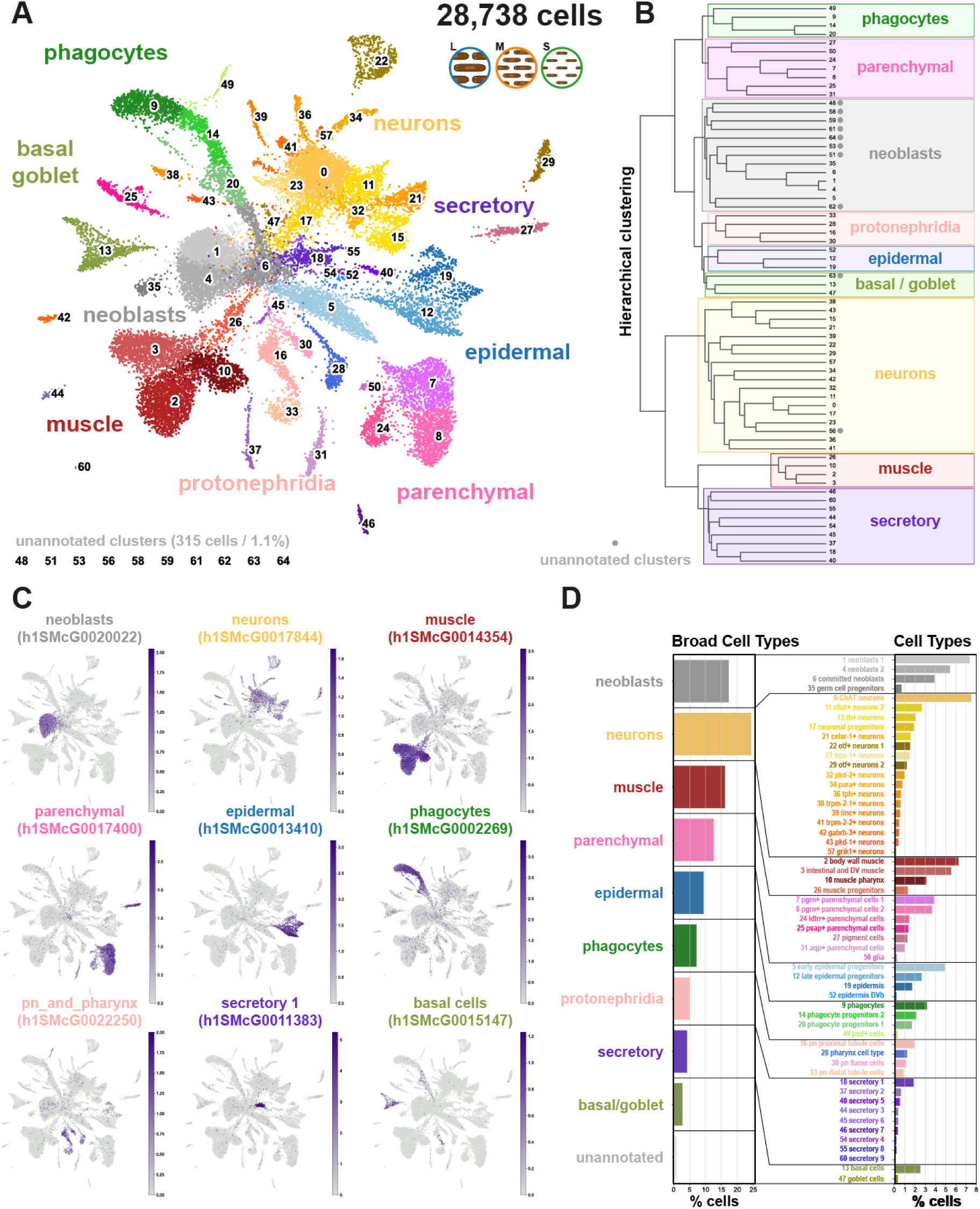
The *Schmidtea mediterranea* Sizes dataset. A: UMAP visualisation of the *Schmidtea mediterranea* Sizes single-cell atlas with clusters coloured according to their cell cluster classification. B: Hierarchical clustering dendrogram of cell clusters to define broad group identities. Grey dots denote the unannotated clusters. C: UMAP features plots of markers of the major broad types. D: Cell cluster percentage over the total at the broad cell groups and the cell cluster levels.

### Asexual planarians of different sizes are made of the same cell types

We aimed to determine if all body sizes have the same cell types, building on previous studies that found no evidence of size-dependent cell types (*26*). However, the increased resolution of our dataset raised the possibility of identifying more specialised cell types that could be size dependent. For instance, larger organisms may have specialised energy storage types. Furthermore, processes like sexualization are known to lead to the formation of new organs (e.g. gonads and reproductive apparatuses). Therefore, to determine if there are body size specific cell types we assessed if all detected cell populations were present across all body sizes.

We first examined qualitatively the distribution of L, M and S cells in the UMAP space. Interpreting UMAPs in such a way is controversial as dimensionality reduction techniques and low-dimensional embedding of single-cell data are known to introduce distortions of the dataset (*38*). On the other hand, our multiplexed analysis to minimise batch effects allows us to expect a homogeneous distribution of all three samples, except for biological differences. Therefore, we investigated UMAP distribution differences as an exploratory analysis to motivate further inquiry. In our dataset, L, M and S cells were homogeneously distributed throughout the UMAP except in regions corresponding to the basal cells (cluster 13), the epidermis (cluster 19) and secretory 4 (cluster 54) (Figure 3A). In order to investigate if these effects are underlied by biological differences or stochastic variation due to dimensionality reduction, we explored the parameter space of our PCA, k-nearest neighbours embedding and UMAP (Supplementary Figure 3). This analysis revealed that UMAP differences in basal cells were observable in all analyses with at least 55 Principal Components, and in spite of the k-nearest neighbour parameter. This strongly suggests that the differences in the basal cell cluster are of biological source and motivated further inquiry.

**Figure 3.**
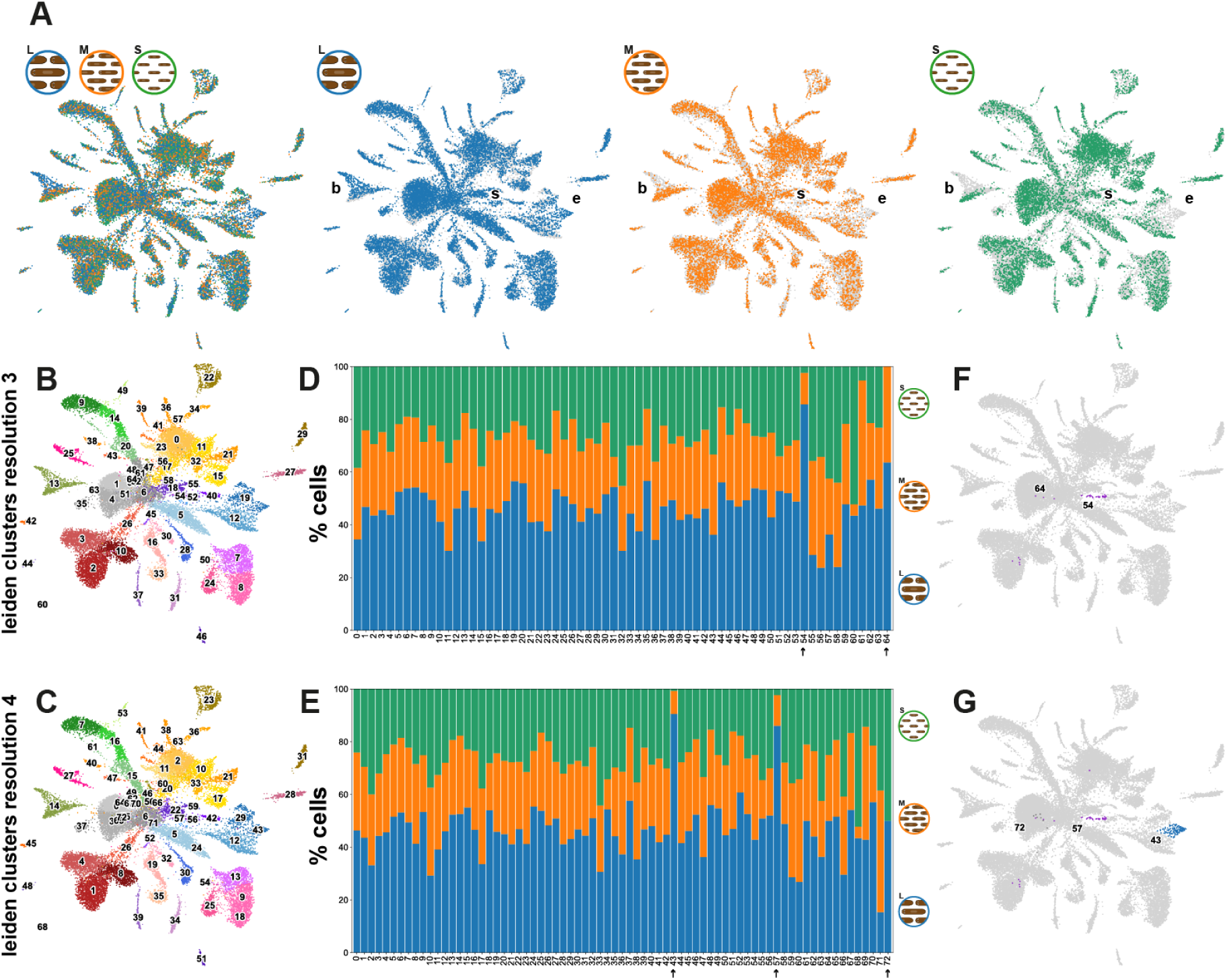
Cell type frequency in planarians of different sizes. A: UMAP of L, M and S cells in the UMAP space merged and by individual body size category. B-C: UMAP based on Leiden clustering at resolution 3 (B) and resolution 4 (C). D-E: stacked barplot of cell percentages based on Leiden clustering at resolution 3 (D) and resolution 4 (E). Arrows point to the clusters with the lowest percentages. F-G: UMAPs highlighting cell clusters 54 and 64 of resolution 3 (F) and 43, 57 and 72 of resolution 4 (G) with the lowest percentages.

We then asked if these differences translated into size category specific cell clusters. To decouple this analysis from the resolution parameter of the cell clustering algorithm, we performed this analysis using 4 different resolutions of the leiden algorithm (1–4) (Figure 3B-C). This analysis showed that all 3 size categories have cells from all clusters except for the smallest clusters, i.e. cluster 64 of resolution 3 (11 cells, Figure 3D) and cluster 72 of resolution 4 (8 cells, Figure 3E). This strongly suggests that this is a sampling effect due to the small cell numbers of these clusters. Further to that, cluster 54 of resolution 3 and clusters 43 and 57 of resolution 4 are largely made of L and M cells (Figure 3D-G), with only a few S cells (1 in each case). These clusters correspond to the differences observed before on UMAP space. Cells from cluster 19 of resolution 3, corresponding to the epidermis (Figure 3B), are sub-clustered in 2 clusters of resolution 4, clusters 29 and 43 (Figure 3C). The latter cluster is made majoritarily of cells from the Large sample (125 L, 12 M and 1 S) and expresses markers of both epidermis and late epidermal progenitors (Supplementary File 4). Cluster 13 of resolution 3, containing the basal cells (Figure 3B), remains one cluster (14) at resolution 4 (Figure 3C). Therefore, we argue that the basal cells are one cell type and the differences observed in UMAP space correspond to differences in the state of basal cells. Overall, this analysis shows that asexual planarians of our 3 size categories are made of the same essential cell clusters but that differences in cell type abundance may exist.

### Differences in cell proportions in planarians of different sizes at single cell resolution

We then analysed the differences in cell type frequency between L, M and S planarians. In order to visualise these differences in the UMAP space we plotted the percentage ratios of each cell cluster in large vs small planarians (Figure 4A). The values examined correspond to the logarithmic ratio between the percentage of each cluster in L samples vs S samples. In this representation, a log2 ratio of 1 indicates that the cell cluster percentage is double in L than in S samples.

**Figure 4.**
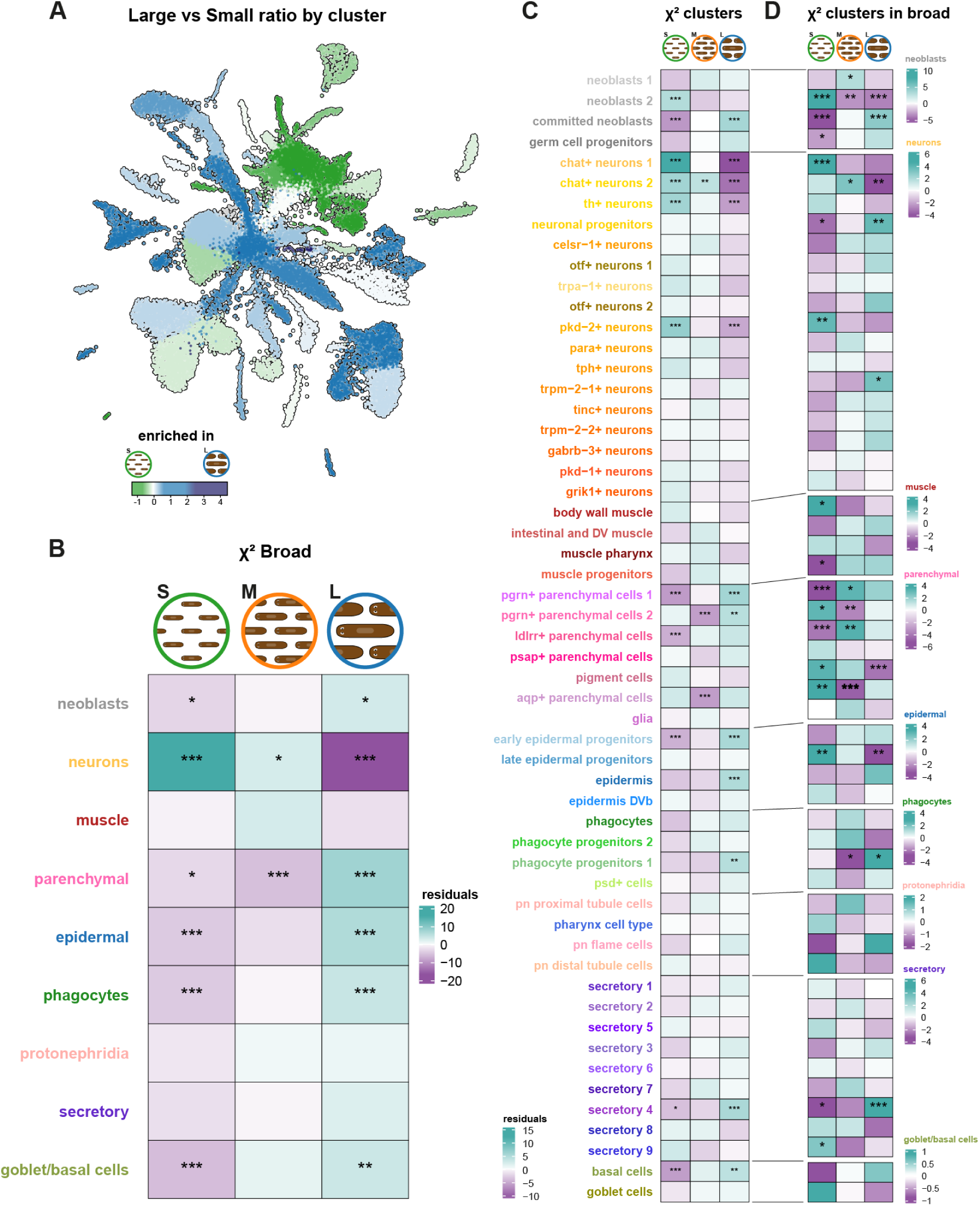
Differences in cell proportions in planarians of distinct sizes. A: UMAP visualising the percentage log2 ratios of cell clusters in L vs. S planarians. B: Heatmap representation of the results of the chi-square test examining the association between S, M, and L in broad groups. C: Heatmap representation of the results of the chi-square test examining the association between S, M, and L in the 55 annotated clusters. D: Heatmap representation of the results of the chi-square test examining the association between S, M, and L in each cluster of the broad group. Colour intensity represents the residuals from the chi-square test; green tones indicate increased frequency of cell types (surplus of observed counts in that category) and purple tones indicate decreased frequency of cell types (deficit in observed counts in that category). Significance levels are denoted by asterisks (*) based on the Benjamini-Hochberg adjusted p-values. * represents p_adj < 0.05. ** represent p_adj < 0.01. *** represent p_adj < 0.001.

We analysed the frequencies using a chi squared test with a Benjamini-Hochberg p-value adjustment (Supplementary File 7). To elucidate if our results compare well with those published by microscopical observation, we first analysed the dataset grouped by broad type category. This grouping made our categories largely comparable with those analysed by Baguñà and Romero (*26*). Our analysis revealed an increased frequency of neurons in small planarians, and a decrease in large planarians (Figure 4B), consistent with the microscopy findings. On the contrary, we observed significantly increased frequencies in large, and decreased in small, for the neoblasts, parenchymal, epidermal, phagocytes, and goblet/basal cell broad categories (Figure 4B). The increases in parenchymal, phagocytes and goblet/basal are also consistent with the reported increases of fixed parenchymal cells, gastrodermal and goblet cells in the microscopy data. Our analysis revealed tendencies inconsistent with those reported by Baguña and Romero for the neoblast and epidermis broad groups, which were described to be more frequent in small planarians.

Next, we repeated this analysis in our 55 annotated clusters (Figure 4C). This analysis revealed that the clusters neoblasts 2 and committed neoblasts showed opposite trends, with the neoblasts 2 overrepresented in small planarians, and the committed neoblasts underrepresented in small planarians and overrepresented in large planarians instead. This can potentially explain the discrepancy with the microscopy dataset, as this dataset did not include progenitor cells. It is unclear how this study classified cells that have started to differentiate, but likely these cells already displayed differentiation markers and were classified within their differentiated type category. Next, we examined the individual neuron clusters. We found significant overrepresentation in small and underrepresentation in large for the chat+ neurons clusters 1 and 2, the th+ neurons and the pkd-2+ neurons. Most other neuron clusters had similar tendencies but were not significant in this analysis. Among the parenchymal broad group, we found that the pgrn+ clusters 1 and 2 were significantly enriched in large planarians and significantly underrepresented in small and medium planarians. Furthermore, the ldlrr+ cells were also found significantly reduced in small planarians. Interestingly, the aqp+ cells were overrepresented in medium planarians, and was the only cluster with this behaviour. The biology and function of the parenchymal cells is still enigmatic, and the biological significance of this observation is unclear. From the epidermal cells, we found significant enrichment in large planarians for the early epidermal progenitors and the epidermis. We also found a significant enrichment of phagocyte progenitors 1, secretory 4 cells and the basal cells. Altogether, this analysis shows that single cell transcriptomics has higher resolution and is able to detect significant differences at the cell cluster level, beyond the broad type categories.

Next, we analysed each of these clusters, but compared to their broad category (Figure 4D). This analysis aimed to identify cell types that behave contrary to their broad type. Our results corroborated the contrary tendencies of neoblasts 2 and committed neoblasts. We also observed that trpm-2-2+ neurons were enriched in the neuronal subgroup of large planarians, a trend opposite to that of neurons as a group. Other clusters behaved similarly, but were not significant. Of note, the neuronal progenitors also have a significant enrichment in the neuronal broad group of large planarians. This analysis also revealed a significant enrichment of body wall muscle in small planarians when compared to the muscle broad group. Within the parenchymal broad group, we found highly significant underrepresentation of pgrn+ and ldlrr+ cells in small planarians and underrepresentation of pigment cells in large planarians. Collectively, these results show that studying allometry with single cell transcriptomics has a high resolution and identifies several cell types that behave contrary to their broad type categories.

### Head cell types are enriched in small planarians, and intestinal and parenchymal cell types are enriched in large planarians

We then tested if the patterns observed could be explained by anatomical features. In order to investigate this we compared the percentage ratios of each cell cluster in large vs small planarians sorted by Broad Type (Figure 5A) to the same values sorted by Body Region (Figure 5B). We assigned cell clusters to anatomical features based on previous literature (Supplementary File 2). In brief, “body wall” corresponded to body wall muscle (*39*), pigment cells (*40–42*), protonephridia distal tubule cells (*43–45*), and the epidermis (*33, 46*). “Intestine” cell types (*47–49*) included phagocytes, goblet cells, basal cells (*47*), psd+ cells (*30*) and intestinal muscle (*39*). All neuronal types as well as the cell cluster described as glia (*50, 51*) were classified as “nervous system”. “Pharynx” contains the pharynx cell type (*30*) and the muscle of the pharynx (*30*). All other cell clusters were considered part of the “parenchyma”, i.e. the space between the body wall and the intestine. This included parenchymal cells (excluding pigment and glia, located in the body wall and the nervous system), secretory cell types, neoblasts and progenitor clusters. This analysis revealed that body regions largely explain the observed frequencies, with nervous system cell types being broadly enriched in small planarians, and intestinal and parenchymal cell types being broadly enriched in large planarians. In order to gain more insight into this notion we examined the most extreme values of these distributions and seeked to validate these observations by querying previously published in situ hybridisation experiments (*28*). Cluster 42 cells were the nervous system cell cluster with the highest L to S ratio, a pattern dissimilar to other neuronal cell types (Figure 5B). In fact, cluster 42 cells were not enriched in heads but are distributed throughout the body (Figure 5C). Cluster 55 secretory cells had the lowest enrichment in L samples (Figure 5B). Interestingly, their distribution showed accumulation in head regions (Figure 5D), explaining the enrichment in S samples, similar to that of neuron types. In stark contrast, cluster 44 secretory cells had the strongest enrichment in L samples (Figure 5B). Consistently, we observed that these cells are broadly distributed throughout the parenchyma, but depleted from the head region (Figure 5E). Finally, we examined cluster 24, containing ldlrr+ parenchymal cells (*28, 30*) as it showed prominent enrichment in L samples (Figure 5B). The expression pattern revealed that these cells were associated with gut tissues (Figure 5F). Altogether, these analyses show that small planarians have comparatively more head cell types, including nervous and secretory types, and large planarians instead quantitatively contain more parenchymal and intestinal cell types.

**Figure 5.**
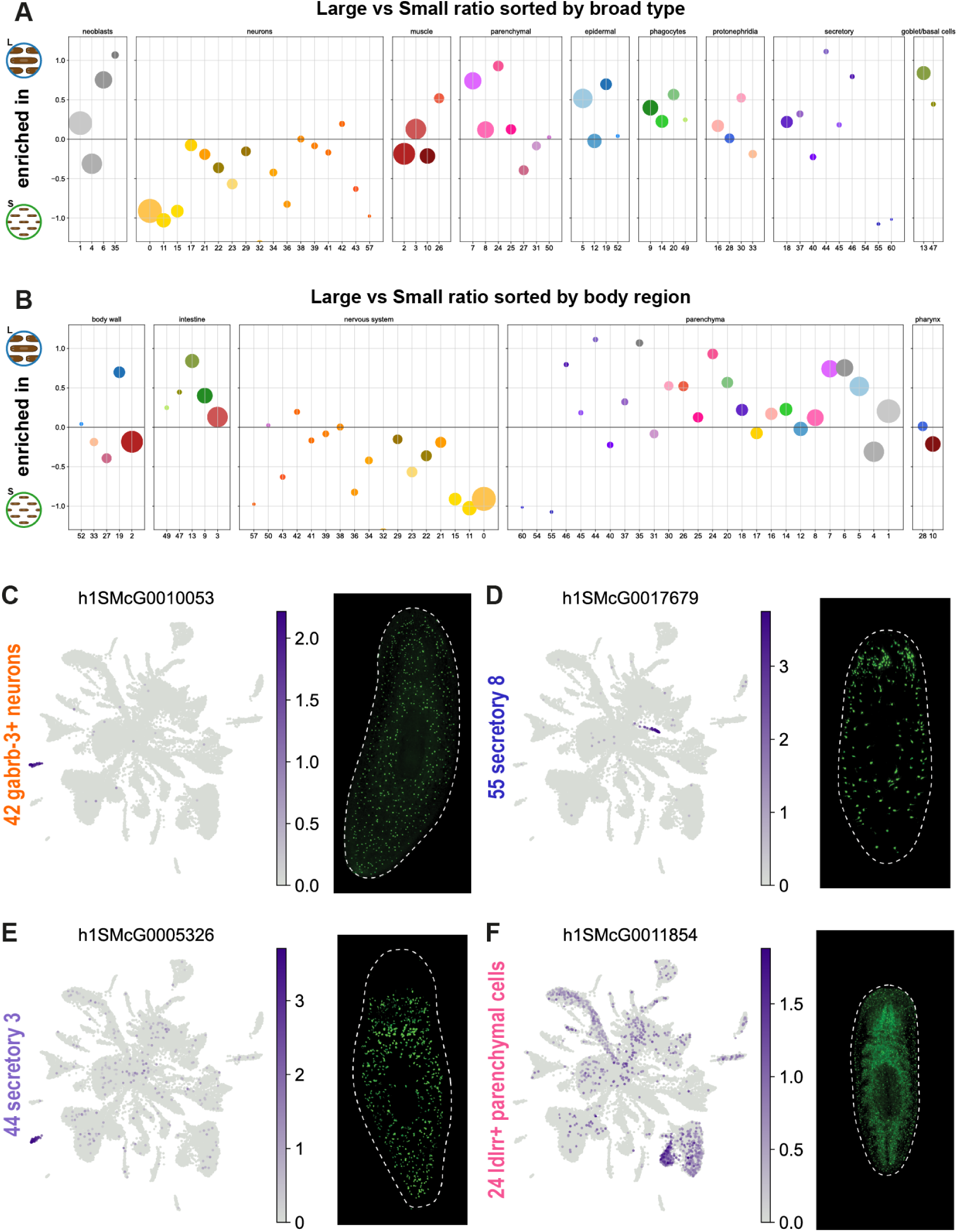
Variations in cell proportions by anatomical features. A: Percentage ratios of cell clusters in L vs S planarians sorted by broad group. Dot size represents cluster size in cell number. B: Percentage ratios of cell clusters in L vs S planarians sorted by body region. Dot size represents cluster size in cell number. In both plots cluster 54, corresponding to secretory 4 cells, lies outside of the plotted scale with a high enrichment in L. C: left - Feature plot of marker gene of *gabrb-3*+ neurons (cluster 42); right - *in situ* picture from the same gene showing homogeneous distribution throughout the body. D: left - Feature plot of marker gene of secretory 8 (cluster 55). right - *in situ* picture from the same gene showing an enrichment of this cell type in the head. E: Feature plot of marker gene of secretory 3 (cluster 44). right - *in situ* picture from the same gene showing the exclusion of this cell type from the head region. F: Feature plot of marker gene of *ldrr*+ parenchymal cells (cluster 24). right - *in situ* picture from the same gene showing the association of this cell type with the gut tissue. All *in situ* pictures retrieved from the database digiworm (https://digiworm.wi.mit.edu/).

### Differential gene expression analysis reveals cell type specific size-related gene programs

We then aimed at determining if cell types change their gene expression patterns in response to animal size. For this analysis we focused on large and small samples. Taking full advantage of our multiplexed single cell approach, we aggregated cluster information to generate pseudobulk (*52–54*) count tables for each specific cluster (Figure 6A). We used the two libraries of the experiment (Figure 1A) as pseudoreplicates. The two libraries come from the same experiment and can only be considered technical replicates of the tagmentation and 4th barcoding step. However, we reasoned that these are indeed made of completely different cells. It has been recently shown that pseudoreplication (i.e. computationally distributing cells in random groups to obtain replicate information) works well for single cell differential gene expression analysis (*55*). Arguably, this is because in single cell approaches there is an additional level of biological replication in the single cell barcoding. We analysed the 65 independent cluster-aggregated count tables using DEseq2 (*52*) to identify genes differentially regulated by size (Supplementary File 8). To elucidate if there are cell types that respond more dynamically to size differences we examined the relationship between cluster size and the number of differential genes detected (Figure 6B) in each cell cluster. The number of differential genes increased with cluster size, since this is correlated with the number of reads that go into the analysis, maximising its power. However, two clusters lie outside of this distribution, with a much higher number of differentially regulated genes: the epidermis (cluster 19) and the basal cells (cluster 13). These are the clusters identified in the UMAP analysis, likely explaining the UMAP differences. These lists overlapped by only 3 genes (Figure 6C), indicating that the differences in gene expression of epidermal and basal cells are largely independent and cell type specific. This analysis shows that the epidermis and the basal cells are the planarian cell types that respond more dynamically to animal size at the gene expression level, with largely independent genetic programs.

**Figure 6.**
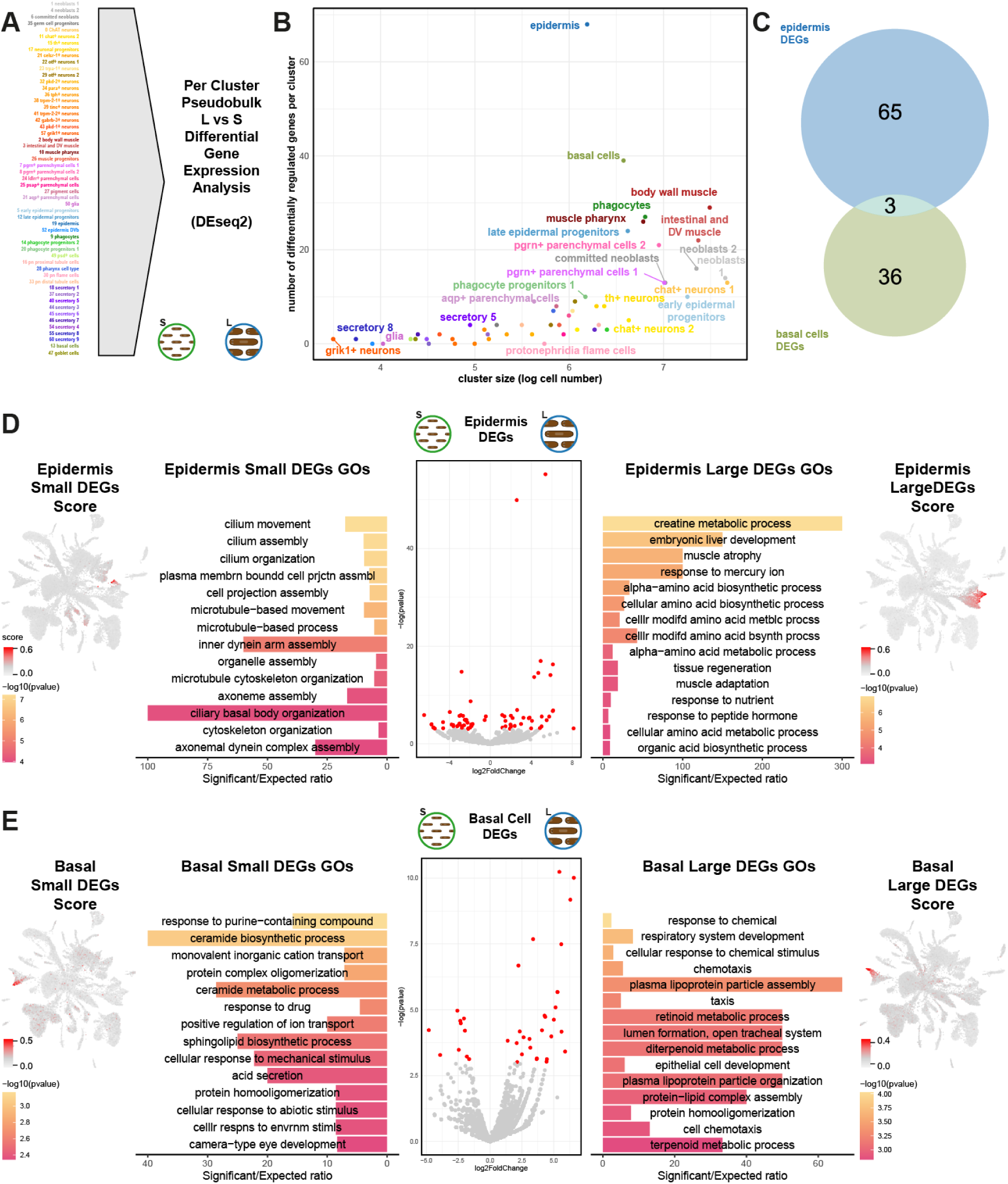
Size-dependent cell type specific genetic programs in large and small planarians. A: Generation of pseudo-bulk count tables for the 55 annotated cell clusters. B: Differential gene expression analysis with DEseq2. Scatter plot showing the number of differentially regulated genes per cluster compared to the log-transformed cell count within each cluster. C: Overlap between differentially expressed genes detected in epidermis and basal cells. D: Differential gene expression analysis, GO term enrichment analysis and UMAP visualisation of gene scores, comparing the epidermis of L vs. S planarians. Bar size indicates ratio of significant/expected fraction of annotated genes with a given GO term. Colour gradient indicates adjusted p-value (Fisher’s exact test). Scored expression of the group of differentially regulated genes in red. E: Differential gene expression analysis, GO term enrichment analysis and UMAP visualisation of gene scores, comparing the basal of L vs. S planarians. Colours, bars and scores as in D.

**Figure 7.**
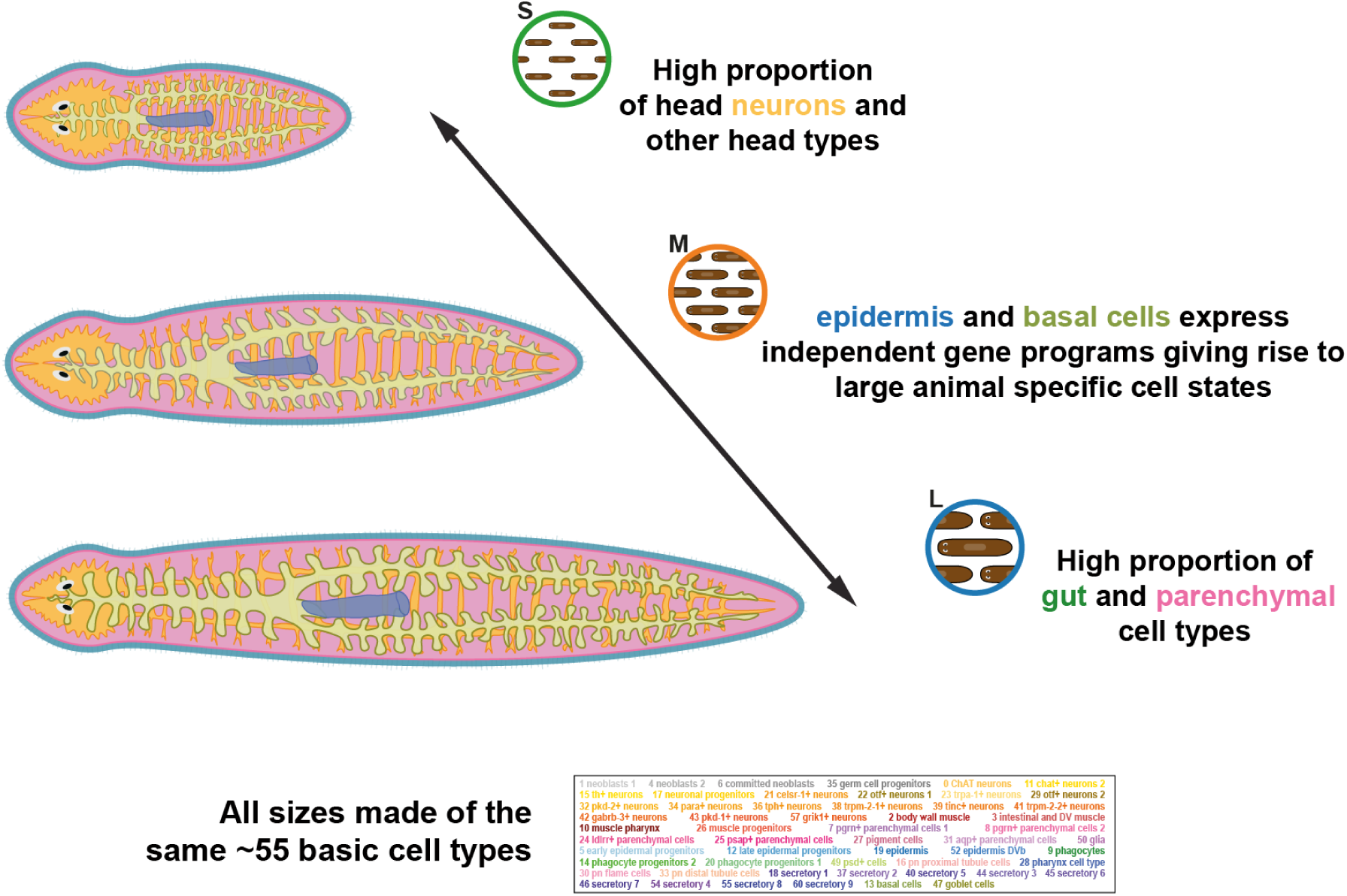
Graphical representation of the cellular and molecular basis for cell type allometry in planarians. Representation of the key allometric variation depicted via single-cell combinatorial barcoding approach. Presence of the ∼55 basic cell types in all planarian sizes. High proportion of neurons of the head region and other non-neuronal head cell types in S planarians. Independent gene programs in epidermis and basal cells give rise to L planarian specific cell states. High proportion of gut and parenchymal cell types in L planarians.

To examine the genes that are differentially regulated in the epidermis and the basal cells we obtained volcano plots and analysed the GO term enrichment of up- and down-regulated genes (Figures 6D-E). Genes differentially expressed in epidermal cells from small animals were enriched in cilium and other related GOs (Figure 6D). Genes differentially expressed in epidermal cells of large animals were enriched in creatine metabolism and other metabolic processes, suggesting that in large planarians epidermal cells carry out metabolic functions (Figure 6D). Key genes in this enrichment are AGAT enzymes (h1SMcG0009479, h1SMcG0009480, h1SMcG0011148), which have been already described in planarian late epidermal progenitors (*28, 30, 31, 33, 56*). Genes differentially expressed in small basal cells were enriched in ceramide sphingolipid biosynthesis genes (Figure 6E). Genes differentially expressed in basal cells of large animals were enriched in lipoprotein particle assembly, retinoid and terpenoid metabolism genes (Figure 6E).

Finally, to elucidate if these genes are exclusively expressed in the epidermis and basal cell clusters or if contrarily they are expressed more generally in other clusters as well, we examined their UMAP feature plots (Supplementary Figure 4). We found that the differentially expressed genes are expressed in other cell types. We then scored the expression of these gene sets to obtain a general view of their grouped expression. This analysis revealed that differentially expressed genes in epidermis and basal cells are enriched in the UMAP areas of differential distribution (Figure 3A), suggesting that these genes explain these differences. Altogether, these analyses show that epidermal and basal cells are the cell types that are most responsive to animal size as they exist in two states with their specific gene expression patterns, largely related to their metabolism. On one hand, large planarians have epidermal cells that express late epidermal progenitor genes, suggesting that their differentiation process is longer. On the other hand, basal cells in large planarians express lipid metabolism genes. This is consistent with the accumulation of lipids around the basal area of the gut described previously (*27*), although the existence of the basal cells was only shown later (*47*).

## Discussion

Cell type allometry is a key question in developmental biology, but it has remained mysterious in many organisms due to the lack of cell type markers (antibodies, RNA probes). Furthermore, cell quantification with methods of immunohistochemistry and/or *in situ* hybridisation is challenging at full body scale. Cell type identification based on microscopic observation presents a subjective component. Finally, these methods can only identify a few cell types and lack resolution.

Here we showed that single cell transcriptomics solves these challenges by providing a method to identify and quantify all major cell types in different samples. Beyond that, scRNA-seq allows accessing the gene expression patterns of each cell type. To achieve this level of resolution it is key to use fixation methods such as ACME (*21*) and multiplexing approaches such as SPLiT-seq (*22*). These methods will open the door to studying cell type allometry and other cell type quantitative studies in virtually every organism.

Planarians are an excellent model to study cell type allometry because of their ability to grow and degrow with food availability. Previous studies have already shown that neurons are enriched in small planarians and that gut and parenchymal cell types are enriched in large planarians. Our study confirmed these trends, but provided higher resolution. For instance, we were able to identify neuron types that are not enriched in small planarians and secretory types enriched in heads. Our results show that the ratios for each cell type are largely explained by their anatomical location, despite their broad cell type identity, with small planarians having an enrichment of head cell types.

Baguña and Romero showed that asexual planarians were made of the same basic 13 broad cell types in varying proportions. We confirm these results, but at a higher resolution of ∼55 annotated cell clusters. Single cell transcriptomics is changing the way we think about cell types, showing that granularity is key to their definition (*9*). We were able to show that there are no cell clusters specific to large or small asexual planarians at our current level of resolution. Remarkably, we were able to distinguish up to 17 types of neurons, 7 types of parenchymal cells and 9 types of secretory cells, and found them all represented in all size categories. While further subdivision of these types might lead to the identification of types specific to small or large planarians in the future, the question would then be if these correspond to cell types or if they constitute different cell states of the same specific cell types.

Importantly, we identify cell states characteristic of large planarians for the epidermis and the basal cells. We argue that these represent states instead of types because 1) they are just defined by a few quantitative differences in gene expression and 2) they share general markers of their cell type. Increasing the resolution of the Leiden clustering algorithm can cluster these cells out, but at these high resolutions other cell types are divided in clusters that lack meaningful markers and are largely explained by noise. Our results delve further into the definitions of cell type and cell state, and will offer insight to other researchers investigating the nature of cell type identity.

Crucially, we are able to access the gene expression patterns of these cell types and states, and identify differentially regulated genes. Further studies will identify more gene expression differences, enabled by deeper resolutions, larger cell numbers and increased number of samples and replicates. This will allow us to study their transcriptional regulation, including the transcription factors that regulate the differences. Combinatorial barcoding scRNA-seq methods such as SPLiT-seq and other single cell analysis methods will be key to decode the cellular and molecular principles of cell type allometry, as well as other developmental paradigms.

Our results are largely consistent with those of Baguñà and Romero’s microscopic study, except for 2 key broad groups: the neoblasts, and the epidermis. Key to understanding this discrepancy is the fact that the microscopy study did not identify progenitor cells. We were able to identify 7 progenitor clusters, the majority of which are enriched in large samples. This also applies to the committed neoblast cluster. It is unclear whether Baguñà and Romero classified progenitor cells with neoblasts or with their corresponding cell fate broad group. The overrepresentation of progenitors in large planarians raises interesting hypotheses about the dynamics of cellular differentiation. For instance the differentiated epidermis contains a group of cells that clusters together with the mature epidermis but contains late epidermal progenitor cell markers, and is highly enriched in large planarians. This could indicate a longer time of differentiation for epidermal cells in large planarians. Future research will address the dynamics of cell differentiation in planarians and other organisms using single cell transcriptomics.

The most dynamic cell type in planarians of different sizes are the basal cells, as they change both their frequency and their gene expression patterns. This type was only described recently (*47*) with proposed metabolic functions. In previous single cell transcriptomic dataset, the basal cells are found clustered together with other gut cell types (*30*), primarily goblet cells (*21*), due to their similarity. Likely, the basal cells host the size-dependent lipid accumulations described by Thomenn *et al*. (*27*). Our data shows that basal cells exist in two states characterised by their gene expression patterns. Basal cells in large animals express lipid particle synthesis and assembly genes. Our results will open the door to further studies about the biology of this enigmatic planarian cell type.

## Availability of Data and Materials

The datasets supporting the conclusions of this article are available in: Code: https://github.com/scbe-lab/planarian_cell_type_allometry GEO: GSE246681

## Acknowledgments

We thank Robert Hedley at the Flow Cytometry Facility at the Dunn School of Pathology (University of Oxford) and Vincent Mason for technical assistance. We thank Jochen Rink and Luca Pandolfini for early access to the new *Schmidtea mediterranea* genome and annotation. We thank all other members of the Stem Cell Biology and Evolution team for discussion and support.

## Authors’ Contributions

JS and EE conceived the study and designed the experiments. EE generated cell dissociations and performed single cell transcriptomic experiments. EE, AP-P and JS performed bioinformatic single cell analyses. MC performed the statistical analysis. JS and EE wrote the manuscript and generated the figures, with contributions from all other authors. All authors read and approved the final version of the manuscript.

## Supplementary Figure Legends

**Supplementary Figure 1.**
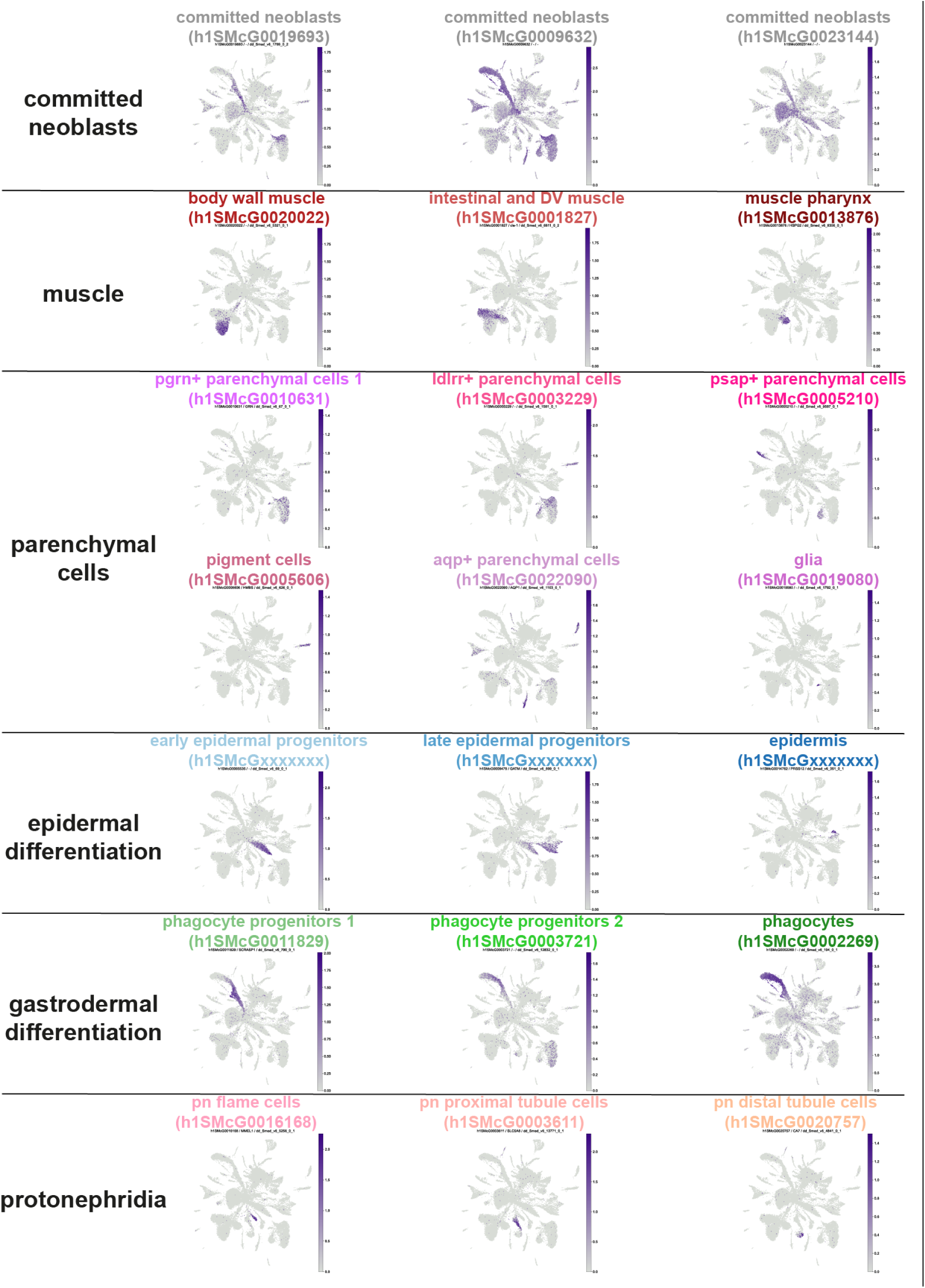
Cluster Annotation. UMAP feature plots of markers of specific cell clusters used to annotate different cell types.

**Supplementary Figure 2.**
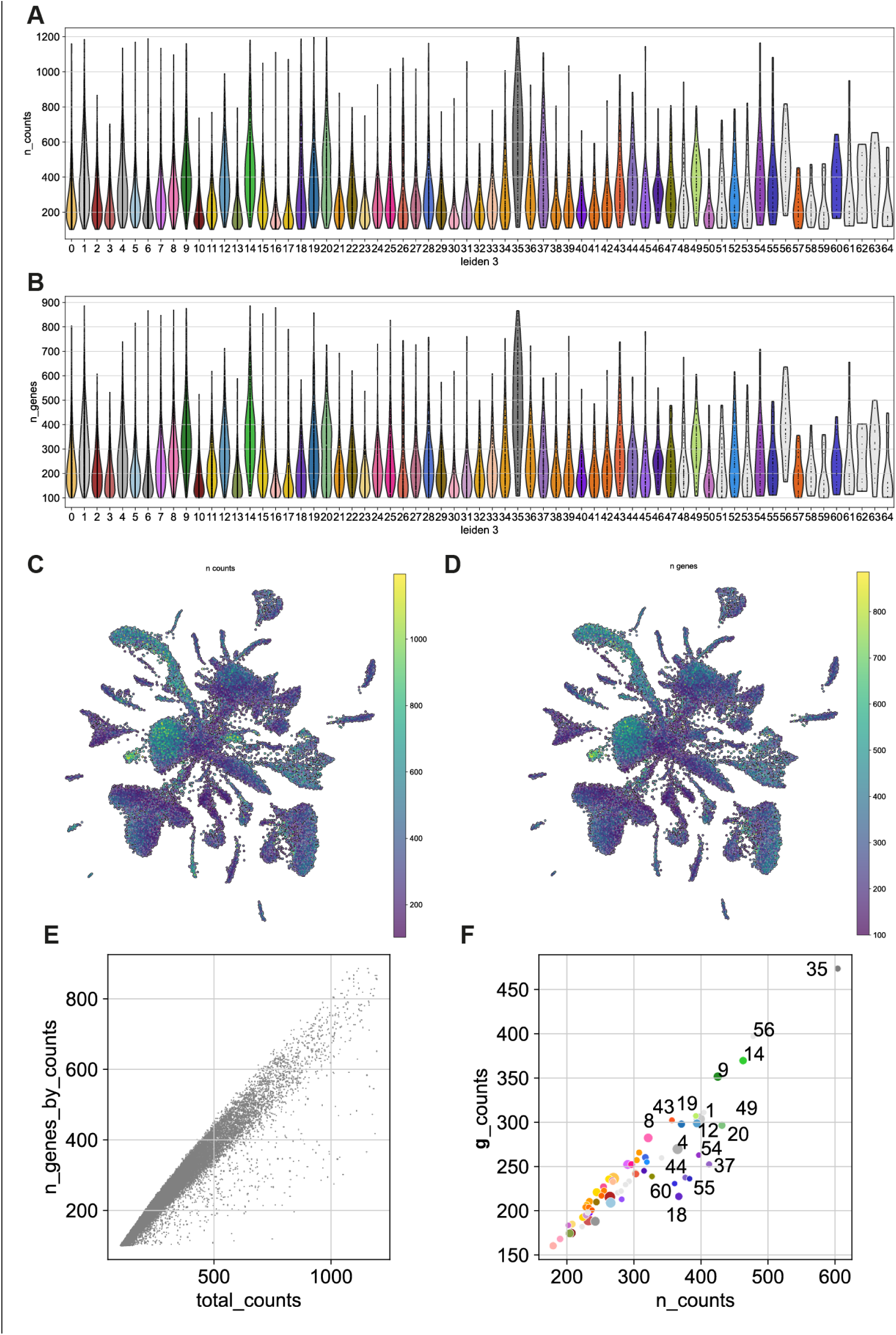
Sizes dataset metrics. A: Violin plots showing the number of UMI counts detected per cell in each cluster. B: Violin plots showing the number of genes detected per cell in each cluster. C: UMAP showing the distribution of UMI counts detected per cell. D: UMAP showing the distribution of genes detected per cell. E: Scatter plot of total number of counts vs. number of genes per cell. F: Scatter plot of average of UMI counts per cluster vs. average of genes per cluster.

**Supplementary Figure 3.**
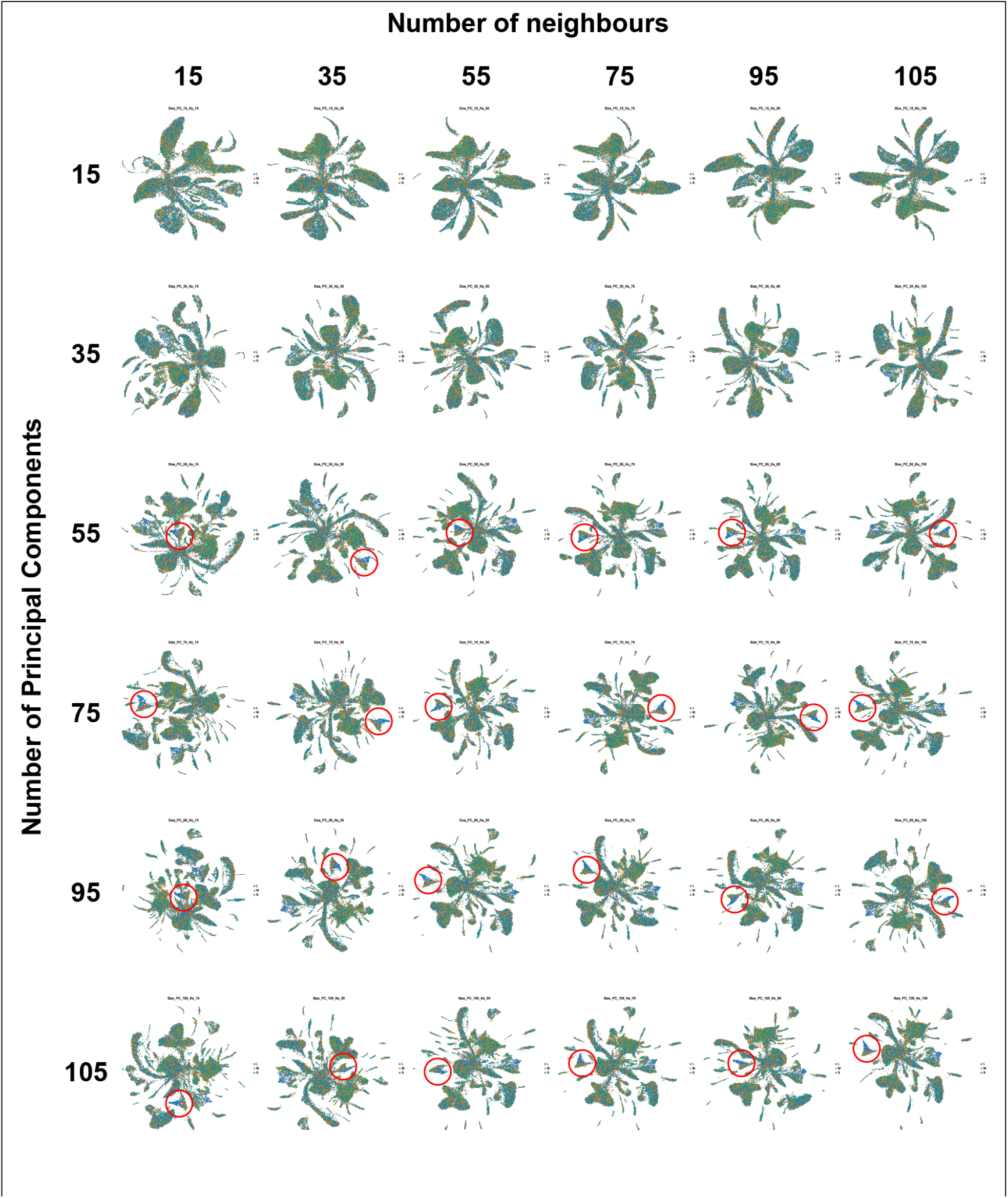
Pre-processing parameter space exploration. UMAP visualisation of the 28,738 cells pre-processed with different parameters for number of neighbours (15-105), and number of principal components (15-105). Red circle indicates the basal cells.

**Supplementary Figure 4.**
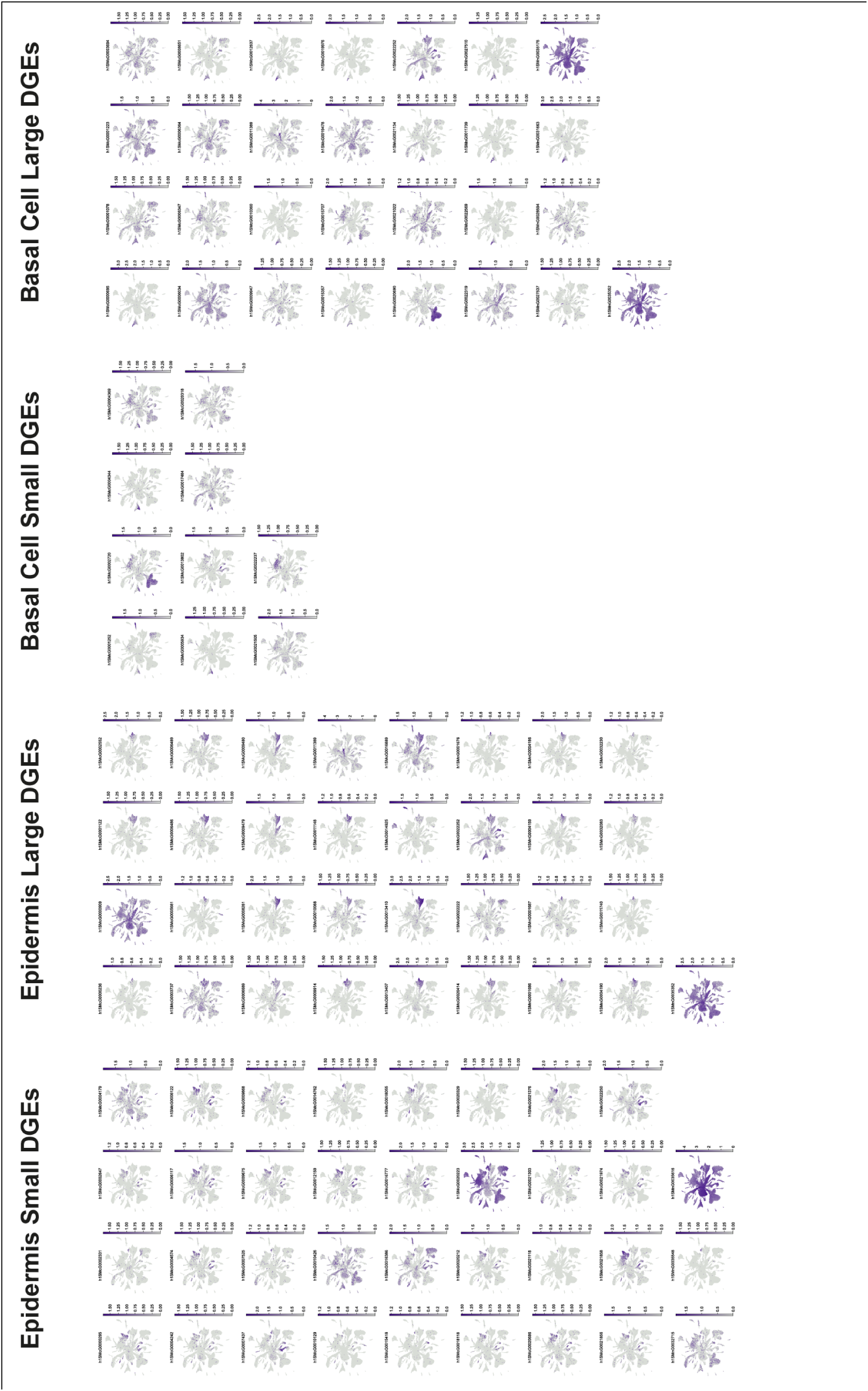
Differential gene expression analysis with DEseq2. Genes differentially regulated in S vs L planarians in the epidermis (left) and the basal cells (right).

## Supplementary Files

Supplementary File 1. **Planarian L, M and S size measurements.**

Supplementary File 2. **Leiden cluster annotation**

Supplementary File 3. ***Schmidtea mediterranea* genome annotation**

Supplementary File 4. **Cluster Wilcoxon markers**

Supplementary File 5. **Cluster Logistic Regression markers**

Supplementary File 6. **Common cluster markers UMAP visualisations**

Supplementary File 7. **Chi-squared statistical tests**

Supplementary File 8. **DEseq2 Differential Gene Expression analyses**

## Methods

### Planarian culture and maintenance

The experiments were conducted using the asexual strain derived from the clonal line Berlin-1 of *Schmidtea mediterranea*. The animals were kept at a temperature of 20°C in 1× Montjuic water, which consists of 1.6 mM NaCl, 1.0 mM CaCl_2_, 1.0 mM MgSO_4_, 0.1 mM MgCl_2_, 0.1 mM KCl, and 1.2 mM NaHCO_3_ dissolved in deionised water. The pH of the solution was adjusted to approximately 7.5 using 1 M HCl. The planarians were fed fresh cow liver every four to five days and were subjected to a minimum one-week starvation period before any experimental procedures were carried out.

### Sized animal selection and photography

The animals were manually selected and classified based on their sizes into different experimental groups through visual observational assessment. Our primary objective was to categorise the animals into four distinct body sizes, specifically designated as Extra Large (XL), Large (L), Medium (M), and Small (S). To ensure that cell numbers are comparable, we selected 10, 25, 50, and 100 animals for the XL, L, M, and S size categories respectively.

In order to obtain visual records, we took photographs, adhering to a standardised methodology of capturing groups of five animals per picture on a A4 graph millimetre paper. The process avoided animal manipulation under scope bright light to avoid inducing stress. We captured low-resolution images using a Lumens PC193 standard field camera, utilising the default settings and a 35mm focal length. We proceeded to measure the area of the individual animals to ensure the accuracy of our examination and quantify the size separation range. The images were processed and analysed using ImageJ 1.51d software, treating them as 8-bit pictures, enabling us to measure the animal’s area in pixels. Subsequently, we calculated the corresponding area in mm2 by comparing the measurements of animal pixels with the known measurements of the pixel scale present on the millimetre paper within each picture.

### ACME dissociation

The ACME solution was freshly prepared by combining commercially sourced DNase/RNase-free distilled water, methanol, glacial acetic acid, and glycerol in a ratio of 13:3:2:2 (*21*). Each sample was placed in a 15 mL Falcon tube. Montjuic water was removed using a Pasteur pipette, and approximately 100-500 μL of 7.5% N-acetyl cysteine in 1× PBS was added to cover the planarians and remove the mucus and protect the RNA. The ACME solution was promptly added to each sample, reaching a final volume of 10 mL per tube. The samples were then left to dissociate at room temperature for 45 minutes on a see-saw motion shaker at 35-45 rpm, with tubes oriented parallel to the direction of movement. We complete dissociation mechanically by pipetting the reactions up and down using 1 mL low binding pipette tips. From this point, the samples were kept on ice to prevent RNA degradation.

To remove cell aggregates and undissociated tissue fragments, each sample was filtered through a 50 μm Celltrics filter into a new 15 mL tube, and then subsequently filtered through a 40 μm Celltrics filter into another tube. The sample was centrifuged at 1000 g for 5 minutes (4 °C), discarding the supernatant except for 1-2 mL, which was used to resuspend the pellet. We filter each sample one more time using 40 μm cell strainer pipette tips (1000 μL) into a new 15-mL tube.

To clean the cells, 7 mL of buffer (1× PBS 1% BSA) was added to each sample, followed by centrifugation at 1000g for 5 minutes (4 °C). The supernatant was removed, and the pellets were resuspended in 900 μL of buffer (1× PBS 1% BSA) and transferred to 1.5-mL Eppendorf tubes. For cryopreservation, 100 μL of DMSO was added to each tube (*57*), and the samples were directly stored at -80 °C.

### SPLiT-seq

Single-cell RNA sequencing libraries were generated using a combinatorial barcoding approach, following a customised adaptation of the SPLiT-seq protocol (*22*). We developed and refined a four-day protocol, incorporating various optimizations, to achieve improved efficiency and reliability.

#### Day1: Round 1, 2, and 3 of DNA Barcoding

For cytoplasm staining, the dissociated samples were labelled with a concentration of 2μL/mL from a 1mg/mL stock solution of Alexa Fluor 488-Conjugated Concanavalin A (Invitrogen). Additionally, for nuclear staining, a concentration of 1μL/mL from a 5mM stock solution of Draq5 (eBioscience) was used. The labelling process took place in the dark at 4°C for a duration of 30-45 minutes. Subsequently, the samples were visualised using a CytoFlex S Flow Cytometer (Beckman Coulter) to determine the count of single cells per sample and measure the percentage of cells in G1 and G2 phases.

The first round of barcoding was carried out through in-cell reverse transcription (RT). Anchored poly (dT) oligos were employed to minimise the number of ribosomal reads. Each reaction utilised ∼5000 singlet events as calculated from the flow cytometry count in a volume of 8μL per well. Following this, a second and third round of barcoding were performed using ligation reactions (*21*).

#### Day2: FACS sorting and Cell Lysis

Following the barcoding process, cells were sorted using a BD FACS Aria III Cell Sorter (BD Biosciences) in 50μL of lysis buffer (comprising Tris pH 8.0 20 mM, NaCl 400 mM, EDTA pH 8.0 100 mM, and SDS 4.4%). A maximum of 25,000 events were sorted into each tube. To ensure the absence of RNase contamination, the FACS instrument was meticulously cleaned with bleach and pre-cooled before sorting. The injection and collection chambers were maintained at 4 °C throughout the sorting process. Sorting was carried out using the BD FACSDiva Software, set up in 4-Way Purity mode, with an 85μm nozzle and moderate-pressure separation (45 Psi).

Subsequently, the volume of each sorted sub-library was adjusted to 100μL, and 10μL of Proteinase K (20 mg/mL) was added to each tube. The lysates were then incubated at 55 °C for 2 hours, with manual shaking of the tubes every 15 minutes. Finally, the lysates were stored at -80 °C for further use.

#### Day3: cDNA Purification, Template Switch and dsDNA amplification

To purify the cDNA from genomic DNA, we utilised 44 μL of Dynabeads™ MyOne™ Streptavidin C1 (Invitrogen) per lysate. These magnetic beads were conjugated with streptavidin, which binds to the biotin present at the 3’-end of the third barcode. The manufacturer’s protocol for Dynabeads nucleic acid purification was strictly followed without any deviations.

Next, the cDNA was converted from single-stranded DNA (ssDNA) to double-stranded DNA (dsDNA) through a template-switch reaction. This reaction involved the following components per sample: 44 μL of 5× RT Buffer (Thermo Scientific), 44 μL of 20% Ficoll PM 400 (Sigma Aldrich), 22 μL of a mixture containing 10 mM of each of the four deoxynucleotide triphosphates (dNTPs) (NEB), 5.5 μL of TSO primer (100 μM), 11 μL of Maxima H Minus RT, and 88 μL of nuclease-free water. The samples were incubated in this reaction mix for 30 minutes at room temperature, followed by 90 minutes at 42 °C with agitation. After incubation, the template switch mix was removed using a magnetic rack.

For cDNA amplification, a total of 121 μL of 2× Kapa HiFi HotStart ReadyMix (Roche), 9.68 μL of PCR_PF (10 μM), 9.68 μL of PCR_PR (10 μM), and 101.64 μL of nuclease-free water were added to the cDNA samples. Each sample was resuspended in 220 μL of the PCR mix and divided into four PCR tubes. The following thermal cycling program was run: an initial denaturation step at 95 °C for 3 minutes, followed by five cycles of denaturation at 98 °C for 20 seconds, annealing at 65 °C for 45 seconds, and extension at 72 °C for 3 minutes. The four PCR reactions were then combined into a single 1.5 mL Eppendorf tube, and the Dynabeads were separated using a magnetic rack. Two hundred microliters of the supernatant, containing the cDNA in suspension, was divided into four wells of a qPCR plate, with 50 μL in each well and 2.5 μL of 20× EvaGreen (Biotium) per well was added. Finally, qPCR was performed until reaching the non-exponential plateau phase.

The magnetic beads were preserved for potential future use. After the template switch reaction, the beads were resuspended in 250 μL of Tris-T buffer and carefully stored at 4 °C. This storage condition ensured the stability and integrity of the beads for potential subsequent applications.

#### Day 4: Tagmentation and Round 4 of Barcoding

The qPCR reactions were subjected to SPRI size selection using Kapa Pure Beads (Roche) at ratios of 0.8× and 0.7×. This step effectively removed fragments smaller than 300bp. The concentration of the resulting libraries was determined using a Qubit fluorometer (Thermo Fisher), and the fragment distribution was assessed using an Agilent 2100 Bioanalyzer with the Agilent High Sensitivity DNA Kit.

Next, the sub-libraries underwent tagmentation using the Nextera DNA Library Preparation Kit (Illumina). The tagmentation process was promptly neutralised using the Monarch PCR & DNA Cleanup Kit (NEB). The samples were then eluted in a final volume of 20 μL using the Elution buffer.

For the fourth and final round of barcoding and library preparation, PCR amplification was performed using tagmentation master primer i7 and library-specific tagmentation primers i5. The resulting products were subjected to size selection using Kapa Pure Beads (Roche) at ratios of 0.7× and 0.6×. The fragment distribution was once again assessed using the Agilent 2100 Bioanalyzer with the Agilent High Sensitivity DNA Kit, and the library concentrations were quantified using the Qubit fluorometer.

### SPLiT-seq read processing

Following sequencing on a NovaSeq 6000 platform (Illumina) by Novogene, our generated data underwent quality control and pre-processing steps to ensure high-quality data for subsequent analyses. These steps included read alignment, barcode demultiplexing, and removal of low-quality or duplicate reads. The sequencing was performed with 150 bp length, paired-end reads. However, the reads were provided without any extensive quality verification apart from a basic chastity check. Therefore, we conducted initial quality checks using the FastQC software. To further improve the quality of the data, we utilized the CutAdapt v2.1 tool. For read 1, we removed Illumina universal adaptors, as well as short and low-quality reads, using the command: “cutadapt -b AGATCGGAAGAG -m 60 -j 4”. Similarly, for read 2, we removed short and low-quality reads, terminal Ns, and the Nextera adapter sequence using the command: “cutadapt -m 94 -j 4 --trim-n -b CTGTCTCTTATA”. To ensure that only paired reads were retained, we employed the Makepairs tool in our preprocessing pipeline. This step helped us maintain the integrity of paired-end reads for subsequent analyses.

We used a new *Schmidtea mediterranea* genome and annotation by the Jochen Rink lab (manuscript in preparation). After the preprocessing steps, we utilized the Drop-seq_tools-2.3.0 software package (available at https://github.com/broadinstitute/Drop-seq) to create sequence dictionary, refFlat, reduced GTF, and interval files. To generate the STAR-2.7.3a index (*58*), we used the following settings: --sjdbOverhang 99, --genomeSAindexNbases 13, and --genomeChrBinNbits 14. This index allows for efficient mapping of the reads to the reference genome. Each of the two sub-libraries sequenced in our study was processed separately. For barcode extraction, verification, and correction (with hamming distance ≤1), we utilized the SPLiTseq toolbox (available at https://github.com/RebekkaWegmann/splitseq_toolbox). This toolbox incorporates many of the components of Drop-seq_tools-2.3.0. Mapping of the processed reads was performed using STAR-2.7.3a with the --quantMode GeneCounts option and default settings. To re-order and merge the aligned and tagged reads, we employed Picard v2.21.1-SNAPSHOT (developed by the Broad Institute, available at http://broadinstitute.github.io/picard/). Specifically, we used the SortSam and MergeBamAlignment tools. For further annotation of the mapped reads, we used the Drop-seq_tools-2.3.0 TagReadWithInterval and TagReadWithGeneFunction tools in sequential order. These tools utilize the custom refFlat and genes.intervals files created earlier to note the mapping location and assign gene function to the reads. The resulting mapping files were then used to create expression matrices for each library individually using the Drop-seq_tools-2.3.0 DigitalExpression tool. We applied the following settings: READ_MQ=0, EDIT_DISTANCE=1, MIN_NUM_GENES_PER_CELL=50, and LOCUS_FUNCTION_LIST=INTRONIC. These settings ensured the generation of accurate expression matrices for subsequent analyses.

### Genome annotation

The *S. mediterranea* genome and annotation were subjected to a snakemake workflow of standard tools for gene annotation, found at https://github.com/apposada/gene_annot. Briefly, the longest isoforms per gene were extracted using gffread on an AGAT-standardised version of the GFF3 annotation file and the genome sequence FASTA. These sequences were translated into proteins using TransDecoder (https://github.com/TransDecoder/TransDecoder) using evidence from BLAST reciprocal best hits against the UniProt database and hmmer queried against the PFAM database, as previously described (*37*). Protein sequences were further annotated using (i) eggNOG against the protein set of all metazoa (*59*), (ii) InterProScan against the PFAM, SFAM, and Panther databases, (iii) BLAST reciprocal best hits against the UniProt database, and (iv) assigning co-orthologues from a set of model organism species using OrthoFinder (*60*). Evidence from (ii, iii, iv) was used to annotate transcription factors into TF classes using also data from AnimalTFDB 3.0 (*61*), as previously described (*37*).

### Single cell transcriptomic analysis

The scRNA-seq data was analysed using Scanpy. The gene expression data from each of the two sub-libraries, encapsulated in the 10x matrices, were uploaded and processed to associate the sample names with the Anndata object. We tagged each cell observation in adata.obs with its sample of origin. Additionally, each gene in the adata.var was tagged with the previously described gene annotations.

We applied additional quality control (QC) measures to the scRNA-seq data in order to eliminate low-quality cells. Specifically, we used the command “sc.pp.filter_cells(adata, min_counts=100)” to filter out cells with a low number of UMIs (Unique Molecular Identifiers), and “sc.pp.filter_cells(adata, min_genes=100)” to filter out cells with a low expression of genes. To further assess the quality of the data, QC metrics were calculated using the command “sc.pp.calculate_qc_metrics(adata, qc_vars=[’mt’], percent_top=None, log1p=False, inplace=True)”. Cells with a low number of expressed genes are often indicative of poor quality or technical artifacts, and therefore, they were excluded from further analysis. Similarly, cells with a low total UMI count may be associated with low RNA content or technical issues, so they were also excluded to improve the downstream analysis quality. To achieve this, we filtered the matrix in the Anndata object, removing cells where the number of expressed genes was less than 900 using the command “adata = adata[adata.obs.n_genes_by_counts < 900, :]“, and cells where the total molecule count was less than 1200 using the command “adata = adata[adata.obs.total_counts < 1200, :]”. Furthermore, we performed normalization to ensure consistent read counts across cells, which was accomplished using the command “sc.pp.normalize_total(adata)”. Subsequently, the data was log-transformed to stabilize the variances, and variable genes were identified to pinpoint genes that exhibited significant expression variation across cells. For this purpose, the top 24,000 variable genes were selected using the command “sc.pp.highly_variable_genes(adata, n_top_genes=24000)”.

To reduce the dimensionality of the dataset, we used PCA (Principal Component Analysis) and UMAP (Uniform Manifold Approximation and Projection for Dimension Reduction). PCA was performed on the first 150 principal components using the command “sc.tl.pca(adata, svd_solver=’arpack’, n_comps=150)”. An elbow plot was created to assess the amount of variance captured by these components. Based on the PCA results, the neighbourhood relationships between cells were determined using the first 95 principal components with the command “sc.pp.neighbors(adata, n_neighbors=55, n_pcs=95)”. For UMAP visualisation, the calculated neighbours were utilised, and parameters such as minimum distance and spread were specified to control the density and layout of the visualisation. This was accomplished using the command “sc.tl.umap(adata, min_dist=0.75, spread=1.25, alpha=1, gamma=1.0)”. Clustering was performed using the Leiden algorithm, and the resolution was set to values 1, 2, 3, and 4 to determine the optimal granularity of the clusters. To evaluate the consistency and robustness of the clustering results, correlation matrices and dendrograms were examined for each clustering resolution. Reliable clustering was indicated by consistent and well-separated clusters in both the correlation matrix and dendrogram. Additionally, known marker genes were referenced to assess whether the clustering algorithm accurately captured the expected biology (*30*). Finally, to identify marker genes associated with each cluster, gene rankings were generated using both the ’wilcoxon’ and ’logreg’ methods. The cluster dendrogram was obtained with the scanpy function “sc.pl.dendrogram”.

To gain further insights into the cellular composition and heterogeneity present in the scRNA-seq data, we visualised cluster-specific marker genes using dot plots, heat maps and features plots. These visualisations provided a concise representation of gene expression within each cluster as well as across clusters. By performing these analyses and examining the expression patterns of known marker genes, we were able to identify distinct cell populations and infer their respective cell types. To obtain log2 ratios between cell numbers in large and small planarians we divided the number of cells in each cluster in each size category by the sum of the cells in the category. We then applied logarithms and obtained the ratios by subtracting the log frequency values. These values were plotted on the UMAP using a custom colour scale, since most values are between -1.3 and 1.1 except cluster 54 (secretory 4) which has an outlier value of 4.4.

### Pseudo bulk and differential gene expression analysis

Using the cell annotation derived from our single cell analysis (i.e. clustering, broad cell types, etc.) as well as cell information such as size of the organism of origin (L,M,S) and cell library, we generated a pseudo bulk matrix by aggregating, for a given gene X, all the counts of gene X from cells coming from the same cluster and library, effectively generating a pseudo bulk matrix with genes in rows and ‘pseudosamples’ (i.e. all combinations of cell type + ‘condition’ i.e. organism size + ‘replicate’ i.e. library) in columns. For a given cell type i, we subsetted this pseudo bulk matrix to keep ‘pseudosamples’ from cell type i as well as genes annotated as “high-confidence” in the gene annotation (see Methods), and performed differential gene expression analysis using DESeq2 (*52*) inside a custom wrapper. Briefly, we filtered genes with less than two counts in at least one replicate and performed a negative binomial test from DESeq2 with contrast “L” over “S”. Genes were considered differentially expressed if the p-value of the test was lower than 0.05.

### In situ hybridization images

All in situ hybridization images were downloaded from https://digiworm.wi.mit.edu/ (*28*). The genes selected are dd_Smed_v4_21541_0_1 (h1SMcG0010053), dd_Smed_v4_750_0_1 (h1SMcG0017679), dd_Smed_v4_43_0_1, (h1SMcG0005326) and dd_Smed_v4_7593_0_1 (h1SMcG0011854).

### Statistical analysis

To assess the relationship between cluster memberships and cell sizes, Pearson’s chi-squared tests for independence were conducted. Each omnibus chi-squared test was followed by a post hoc analysis of the standardised residuals to assess which category combinations were significantly over- or under-represented. Multiple testing was corrected using the Benjamini-Hochberg correction, which was deemed more appropriate than the more commonly used Bonferroni correction as it focused on reducing false discovery rate under multiple testing.

All statistical analysis was conducted in R version 4.3.0 (R Core Team, 2023 - https://www.R-project.org/), using RStudio 2023.03.1+446 (Posit team, 2023 - http://www.posit.co/). Additionally to baseR functionalities, the package suite tidyverse version 2.0.0 (*62*), and packages chisq.posthoc.test version 0.1.2 (Ebbert, 2019 - https://CRAN.R-project.org/package=chisq.posthoc.test), colorspace version 2.1-0 (*63*), and rstatix version 0.7.2 (Kassambara, 2023 - https://CRAN.R-project.org/package=rstatix) were used for analyses and development of heatmaps

## References

1. S. Nakagawa, F. Kar, R. E. O’Dea, J. L. Pick, M. Lagisz, Divide and conquer? Size adjustment with allometry and intermediate outcomes. BMC Biol 15, 107 (2017).

2. M. Kleiber, Body size and metabolism. Hilgardia 6, 315–353 (1932).

3. H. Zeng, What is a cell type and how to define it? Cell 185, 2739–2755 (2022).

4. S. A. Morris, The evolving concept of cell identity in the single cell era. Development 146, (2019).

5. J. S. Fleck, J. G. Camp, B. Treutlein, What is a cell type? Science 381, 733–734 (2023).

6. What Is Your Conceptual Definition of “Cell Type” in the Context of a Mature Organism? Cell Syst 4, 255–259 (2017).

7. C. Trapnell, Defining cell types and states with single-cell genomics. Genome Res 25, 1491–1498 (2015).

8. B. Xia, I. Yanai, A periodic table of cell types. Development 146, (2019).

9. S. Domcke, J. Shendure, A reference cell tree will serve science better than a reference cell atlas. Cell 186, 1103–1114 (2023).

10. M. Mircea, S. Semrau, How a cell decides its own fate: a single-cell view of molecular mechanisms and dynamics of cell-type specification. Biochem Soc Trans 49, 2509–2525 (2021).

11. N. J. Oviedo, P. A. Newmark, A. Sanchez Alvarado, Allometric scaling and proportion regulation in the freshwater planarian Schmidtea mediterranea. Dev Dyn 226, 326–333 (2003).

12. L. Heumos, A. C. Schaar, C. Lance, A. Litinetskaya, F. Drost, L. Zappia, M. D. Lucken, D. C. Strobl, J. Henao, F. Curion, C. Single-cell Best Practices, H. B. Schiller, F. J. Theis, Best practices for single-cell analysis across modalities. Nat Rev Genet 24, 550–572 (2023).

13. T. Stuart, R. Satija, Integrative single-cell analysis. Nat Rev Genet 20, 257–272 (2019).

14. A. A. Khozyainova, A. A. Valyaeva, M. S. Arbatsky, S. V. Isaev, P. S. Iamshchikov, E. V. Volchkov, M. S. Sabirov, V. R. Zainullina, V. I. Chechekhin, R. S. Vorobev, M. E. Menyailo, P. A. Tyurin-Kuzmin, E. V. Denisov, Complex Analysis of Single-Cell RNA Sequencing Data. Biochemistry (Mosc) 88, 231–252 (2023).

15. P. Sant, K. Rippe, J. P. Mallm, Approaches for single-cell RNA sequencing across tissues and cell types. Transcription, 1–19 (2023).

16. S. Wang, S. T. Sun, X. Y. Zhang, H. R. Ding, Y. Yuan, J. J. He, M. S. Wang, B. Yang, Y. B. Li, The Evolution of Single-Cell RNA Sequencing Technology and Application: Progress and Perspectives. Int J Mol Sci 24, (2023).

17. S. Aldridge, S. A. Teichmann, Single cell transcriptomics comes of age. Nat Commun 11, 4307 (2020).

18. E. Denisenko, B. B. Guo, M. Jones, R. Hou, L. de Kock, T. Lassmann, D. Poppe, O. Clement, R. K. Simmons, R. Lister, A. R. R. Forrest, Systematic assessment of tissue dissociation and storage biases in single-cell and single-nucleus RNA-seq workflows. Genome Biol 21, 130 (2020).

19. R. Massoni-Badosa, G. Iacono, C. Moutinho, M. Kulis, N. Palau, D. Marchese, J. Rodriguez-Ubreva, E. Ballestar, G. Rodriguez-Esteban, S. Marsal, M. Aymerich, D. Colomer, E. Campo, A. Julia, J. I. Martin-Subero, H. Heyn, Sampling time-dependent artifacts in single-cell genomics studies. Genome Biol 21, 112 (2020).

20. S. C. van den Brink, F. Sage, A. Vertesy, B. Spanjaard, J. Peterson-Maduro, C. S. Baron, C. Robin, A. van Oudenaarden, Single-cell sequencing reveals dissociation-induced gene expression in tissue subpopulations. Nat Methods 14, 935–936 (2017).

21. H. Garcia-Castro, N. J. Kenny, M. Iglesias, P. Alvarez-Campos, V. Mason, A. Elek, A. Schonauer, V. A. Sleight, J. Neiro, A. Aboobaker, J. Permanyer, M. Irimia, A. Sebe-Pedros, J. Solana, ACME dissociation: a versatile cell fixation-dissociation method for single-cell transcriptomics. Genome Biol 22, 89 (2021).

22. A. B. Rosenberg, C. M. Roco, R. A. Muscat, A. Kuchina, P. Sample, Z. Yao, L. T. Graybuck, D. J. Peeler, S. Mukherjee, W. Chen, S. H. Pun, D. L. Sellers, B. Tasic, G. Seelig, Single-cell profiling of the developing mouse brain and spinal cord with split-pool barcoding. Science 360, 176–182 (2018).

23. J. Baguñà, R. Romero, E. Saló, J. Collet, C. Auladell, M. Ribas, M. Riutort, J. García-Fernàndez, F. Burgaya, D. Bueno, “Growth, Degrowth and Regeneration as Developmental Phenomena in Adult Freshwater Planarians” in Experimental Embryology in Aquatic Plants and Animals, H.-J. Marthy, Ed. (Springer US, Boston, MA, 1990), pp. 129–162.

24. M. Ivankovic, R. Haneckova, A. Thommen, M. A. Grohme, M. Vila-Farre, S. Werner, J. C. Rink, Model systems for regeneration: planarians. Development 146, (2019).

25. P. W. Reddien, The Cellular and Molecular Basis for Planarian Regeneration. Cell 175, 327–345 (2018).

26. J. Baguñà, R. Romero, Quantitative analysis of cell types during growth, degrowth and regeneration in the planarians Dugesia mediterranea and Dugesia tigrina. Hydrobiologia 84, 181–194 (1981).

27. A. Thommen, S. Werner, O. Frank, J. Philipp, O. Knittelfelder, Y. Quek, K. Fahmy, A. Shevchenko, B. M. Friedrich, F. Julicher, J. C. Rink, Body size-dependent energy storage causes Kleiber’s law scaling of the metabolic rate in planarians. Elife 8, (2019).

28. C. T. Fincher, O. Wurtzel, T. de Hoog, K. M. Kravarik, P. W. Reddien, Cell type transcriptome atlas for the planarian Schmidtea mediterranea. Science 360, (2018).

29. H. Garcia-Castro, J. Solana, Single-cell transcriptomics in planaria: new tools allow new insights into cellular and evolutionary features. Biochem Soc Trans 50, 1237–1246 (2022).

30. M. Plass, J. Solana, F. A. Wolf, S. Ayoub, A. Misios, P. Glazar, B. Obermayer, F. J. Theis, C. Kocks, N. Rajewsky, Cell type atlas and lineage tree of a whole complex animal by single-cell transcriptomics. Science 360, eaaq1723 (2018).

31. J. C. van Wolfswinkel, D. E. Wagner, P. W. Reddien, Single-cell analysis reveals functionally distinct classes within the planarian stem cell compartment. Cell Stem Cell 15, 326–339 (2014).

32. O. Wurtzel, L. E. Cote, A. Poirier, R. Satija, A. Regev, P. W. Reddien, A Generic and Cell-Type-Specific Wound Response Precedes Regeneration in Planarians. Dev Cell 35, 632–645 (2015).

33. O. Wurtzel, I. M. Oderberg, P. W. Reddien, Planarian Epidermal Stem Cells Respond to Positional Cues to Promote Cell-Type Diversity. Dev Cell 40, 491–504 e495 (2017).

34. B. W. Benham-Pyle, C. E. Brewster, A. M. Kent, F. G. Mann, Jr., S. Chen, A. R. Scott, A. C. Box, A. Sanchez Alvarado, Identification of rare, transient post-mitotic cell states that are induced by injury and required for whole-body regeneration in Schmidtea mediterranea. Nat Cell Biol 23, 939–952 (2021).

35. L. S. Swapna, A. M. Molinaro, N. Lindsay-Mosher, B. J. Pearson, J. Parkinson, Comparative transcriptomic analyses and single-cell RNA sequencing of the freshwater planarian Schmidtea mediterranea identify major cell types and pathway conservation. Genome Biol 19, 124 (2018).

36. J. Schindelin, I. Arganda-Carreras, E. Frise, V. Kaynig, M. Longair, T. Pietzsch, S. Preibisch, C. Rueden, S. Saalfeld, B. Schmid, J. Y. Tinevez, D. J. White, V. Hartenstein, K. Eliceiri, P. Tomancak, A. Cardona, Fiji: an open-source platform for biological-image analysis. Nat Methods 9, 676–682 (2012).

37. P. Alvarez-Campos, H. Garcia-Castro, E. Emili, A. Perez-Posada, D. A. Salamanca-Diaz, V. Mason, B. Metzger, A. E. Bely, N. Kenny, B. D. Ozpolat, J. Solana, Annelid adult cell type diversity and their pluripotent cellular origins. bioRxiv, (2023).

38. T. Chari, L. Pachter, The specious art of single-cell genomics. PLoS Comput Biol 19, e1011288 (2023).

39. M. L. Scimone, O. Wurtzel, K. Malecek, C. T. Fincher, I. M. Oderberg, K. M. Kravarik, P. W. Reddien, foxF-1 Controls Specification of Non-body Wall Muscle and Phagocytic Cells in Planarians. Curr Biol 28, 3787–3801 e3786 (2018).

40. X. He, N. Lindsay-Mosher, Y. Li, A. M. Molinaro, J. Pellettieri, B. J. Pearson, FOX and ETS family transcription factors regulate the pigment cell lineage in planarians. Development 144, 4540–4551 (2017).

41. B. M. Stubenhaus, J. P. Dustin, E. R. Neverett, M. S. Beaudry, L. E. Nadeau, E. Burk-McCoy, X. He, B. J. Pearson, J. Pellettieri, Light-induced depigmentation in planarians models the pathophysiology of acute porphyrias. Elife 5, (2016).

42. C. Wang, X. S. Han, F. F. Li, S. Huang, Y. W. Qin, X. X. Zhao, Q. Jing, Forkhead containing transcription factor Albino controls tetrapyrrole-based body pigmentation in planarian. Cell Discov 2, 16029 (2016).

43. J. C. Rink, H. T. Vu, A. Sanchez Alvarado, The maintenance and regeneration of the planarian excretory system are regulated by EGFR signaling. Development 138, 3769–3780 (2011).

44. H. Thi-Kim Vu, J. C. Rink, S. A. McKinney, M. McClain, N. Lakshmanaperumal, R. Alexander, A. Sanchez Alvarado, Stem cells and fluid flow drive cyst formation in an invertebrate excretory organ. Elife 4, (2015).

45. M. L. Scimone, M. Srivastava, G. W. Bell, P. W. Reddien, A regulatory program for excretory system regeneration in planarians. Development 138, 4387–4398 (2011).

46. H. T. Vu, S. Mansour, M. Kucken, C. Blasse, C. Basquin, J. Azimzadeh, E. W. Myers, L. Brusch, J. C. Rink, Dynamic Polarization of the Multiciliated Planarian Epidermis between Body Plan Landmarks. Dev Cell 51, 526–542 e526 (2019).

47. D. J. Forsthoefel, N. I. Cejda, U. W. Khan, P. A. Newmark, Cell-type diversity and regionalized gene expression in the planarian intestine. Elife 9, (2020).

48. D. J. Forsthoefel, A. E. Park, P. A. Newmark, Stem cell-based growth, regeneration, and remodeling of the planarian intestine. Dev Biol 356, 445–459 (2011).

49. D. J. Forsthoefel, F. A. Waters, P. A. Newmark, Generation of cell type-specific monoclonal antibodies for the planarian and optimization of sample processing for immunolabeling. BMC Dev Biol 14, 45 (2014).

50. R. H. Roberts-Galbraith, J. L. Brubacher, P. A. Newmark, A functional genomics screen in planarians reveals regulators of whole-brain regeneration. Elife 5, (2016).

51. I. E. Wang, S. W. Lapan, M. L. Scimone, T. R. Clandinin, P. W. Reddien, Hedgehog signaling regulates gene expression in planarian glia. Elife 5, (2016).

52. M. I. Love, W. Huber, S. Anders, Moderated estimation of fold change and dispersion for RNA-seq data with DESeq2. Genome Biol 15, 550 (2014).

53. M. D. Luecken, F. J. Theis, Current best practices in single-cell RNA-seq analysis: a tutorial. Mol Syst Biol 15, e8746 (2019).

54. J. W. Squair, M. Gautier, C. Kathe, M. A. Anderson, N. D. James, T. H. Hutson, R. Hudelle, T. Qaiser, K. J. E. Matson, Q. Barraud, A. J. Levine, G. La Manno, M. A. Skinnider, G. Courtine, Confronting false discoveries in single-cell differential expression. Nat Commun 12, 5692 (2021).

55. C. Hafemeister, F. Halbritter, Single-cell RNA-seq differential expression tests within a sample should use pseudo-bulk data of pseudo-replicates. bioRxiv, 2023.2003.2028.534443 (2023).

56. G. T. Eisenhoffer, H. Kang, A. Sanchez Alvarado, Molecular analysis of stem cells and their descendants during cell turnover and regeneration in the planarian Schmidtea mediterranea. Cell Stem Cell 3, 327–339 (2008).

57. A. Guillaumet-Adkins, G. Rodriguez-Esteban, E. Mereu, M. Mendez-Lago, D. A. Jaitin, A. Villanueva, A. Vidal, A. Martinez-Marti, E. Felip, A. Vivancos, H. Keren-Shaul, S. Heath, M. Gut, I. Amit, I. Gut, H. Heyn, Single-cell transcriptome conservation in cryopreserved cells and tissues. Genome Biol 18, 45 (2017).

58. A. Dobin, C. A. Davis, F. Schlesinger, J. Drenkow, C. Zaleski, S. Jha, P. Batut, M. Chaisson, T. R. Gingeras, STAR: ultrafast universal RNA-seq aligner. Bioinformatics 29, 15–21 (2013).

59. C. P. Cantalapiedra, A. Hernandez-Plaza, I. Letunic, P. Bork, J. Huerta-Cepas, eggNOG-mapper v2: Functional Annotation, Orthology Assignments, and Domain Prediction at the Metagenomic Scale. Mol Biol Evol 38, 5825–5829 (2021).

60. D. M. Emms, S. Kelly, OrthoFinder: phylogenetic orthology inference for comparative genomics. Genome Biol 20, 238 (2019).

61. H. Hu, Y. R. Miao, L. H. Jia, Q. Y. Yu, Q. Zhang, A. Y. Guo, AnimalTFDB 3.0: a comprehensive resource for annotation and prediction of animal transcription factors. Nucleic Acids Res 47, D33–D38 (2019).

62. W. Hadley, A. Mara, B. Jennifer, C. Winston, M. Lucy D’Agostino, F. Romain, G. Garrett, H. Alex, H. Lionel, H. Jim, K. Max, P. Thomas Lin, M. Evan, B. Stephan Milton, M. Kirill, O. Jeroen, R. David, S. Dana Paige, S. Vitalie, T. Kohske, V. Davis, W. Claus, W. Kara, Y. Hiroaki, Welcome to the Tidyverse. Journal of Open Source Software 4, 1686 (2019).

63. A. Zeileis, J. C. Fisher, K. Hornik, R. Ihaka, C. D. McWhite, P. Murrell, R. Stauffer, C. O. Wilke, colorspace: A Toolbox for Manipulating and Assessing Colors and Palettes. Journal of Statistical Software 96, 1–49 (2020).

