## Supplementary File 4 for "Allometry of cell types in planarians by single cell transcriptomics"

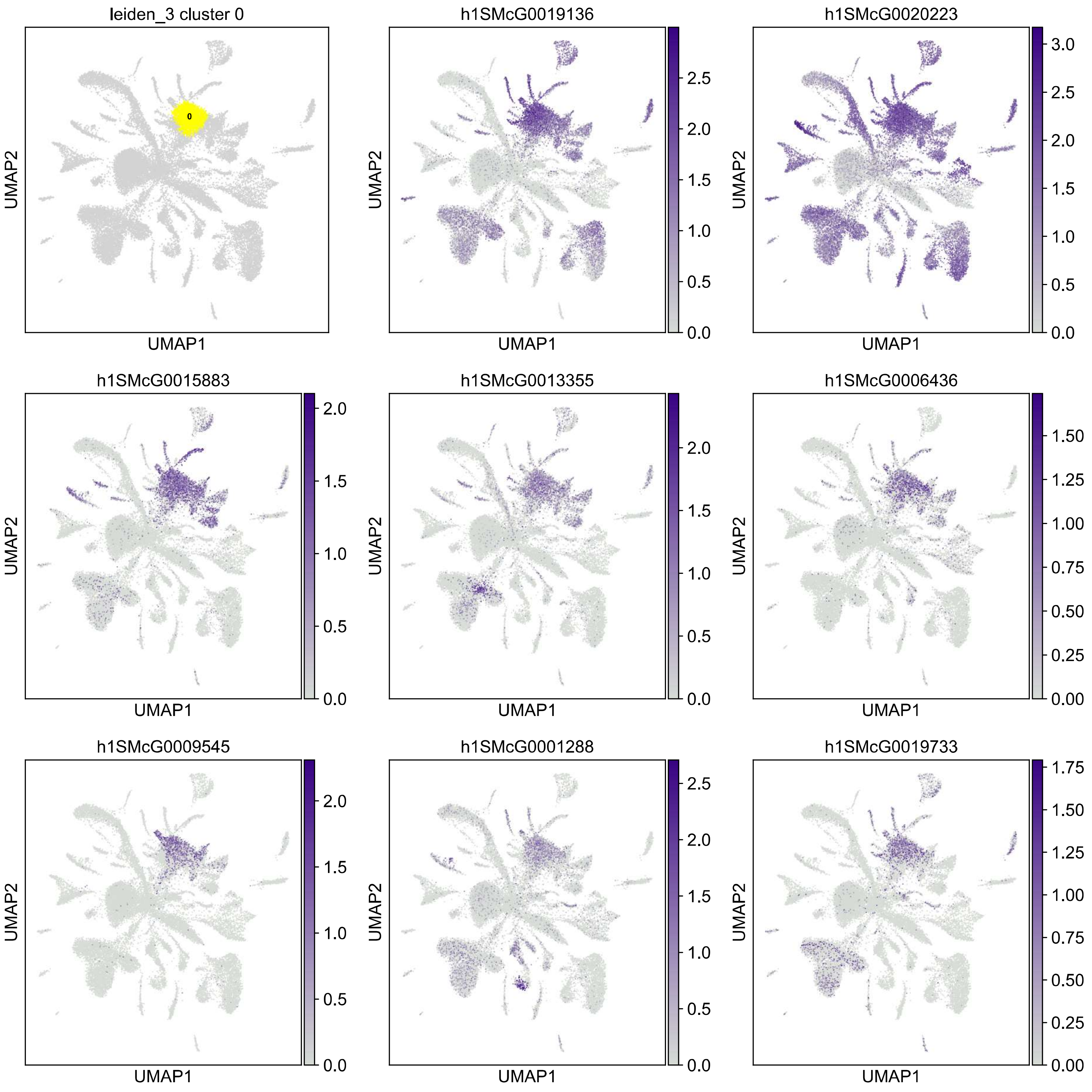
leiden_3 cluster 0
0
h1SMcG0019136
h1SMcG0020223
h1SMcG0015883
h1SMcG0013355
h1SMcG0006436
h1SMcG0009545
h1SMcG0001288
h1SMcG0019733
UMAP1
UMAP2

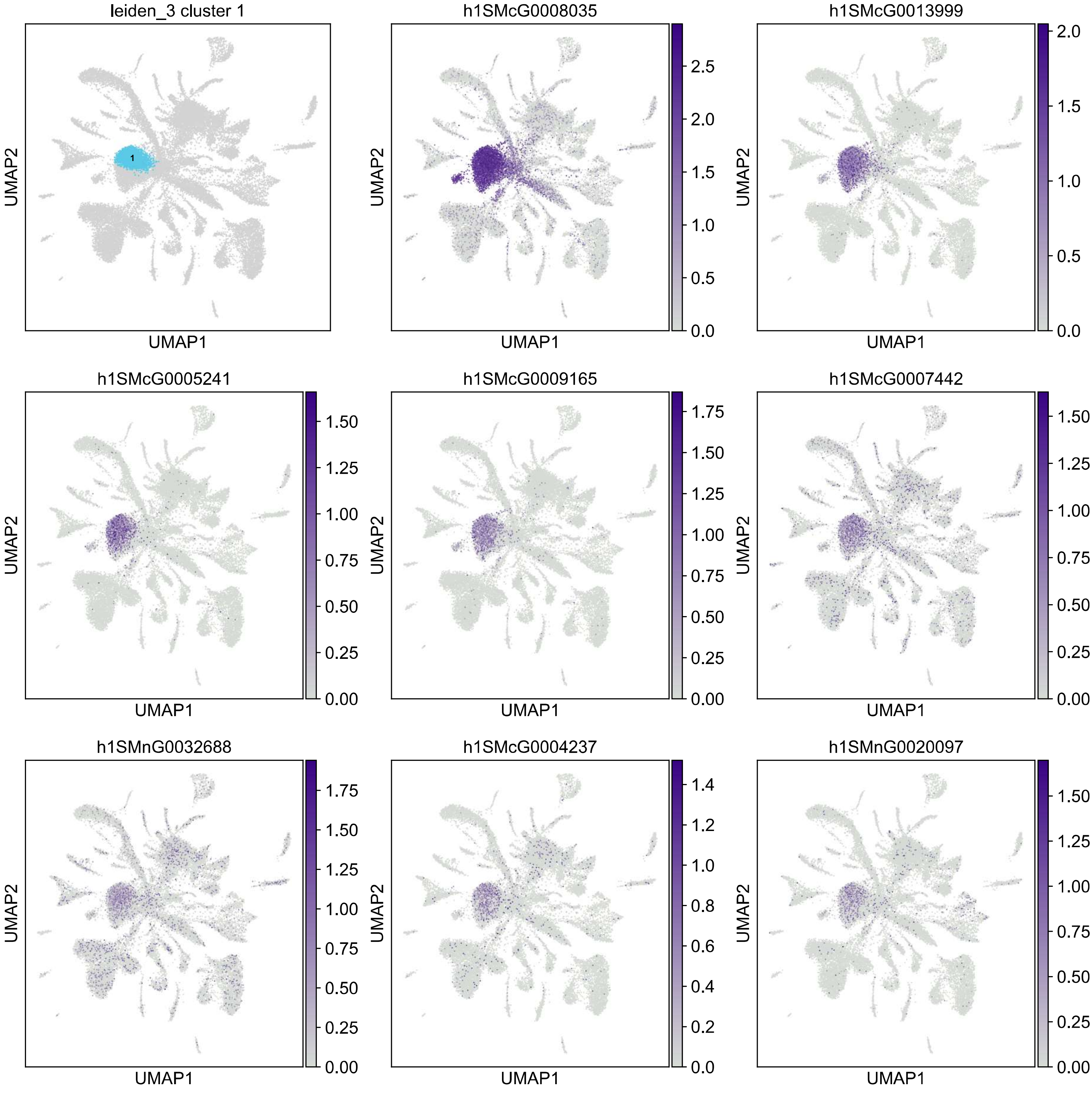

leiden_3 cluster 1
1
h1SMcG0008035
h1SMcG0013999
h1SMcG0005241
h1SMcG0009165
h1SMcG0007442
h1SMnG0032688
h1SMcG0004237
h1SMnG0020097
UMAP1
UMAP2

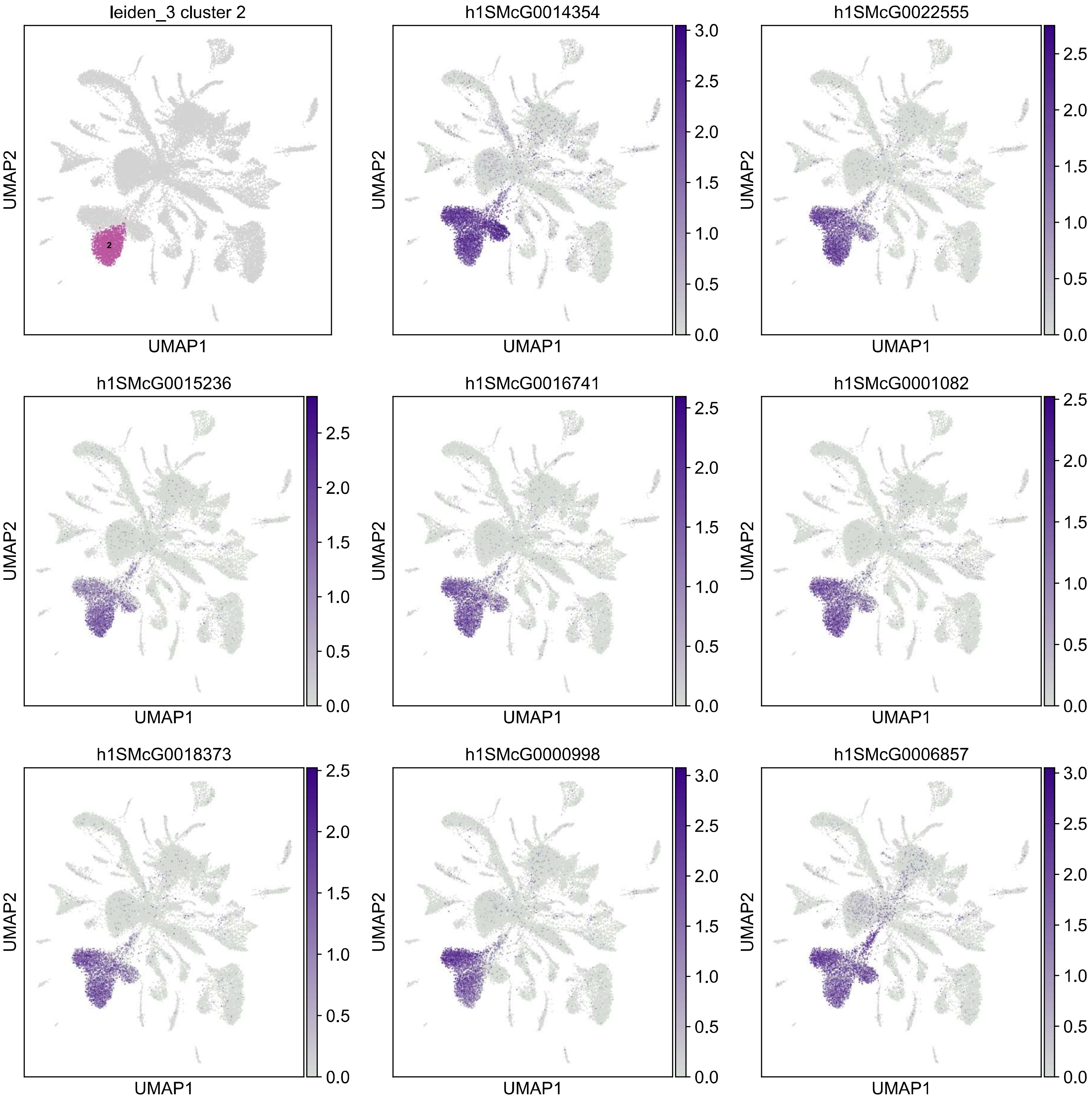
leiden_3 cluster 2
2
h1SMcG0014354
h1SMcG0022555
h1SMcG0015236
h1SMcG0016741
h1SMcG0001082
h1SMcG0018373
h1SMcG0000998
h1SMcG0006857
UMAP1
UMAP2
0.0
0.5
1.0
1.5
2.0
2.5
3.0

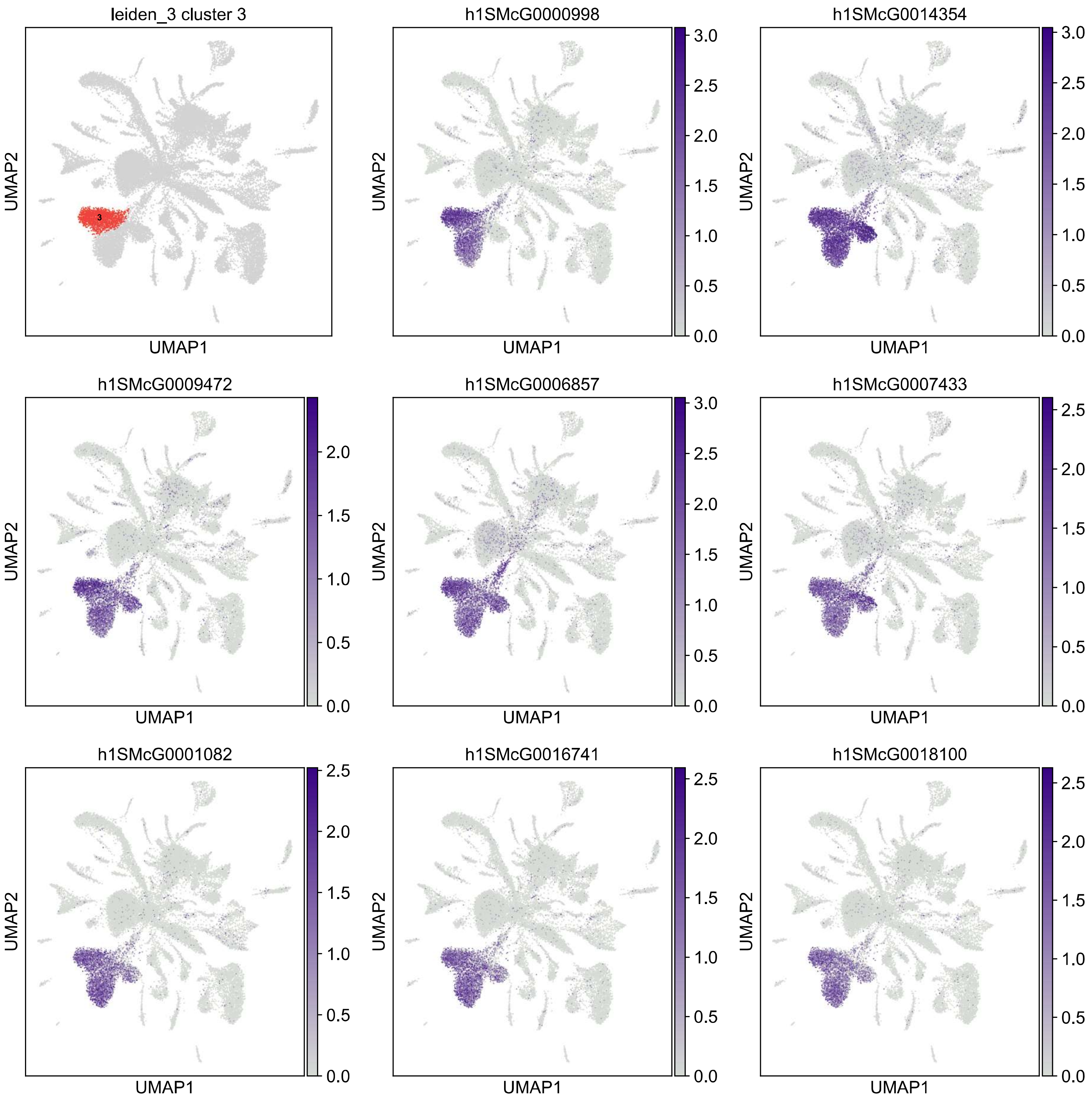
leiden_3 cluster 3
h1SMcG0000998
h1SMcG0014354
h1SMcG0009472
h1SMcG0006857
h1SMcG0007433
h1SMcG0001082
h1SMcG0016741
h1SMcG0018100
UMAP1
UMAP2

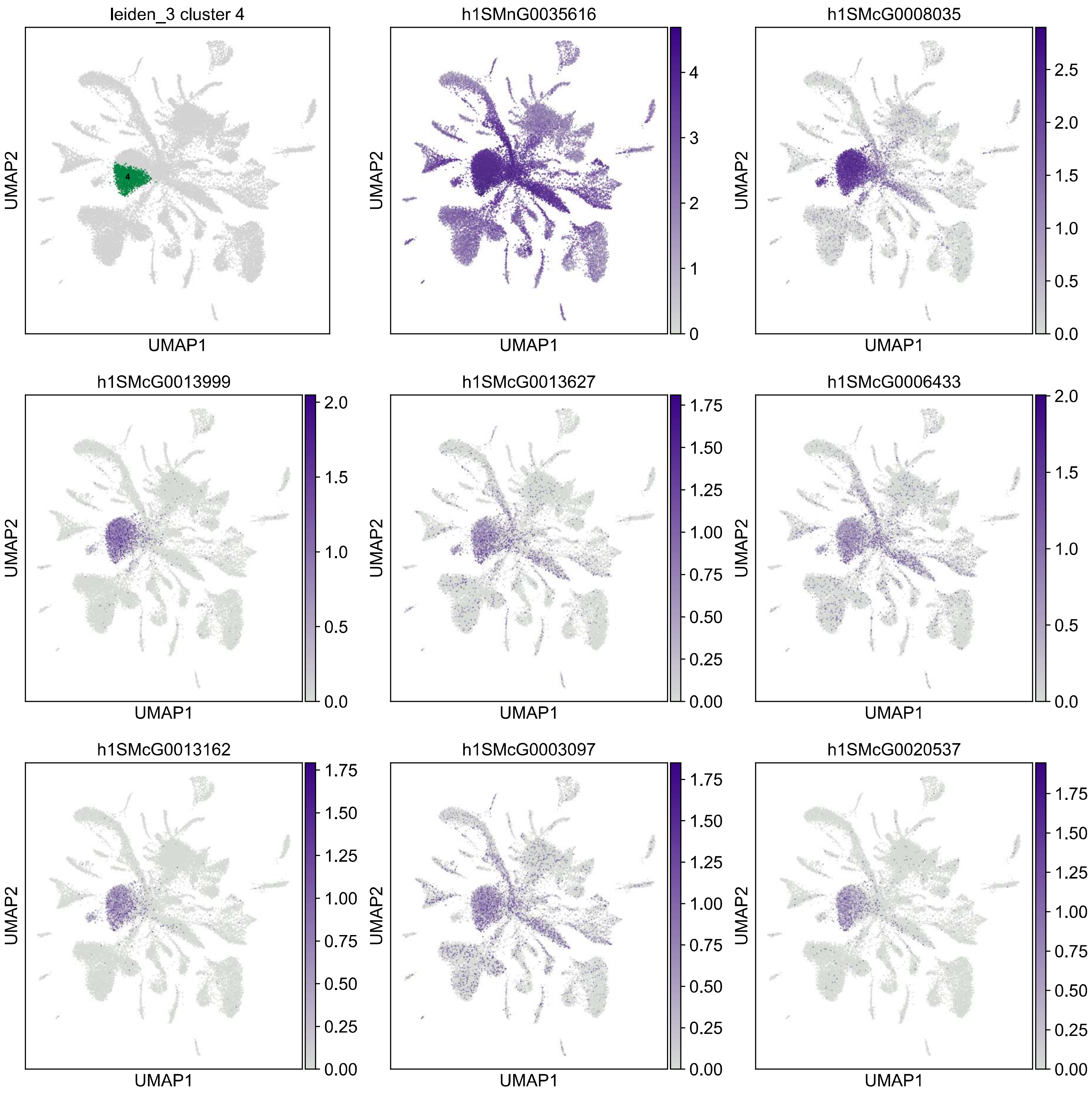

leiden_3 cluster 4
h1SMnG0035616
h1SMcG0008035
h1SMcG0013999
h1SMcG0013627
h1SMcG0006433
h1SMcG0013162
h1SMcG0003097
h1SMcG0020537
UMAP1
UMAP2

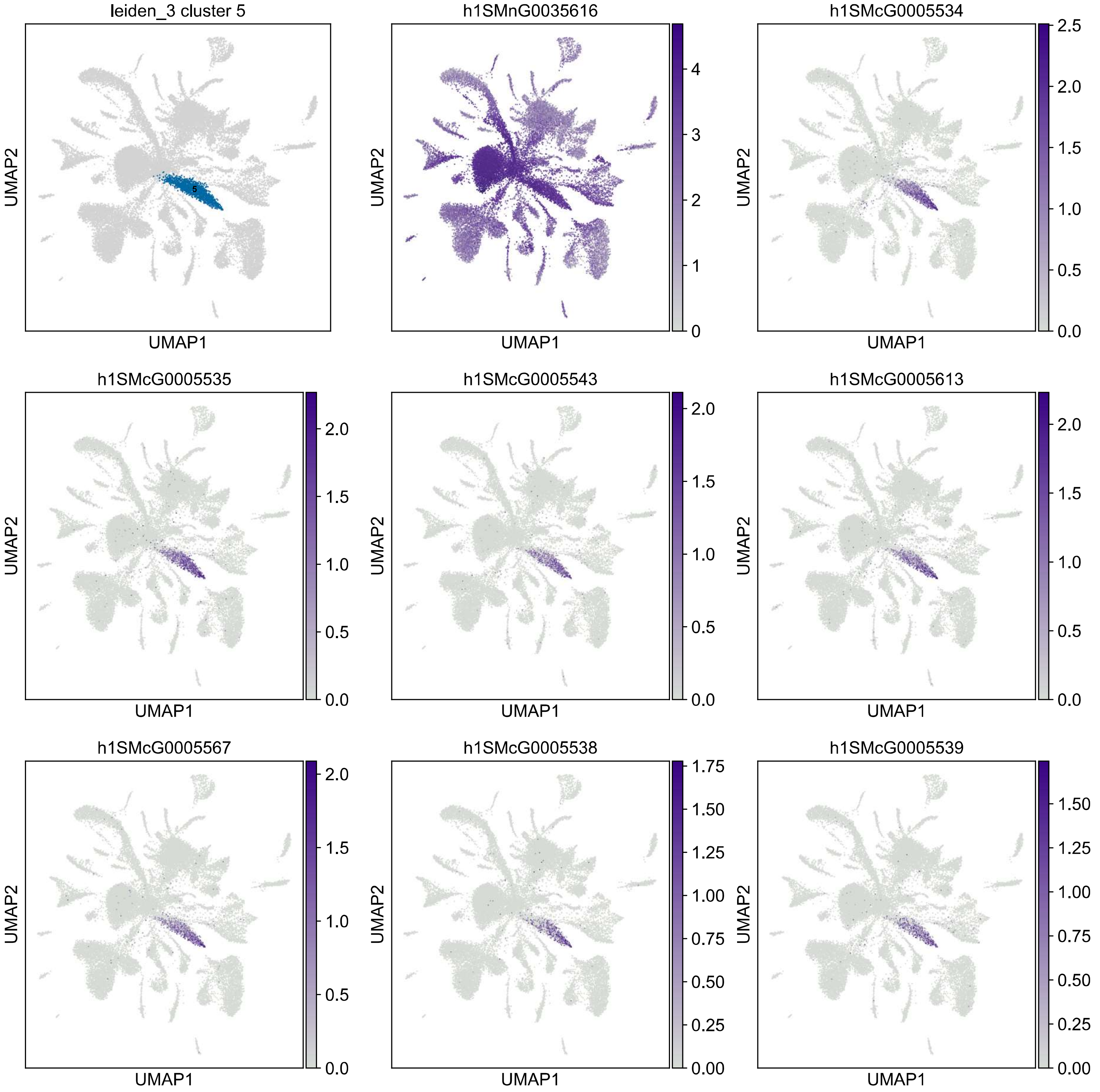

leiden_3 cluster 5
5
h1SMnG0035616
h1SMcG0005534
h1SMcG0005535
h1SMcG0005543
h1SMcG0005613
h1SMcG0005567
h1SMcG0005538
h1SMcG0005539
UMAP1
UMAP2
0
1
2
3
4
0.0
0.5
1.0
1.5
2.0
2.5
0.00
0.25
0.50
0.75
1.00
1.25
1.50
1.75

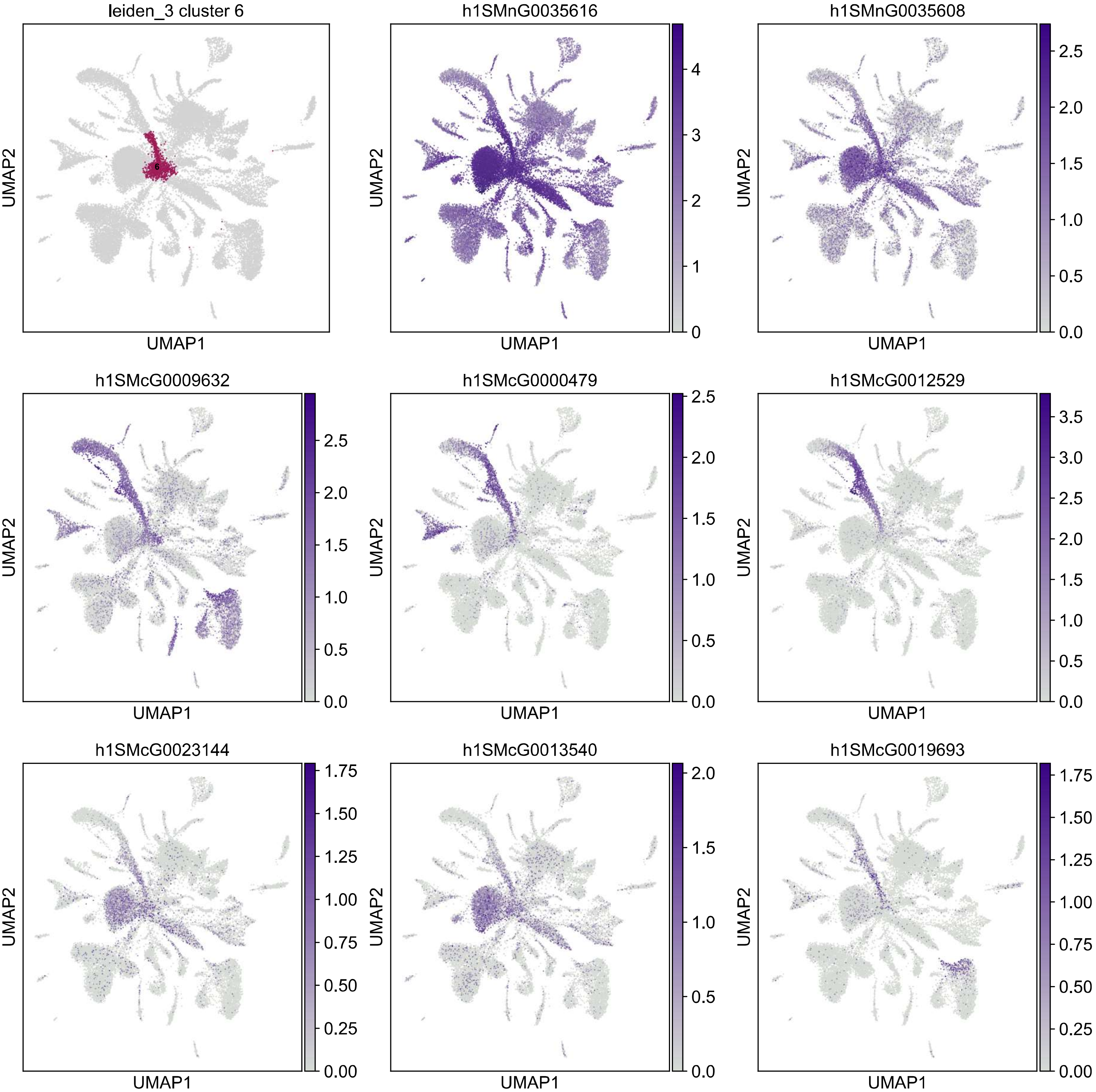
leiden_3 cluster 6
h1SMnG0035616
h1SMnG0035608
h1SMcG0009632
h1SMcG0000479
h1SMcG0012529
h1SMcG0023144
h1SMcG0013540
h1SMcG0019693
UMAP1
UMAP2

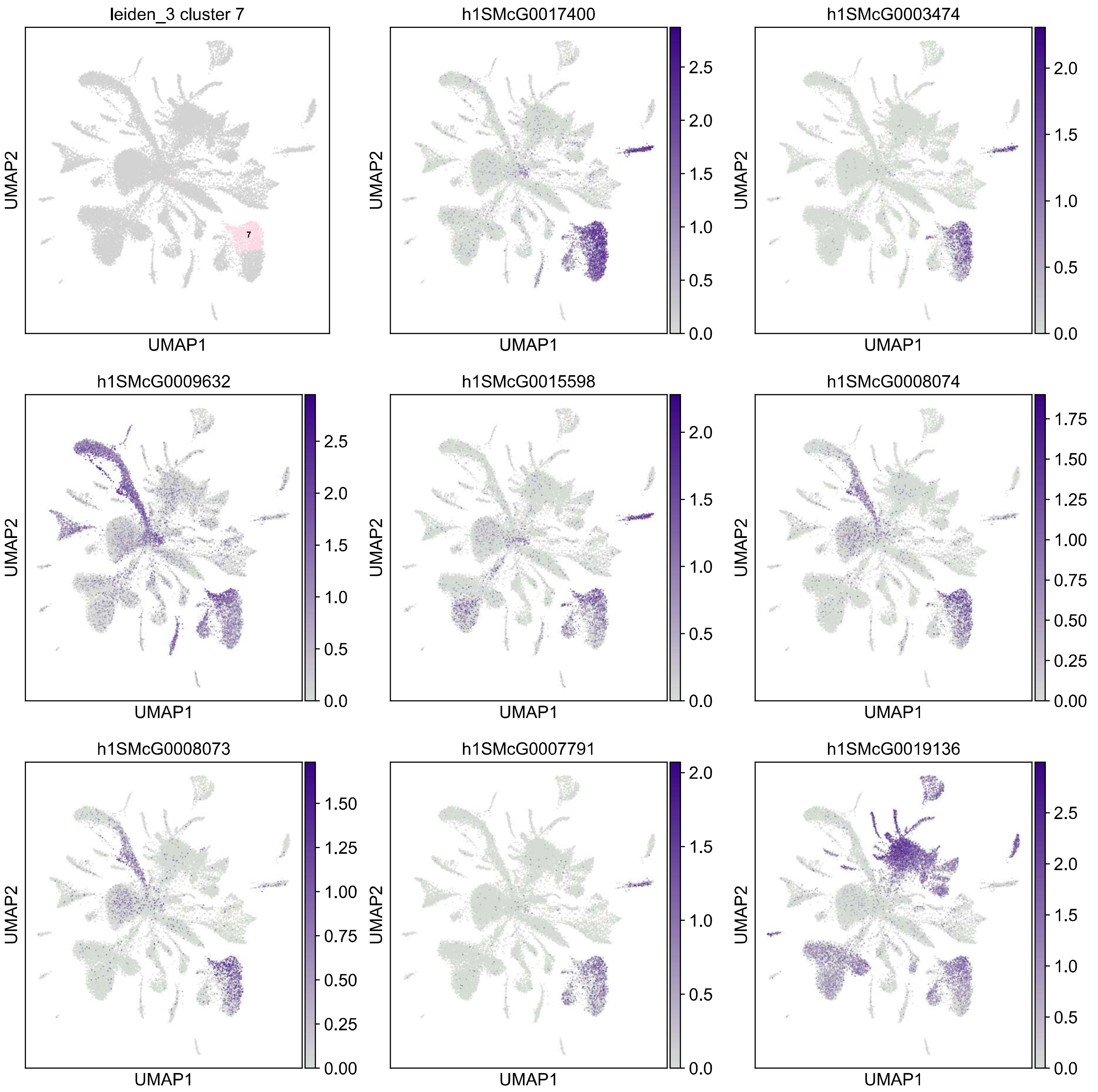

leiden_3 cluster 7
7
h1SMcG0017400
h1SMcG0003474
h1SMcG0009632
h1SMcG0015598
h1SMcG0008074
h1SMcG0008073
h1SMcG0007791
h1SMcG0019136
UMAP1
UMAP2

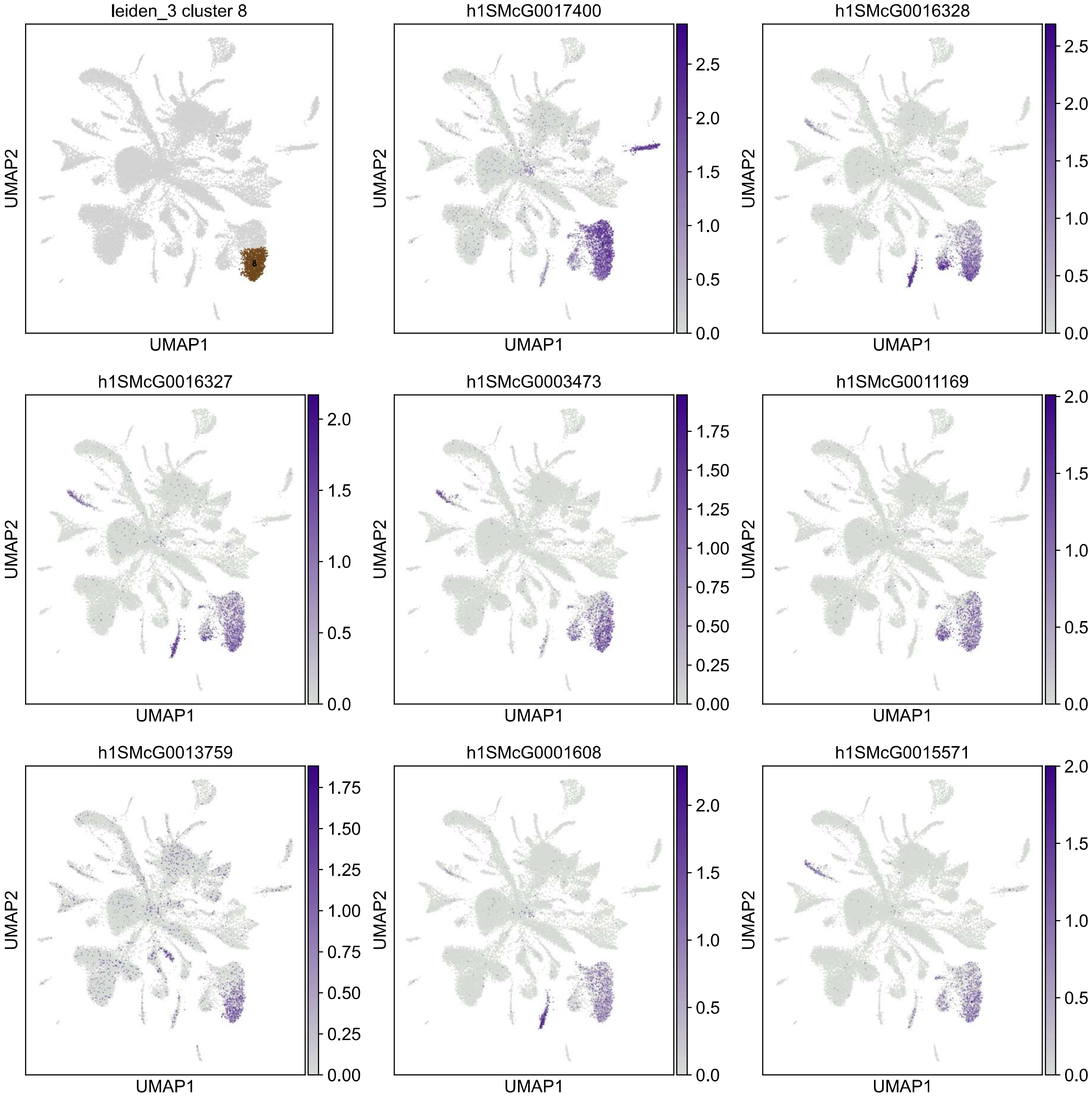

leiden_3 cluster 8
h1SMcG0017400
h1SMcG0016328
h1SMcG0016327
h1SMcG0003473
h1SMcG0011169
h1SMcG0013759
h1SMcG0001608
h1SMcG0015571
UMAP1
UMAP2

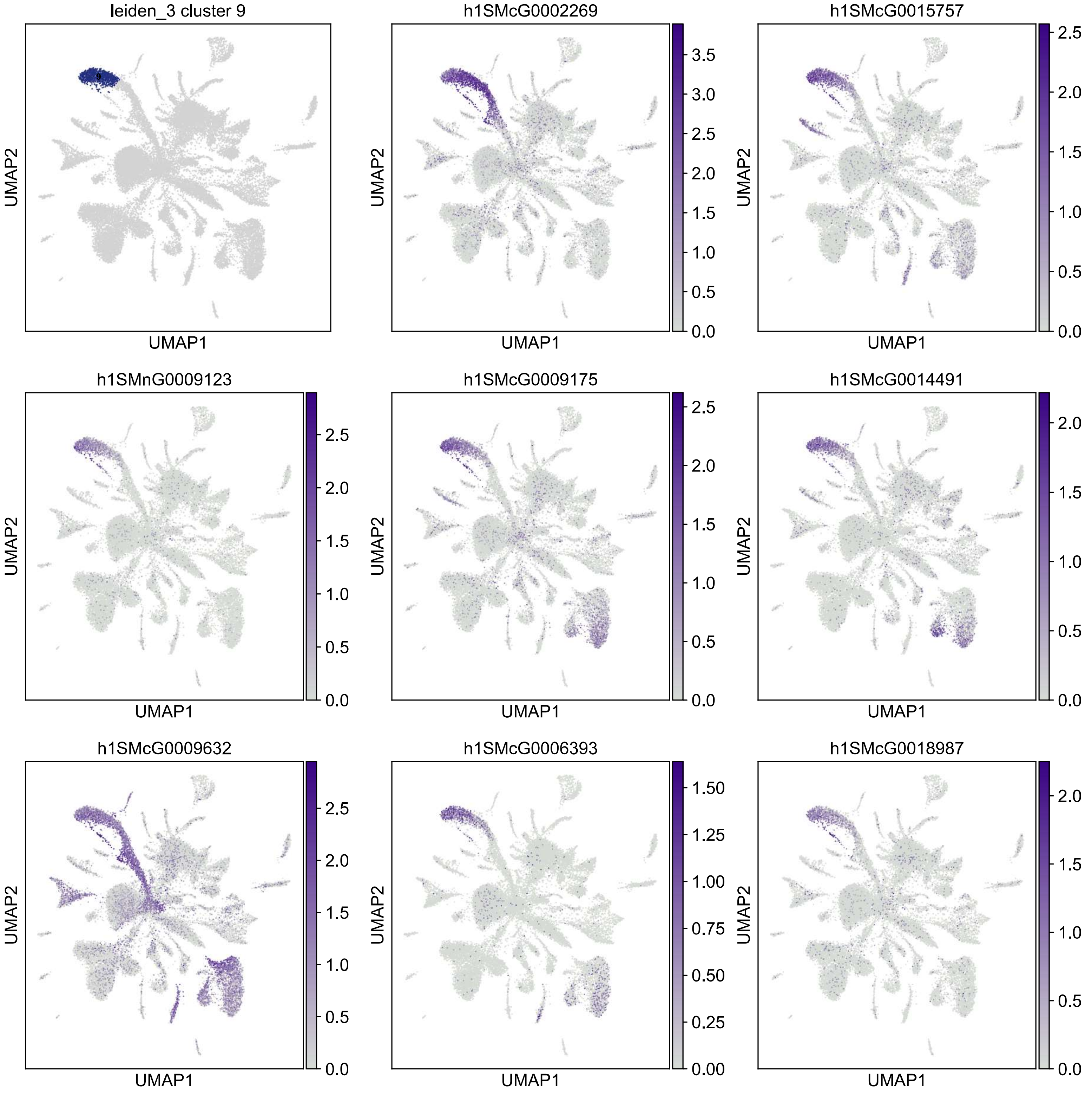
leiden_3 cluster 9
h1SMcG0002269
h1SMcG0015757
h1SMnG0009123
h1SMcG0009175
h1SMcG0014491
h1SMcG0009632
h1SMcG0006393
h1SMcG0018987
UMAP1
UMAP2

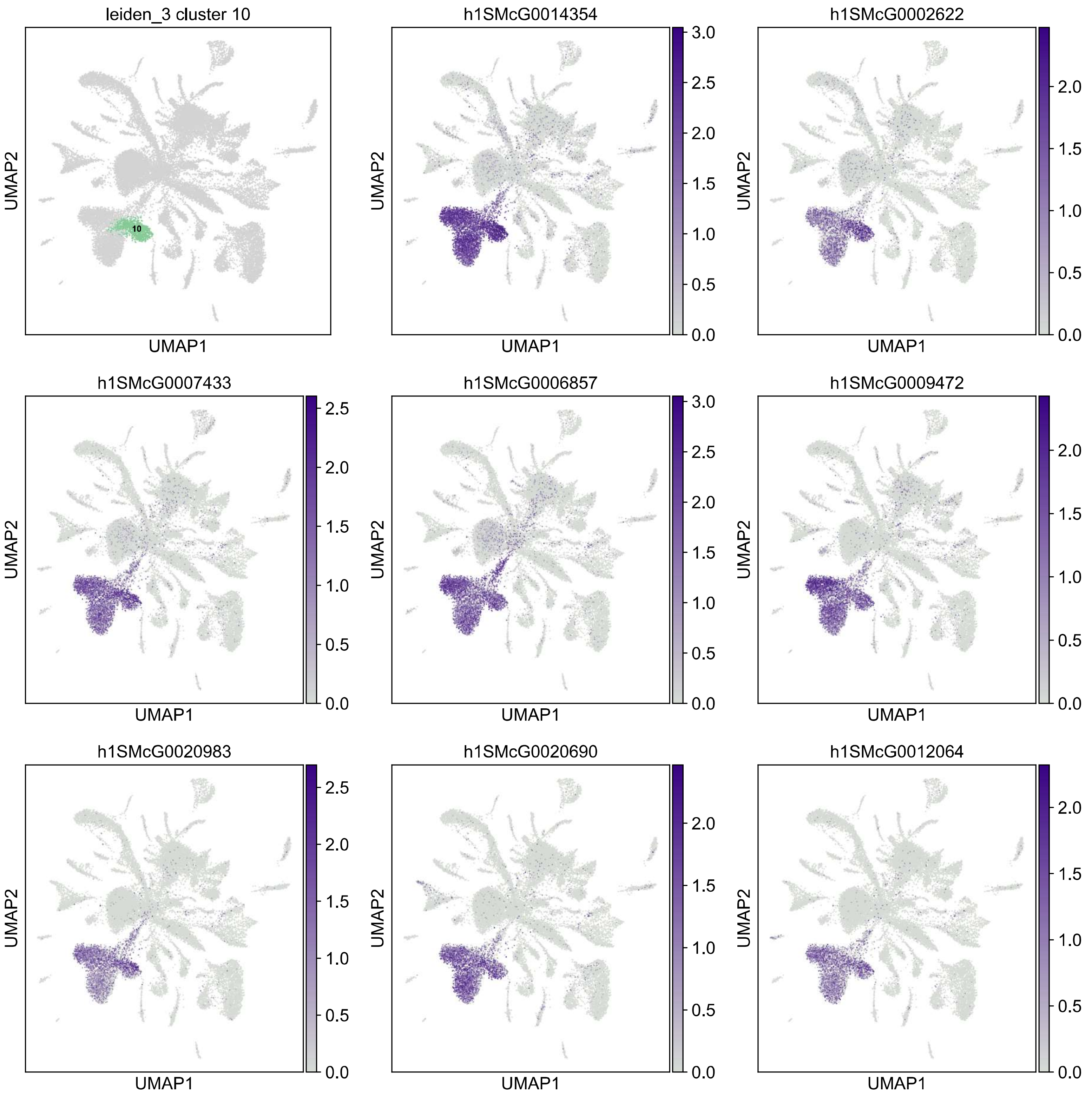

leiden_3 cluster 10
10
h1SMcG0014354
h1SMcG0002622
h1SMcG0007433
h1SMcG0006857
h1SMcG0009472
h1SMcG0020983
h1SMcG0020690
h1SMcG0012064
UMAP1
UMAP2

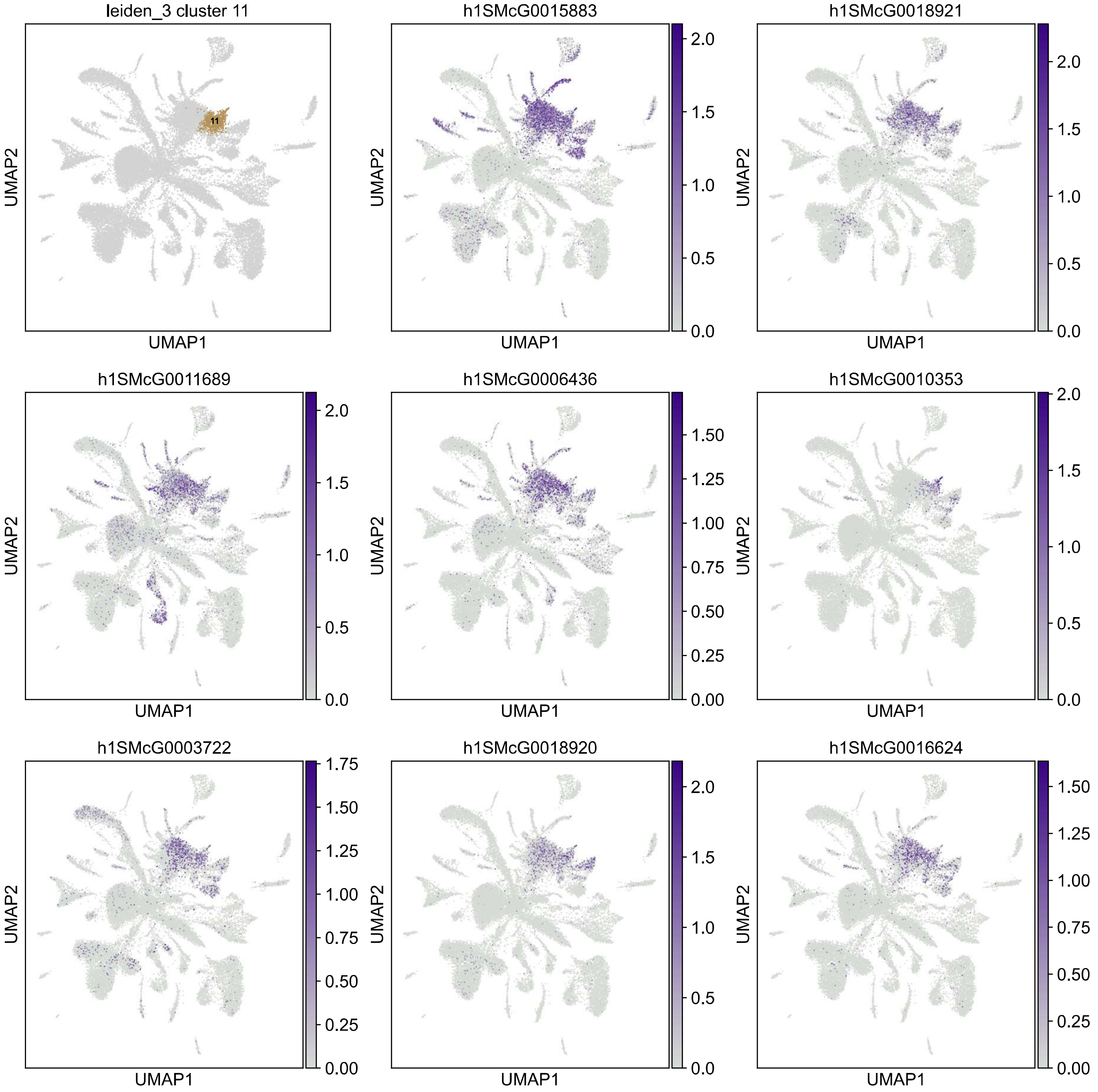

leiden_3 cluster 11
11
h1SMcG0015883
h1SMcG0018921
h1SMcG0011689
h1SMcG0006436
h1SMcG0010353
h1SMcG0003722
h1SMcG0018920
h1SMcG0016624
UMAP1
UMAP2

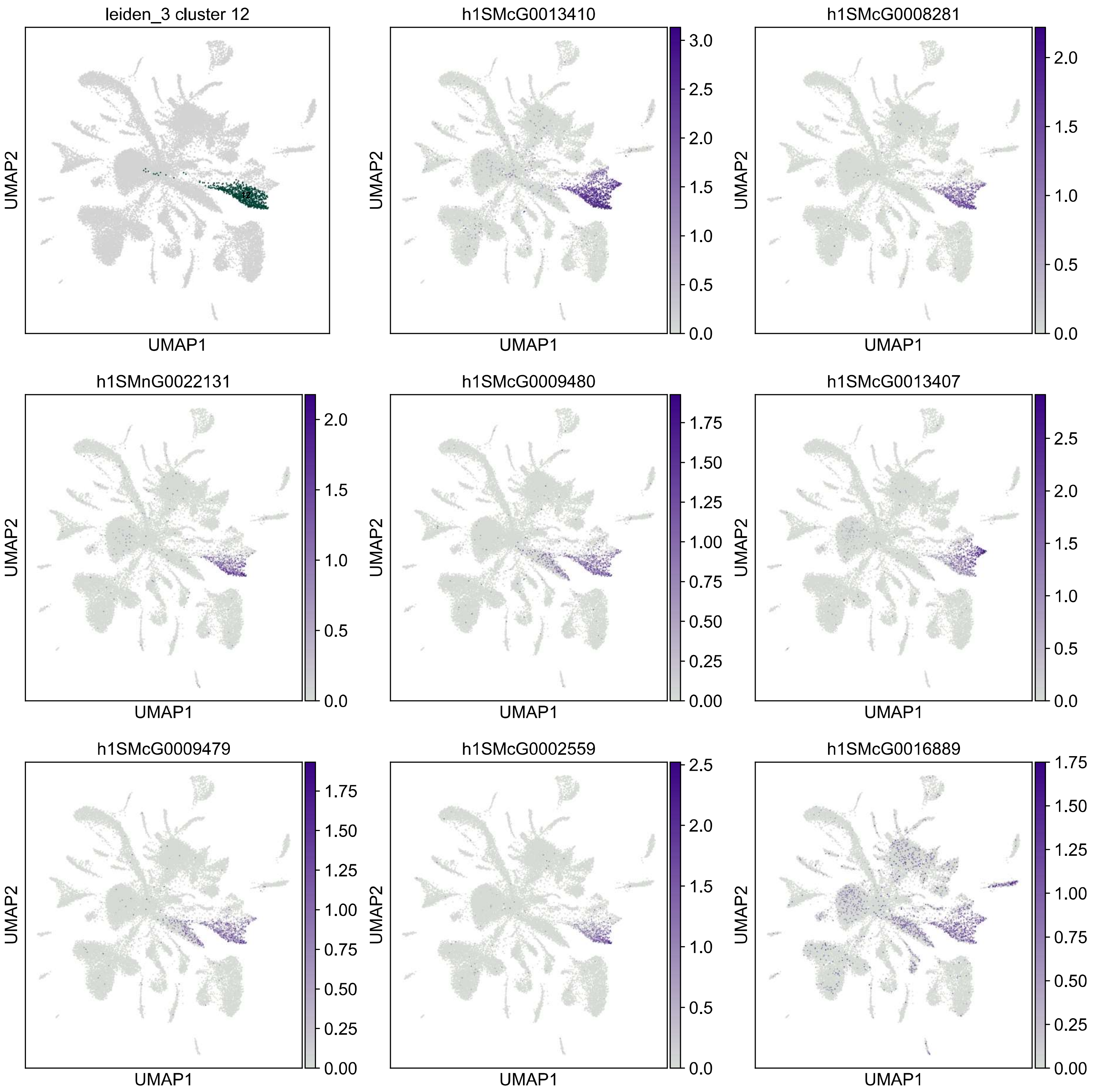
leiden_3 cluster 12
h1SMcG0013410
h1SMcG0008281
h1SMnG0022131
h1SMcG0009480
h1SMcG0013407
h1SMcG0009479
h1SMcG0002559
h1SMcG0016889
UMAP1
UMAP2

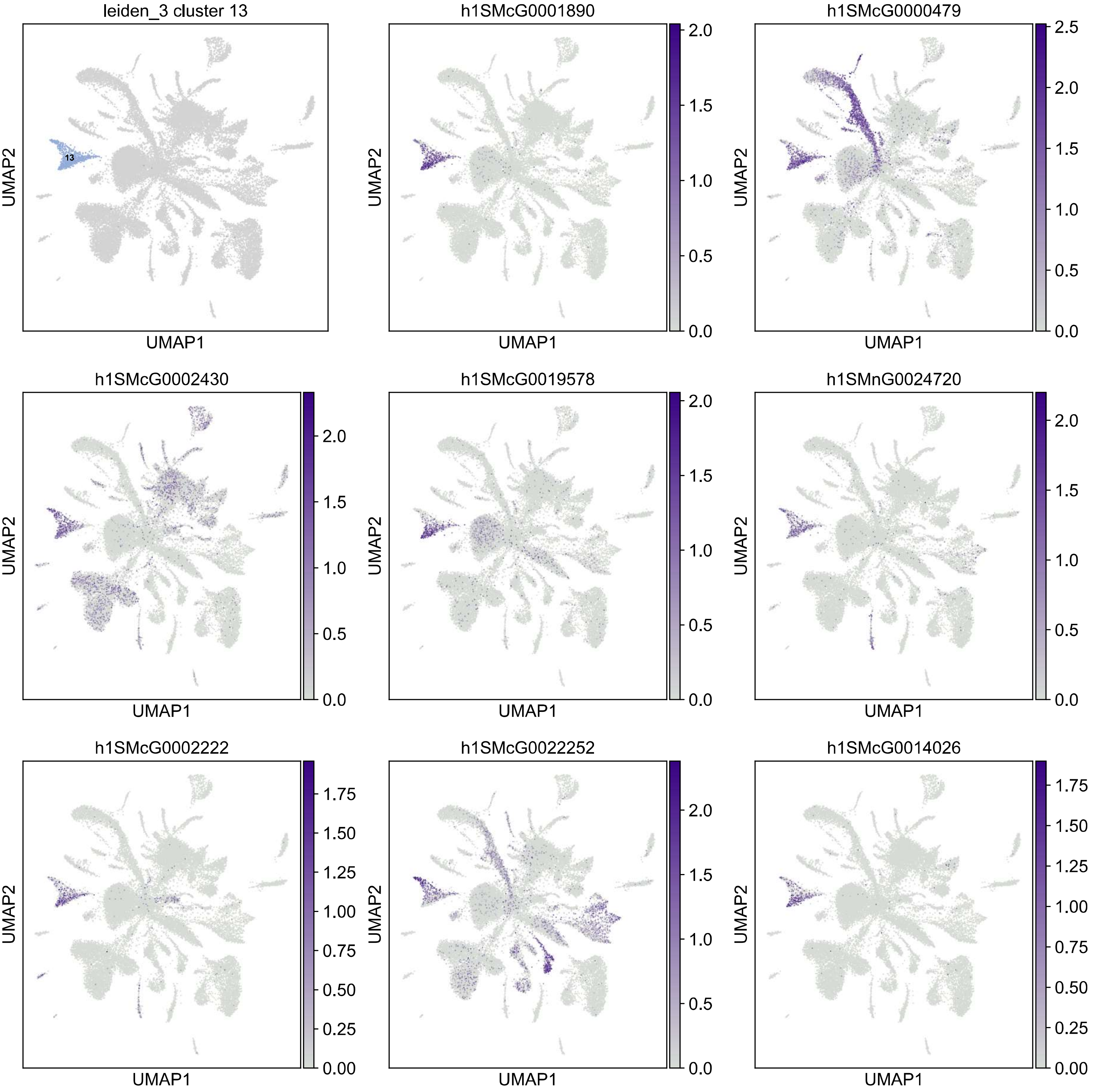
leiden_3 cluster 13
13
h1SMcG0001890
h1SMcG0000479
h1SMcG0002430
h1SMcG0019578
h1SMnG0024720
h1SMcG0002222
h1SMcG0022252
h1SMcG0014026
UMAP1
UMAP2

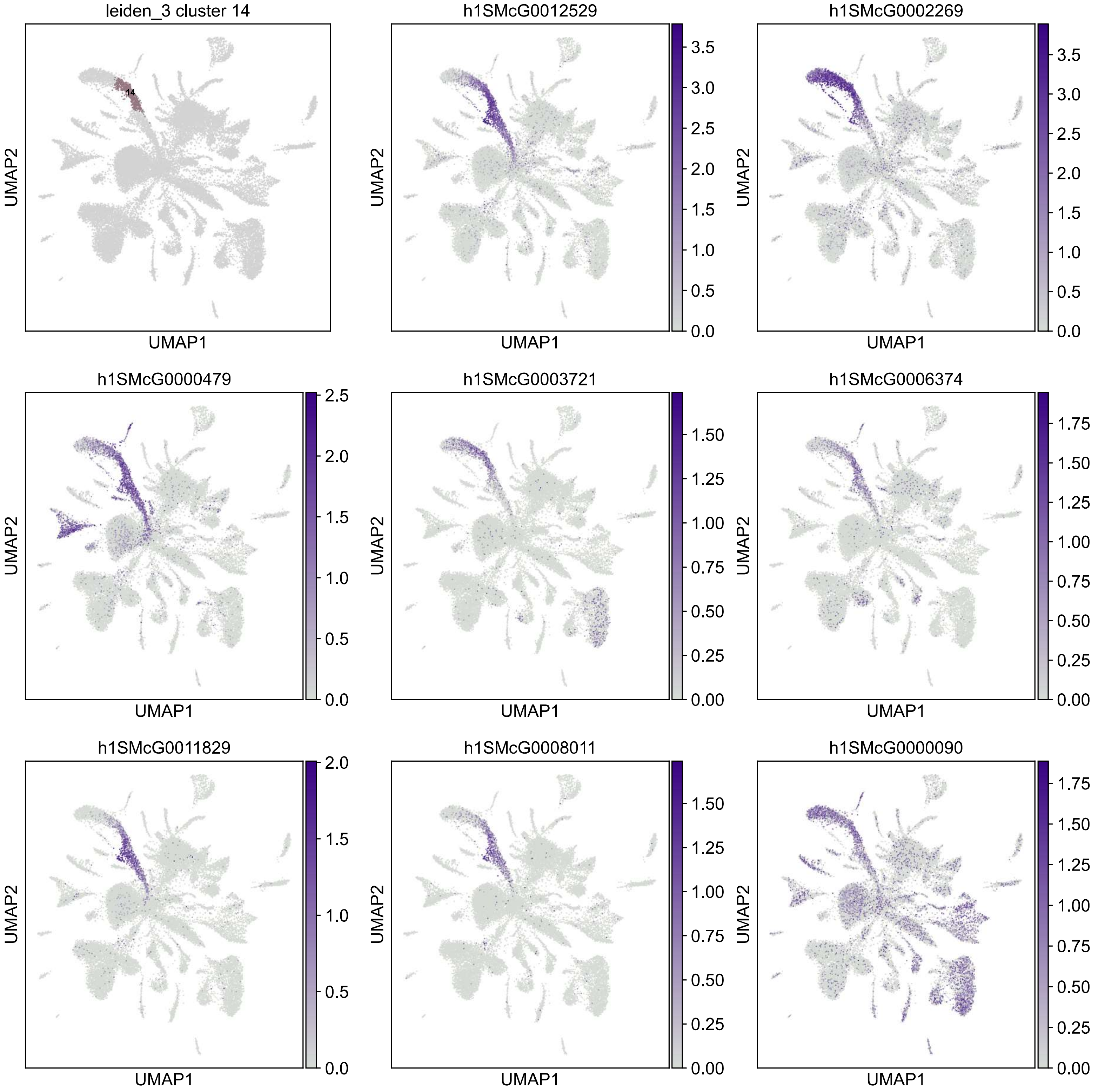

leiden_3 cluster 14
14
UMAP1
UMAP2
h1SMcG0012529
h1SMcG0002269
h1SMcG0000479
h1SMcG0003721
h1SMcG0006374
h1SMcG0011829
h1SMcG0008011
h1SMcG0000090

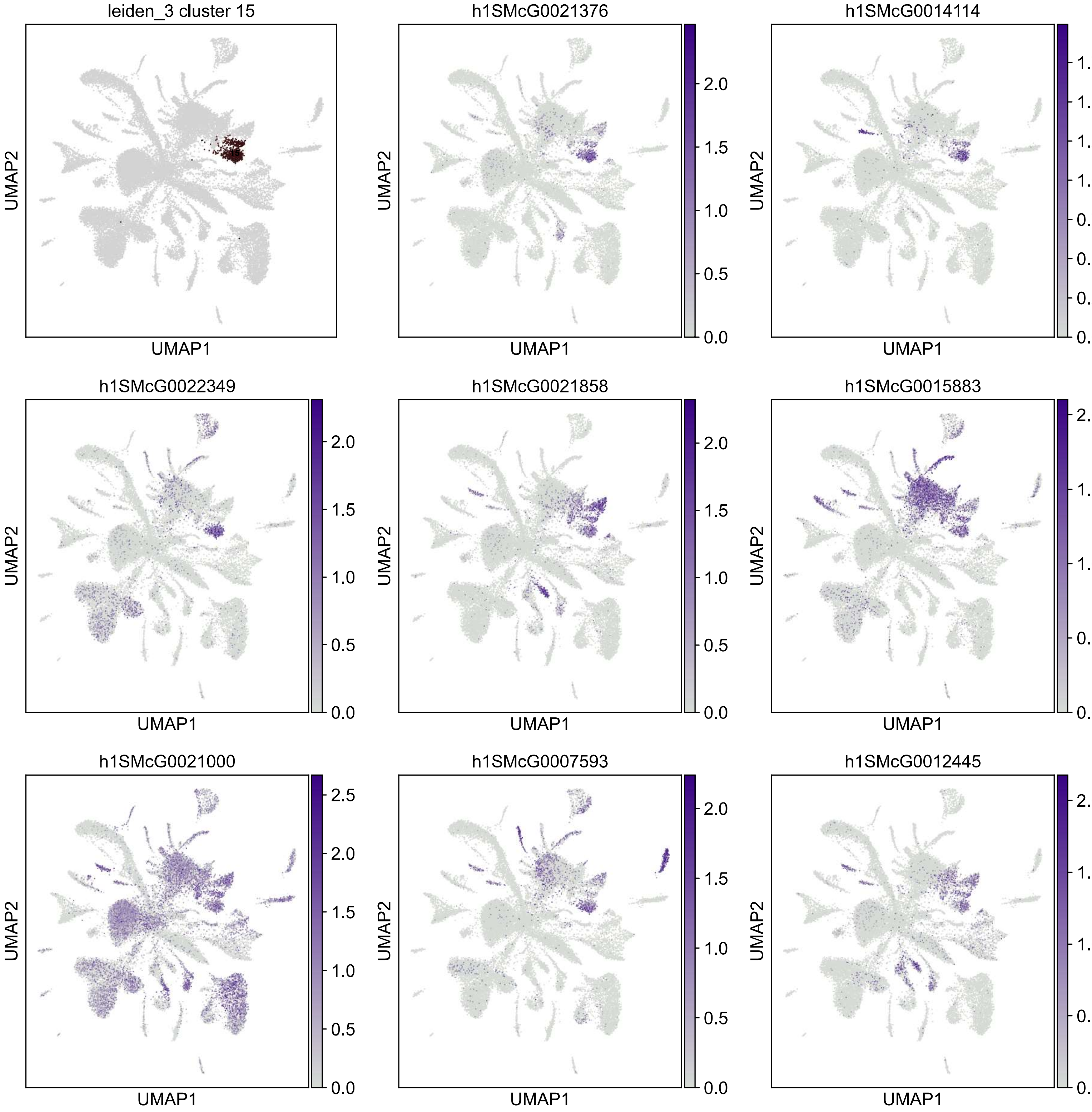
leiden_3 cluster 15
h1SMcG0021376
h1SMcG0014114
h1SMcG0022349
h1SMcG0021858
h1SMcG0015883
h1SMcG0021000
h1SMcG0007593
h1SMcG0012445
UMAP1
UMAP2

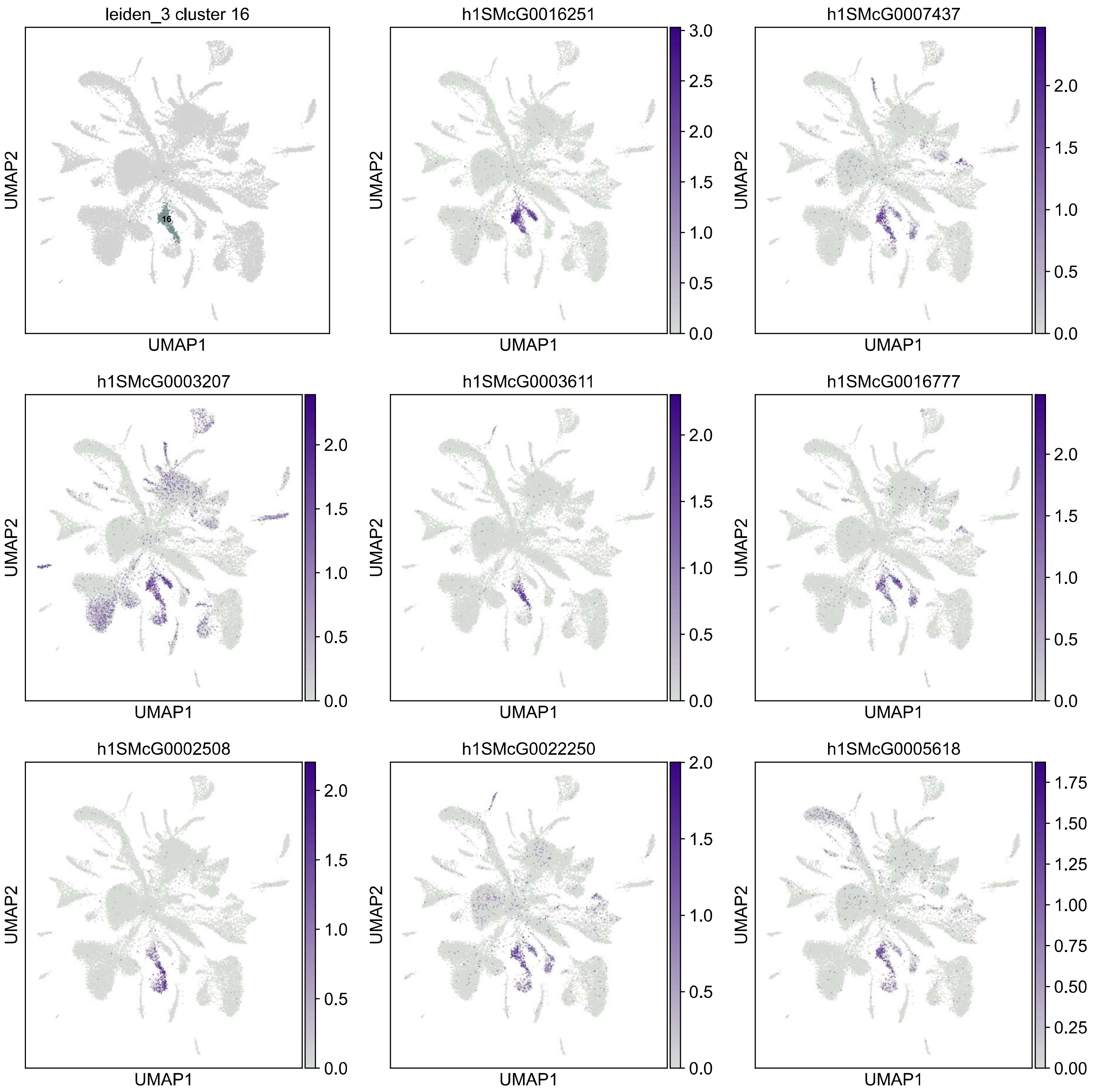
leiden_3 cluster 16
16
h1SMcG0016251
h1SMcG0007437
h1SMcG0003207
h1SMcG0003611
h1SMcG0016777
h1SMcG0002508
h1SMcG0022250
h1SMcG0005618
UMAP1
UMAP2

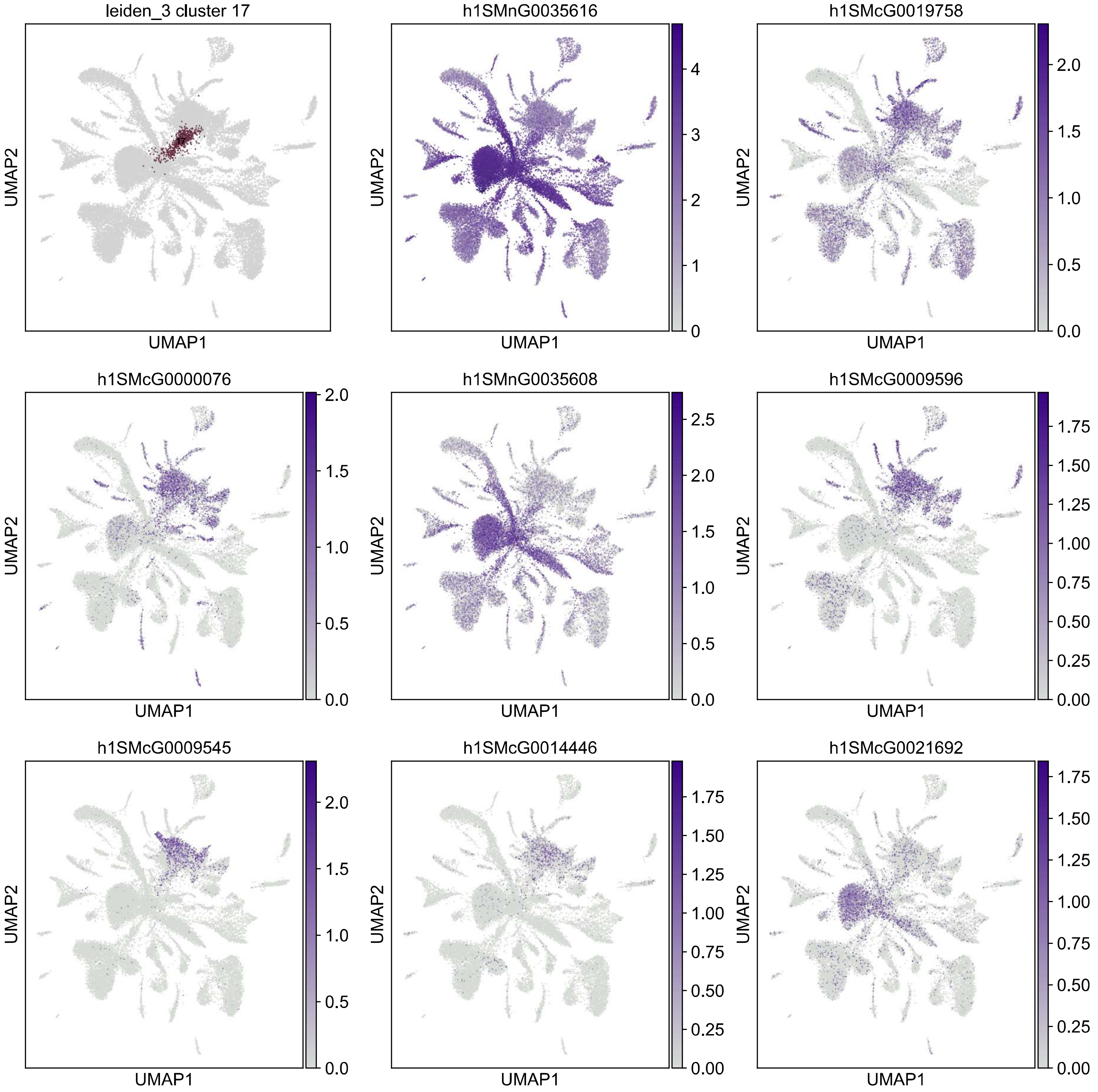

leiden_3 cluster 17
h1SMnG0035616
h1SMcG0019758
h1SMcG0000076
h1SMnG0035608
h1SMcG0009596
h1SMcG0009545
h1SMcG0014446
h1SMcG0021692
UMAP1
UMAP2

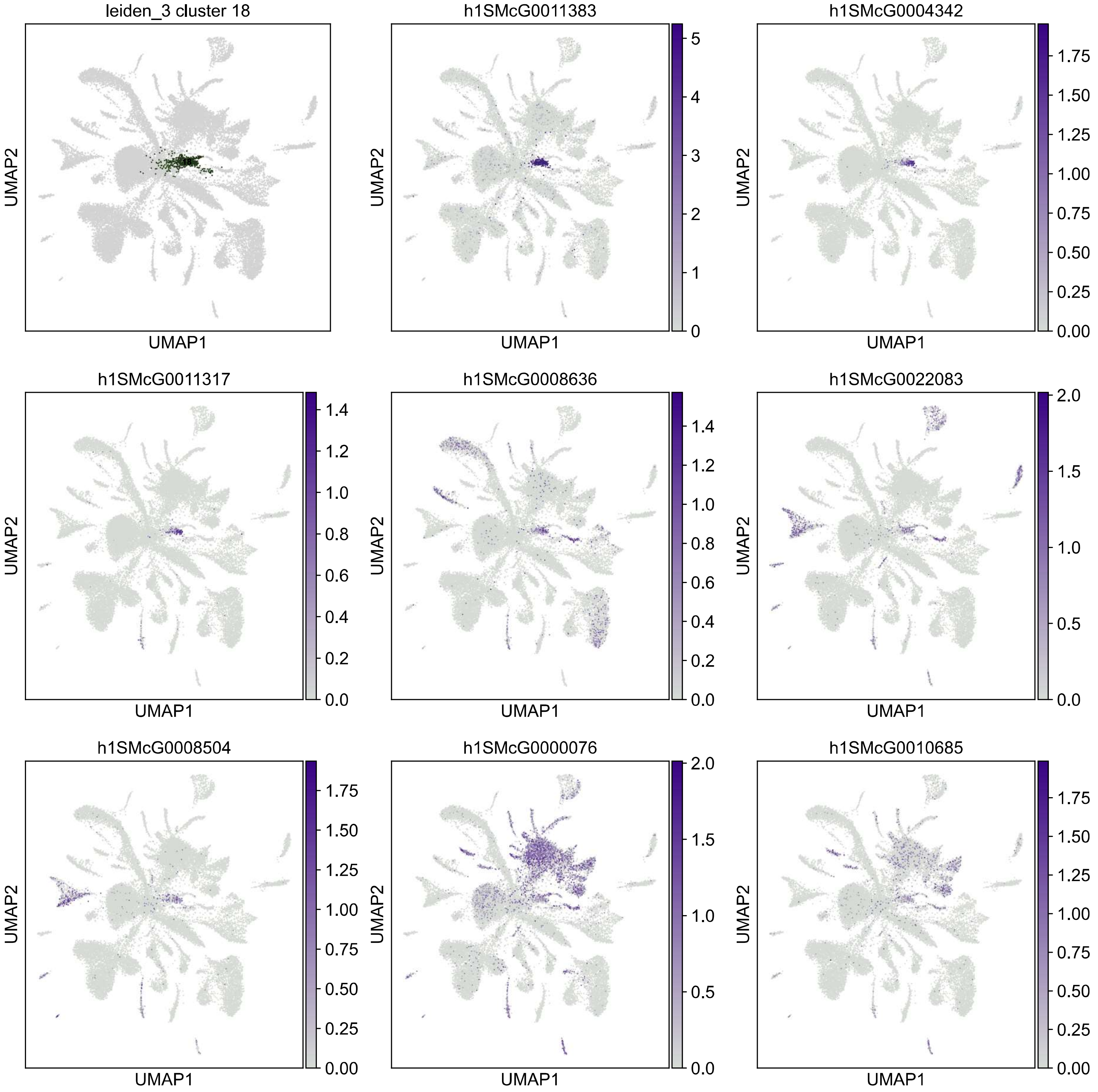

leiden_3 cluster 18
h1SMcG0011383
h1SMcG0004342
h1SMcG0011317
h1SMcG0008636
h1SMcG0022083
h1SMcG0008504
h1SMcG0000076
h1SMcG0010685
UMAP1
UMAP2

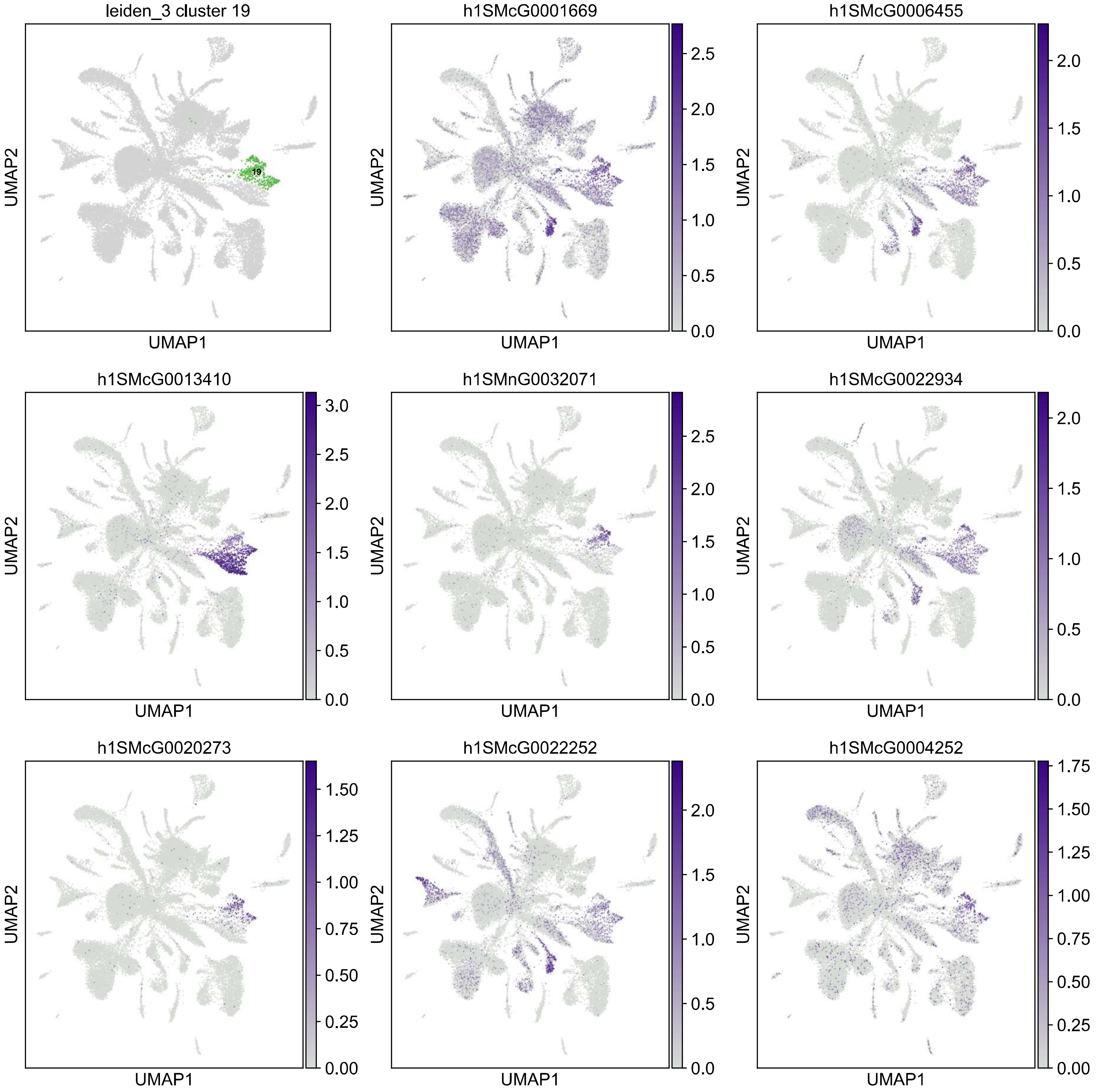

leiden_3 cluster 19
19
h1SMcG0001669
h1SMcG0006455
h1SMcG0013410
h1SMnG0032071
h1SMcG0022934
h1SMcG0020273
h1SMcG0022252
h1SMcG0004252
UMAP1
UMAP2

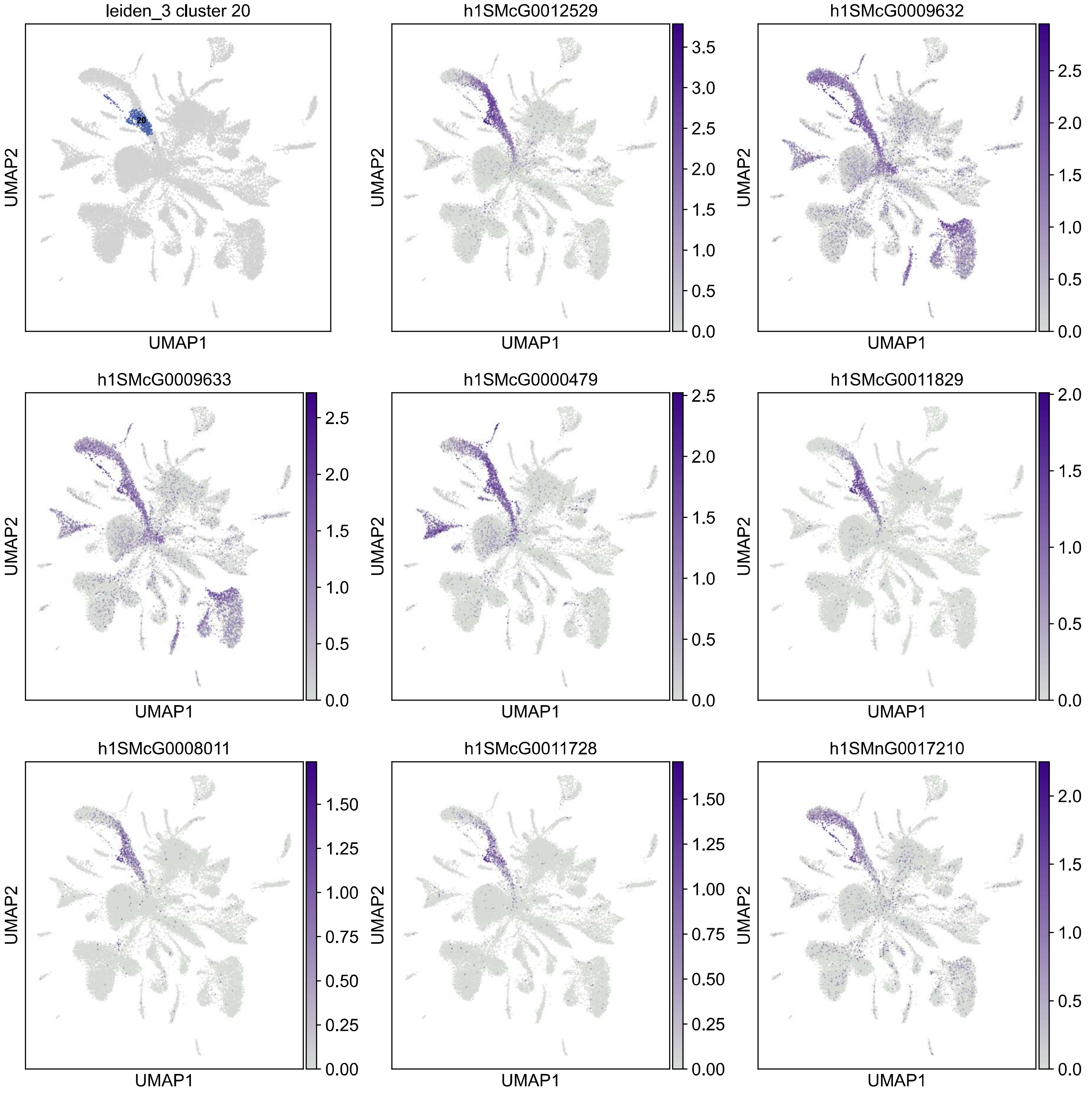

leiden_3 cluster 20
20
h1SMcG0012529
h1SMcG0009632
h1SMcG0009633
h1SMcG0000479
h1SMcG0011829
h1SMcG0008011
h1SMcG0011728
h1SMnG0017210
UMAP1
UMAP2

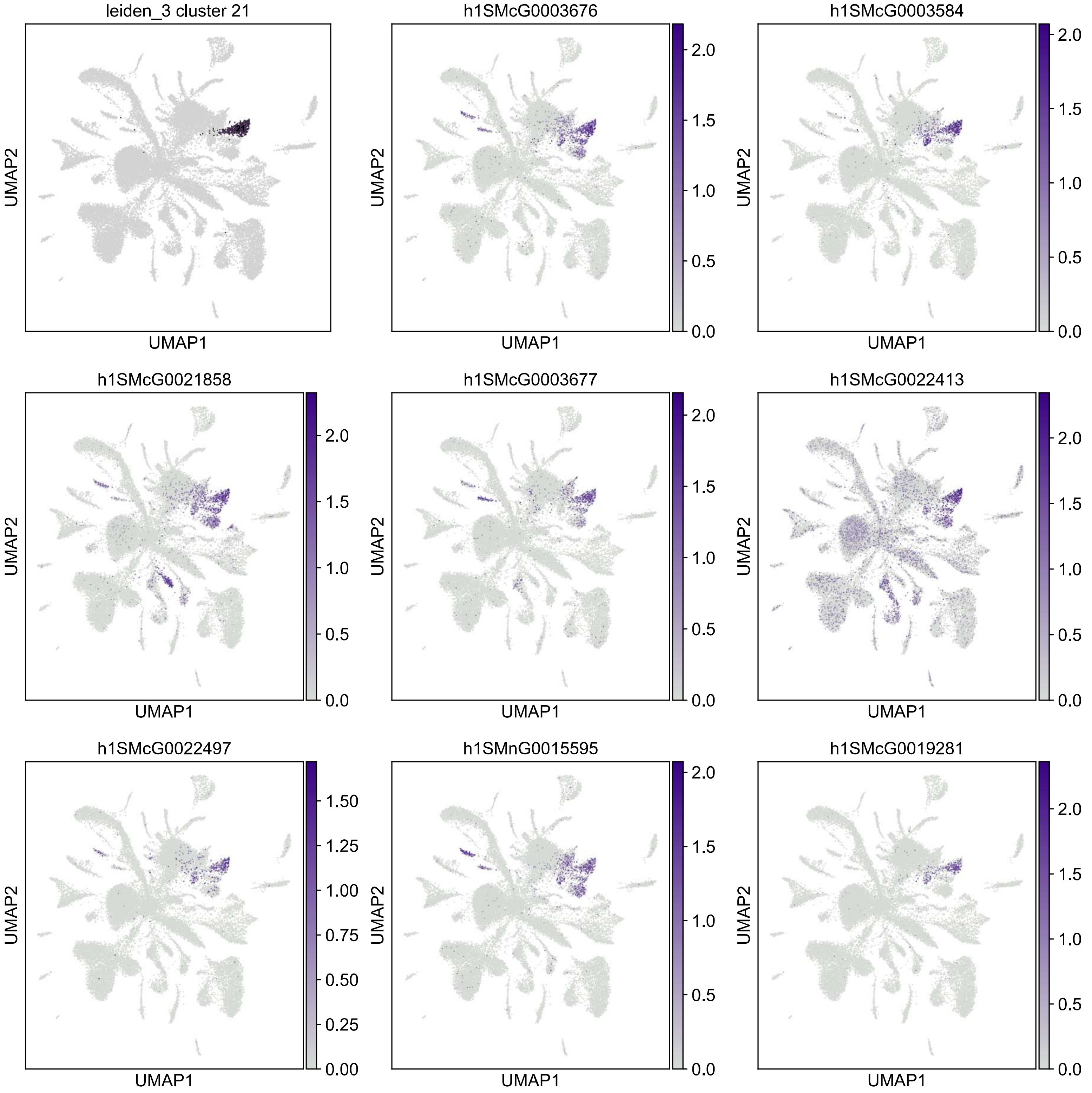

leiden_3 cluster 21
h1SMcG0003676
h1SMcG0003584
h1SMcG0021858
h1SMcG0003677
h1SMcG0022413
h1SMcG0022497
h1SMnG0015595
h1SMcG0019281
UMAP1
UMAP2
0.0
0.5
1.0
1.5
2.0
0.00
0.25
0.50
0.75
1.00
1.25
1.50

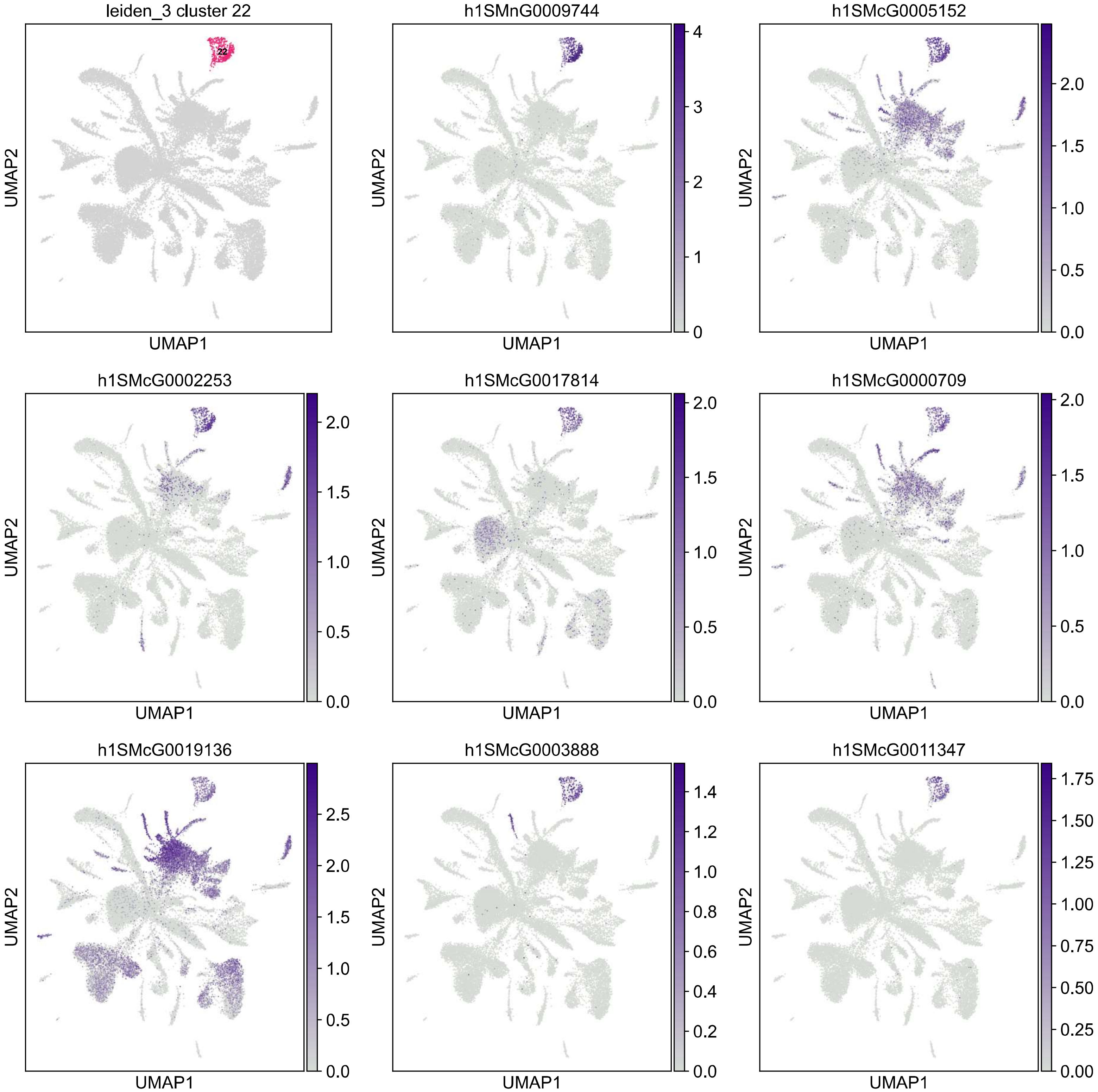

leiden_3 cluster 22
22
h1SMnG0009744
h1SMcG0005152
h1SMcG0002253
h1SMcG0017814
h1SMcG0000709
h1SMcG0019136
h1SMcG0003888
h1SMcG0011347
UMAP1
UMAP2

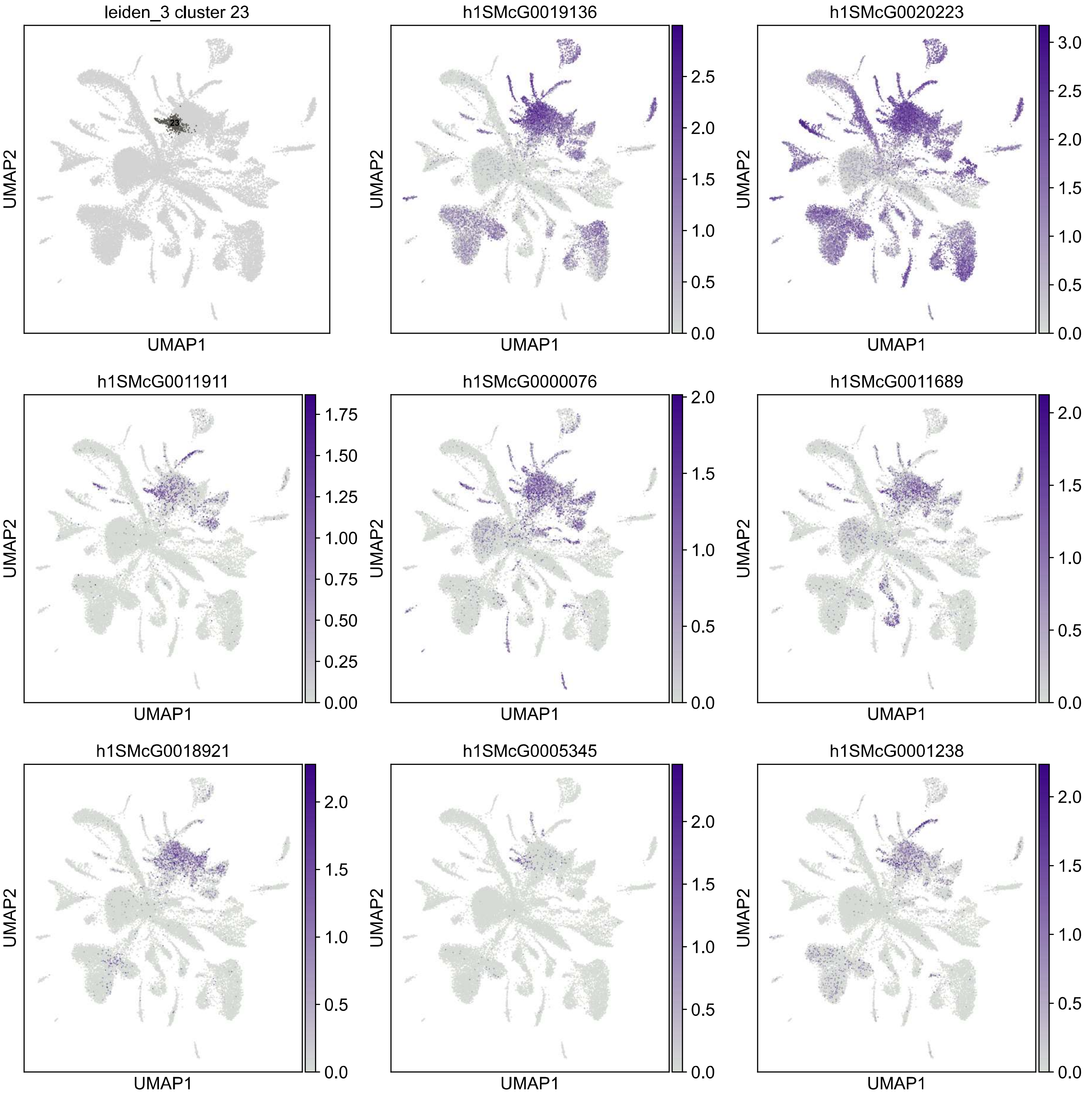
leiden_3 cluster 23
23
h1SMcG0019136
h1SMcG0020223
h1SMcG0011911
h1SMcG0000076
h1SMcG0011689
h1SMcG0018921
h1SMcG0005345
h1SMcG0001238
UMAP1
UMAP2
0.0 0.5 1.0 1.5 2.0 2.5 3.0
0.00 0.25 0.50 0.75 1.00 1.25 1.50 1.75

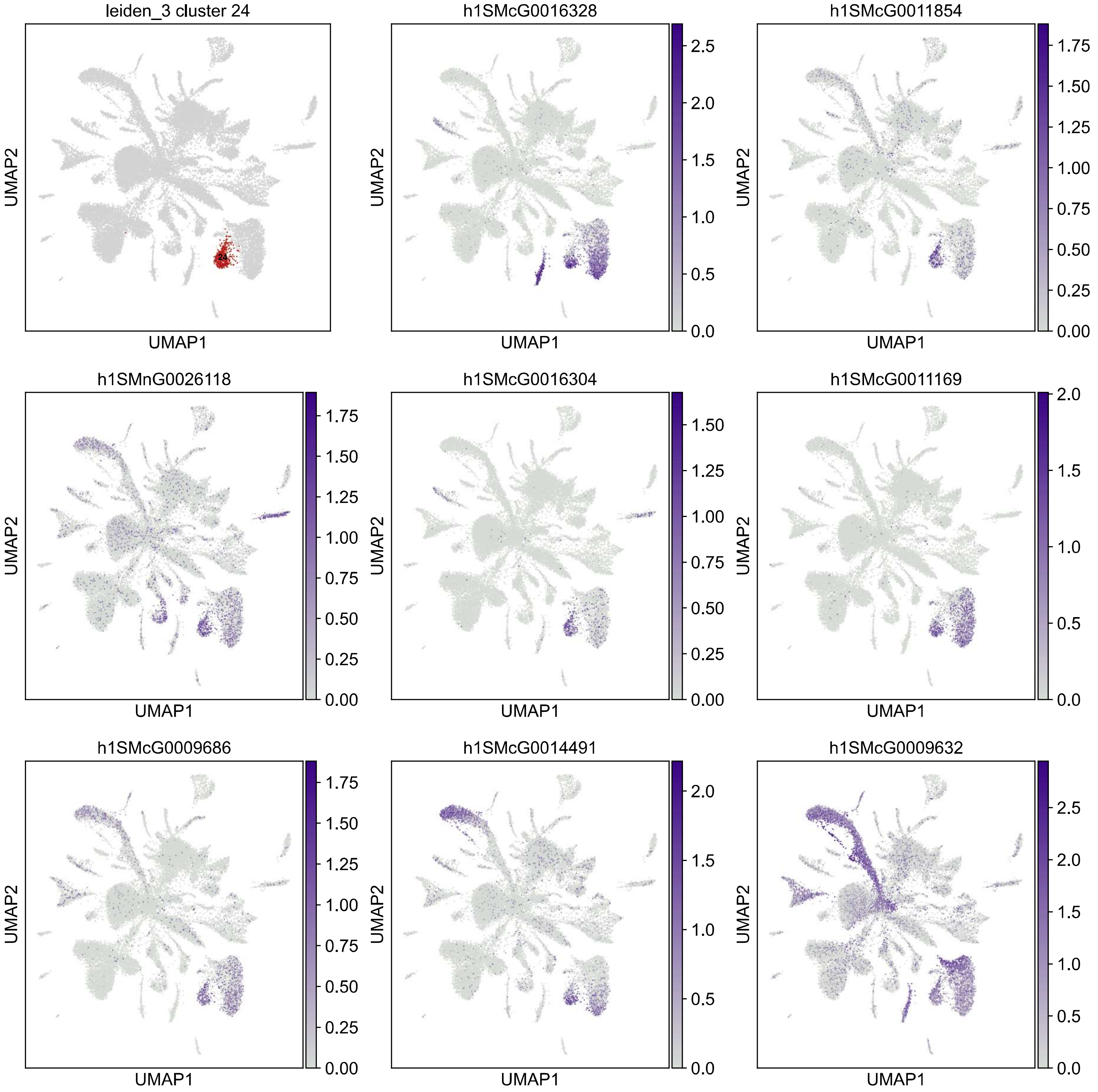

leiden_3 cluster 24
24
h1SMcG0016328
h1SMcG0011854
h1SMnG0026118
h1SMcG0016304
h1SMcG0011169
h1SMcG0009686
h1SMcG0014491
h1SMcG0009632
UMAP1
UMAP2

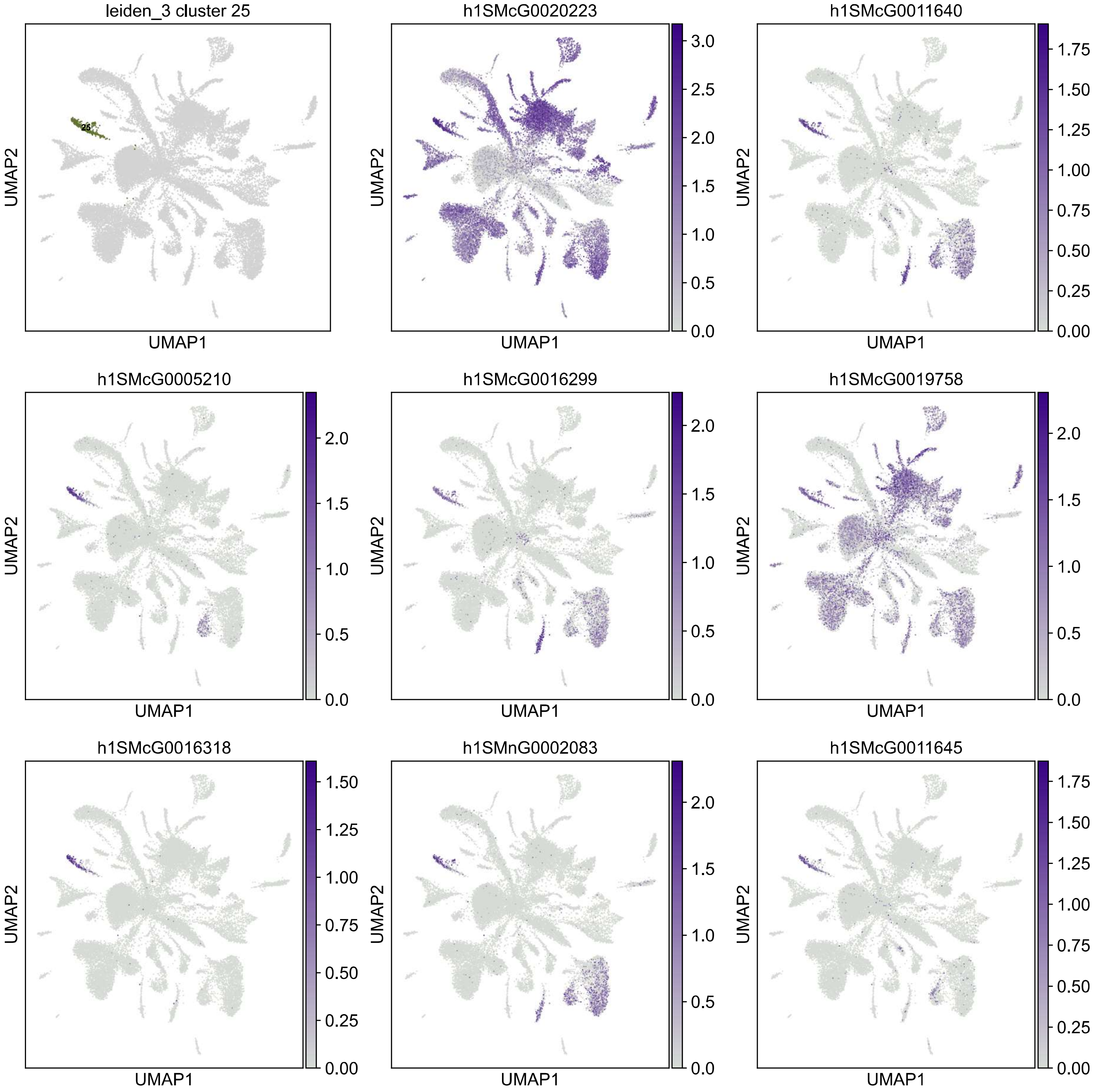

leiden_3 cluster 25
25
h1SMcG0020223
h1SMcG0011640
h1SMcG0005210
h1SMcG0016299
h1SMcG0019758
h1SMcG0016318
h1SMnG0002083
h1SMcG0011645
UMAP1
UMAP2

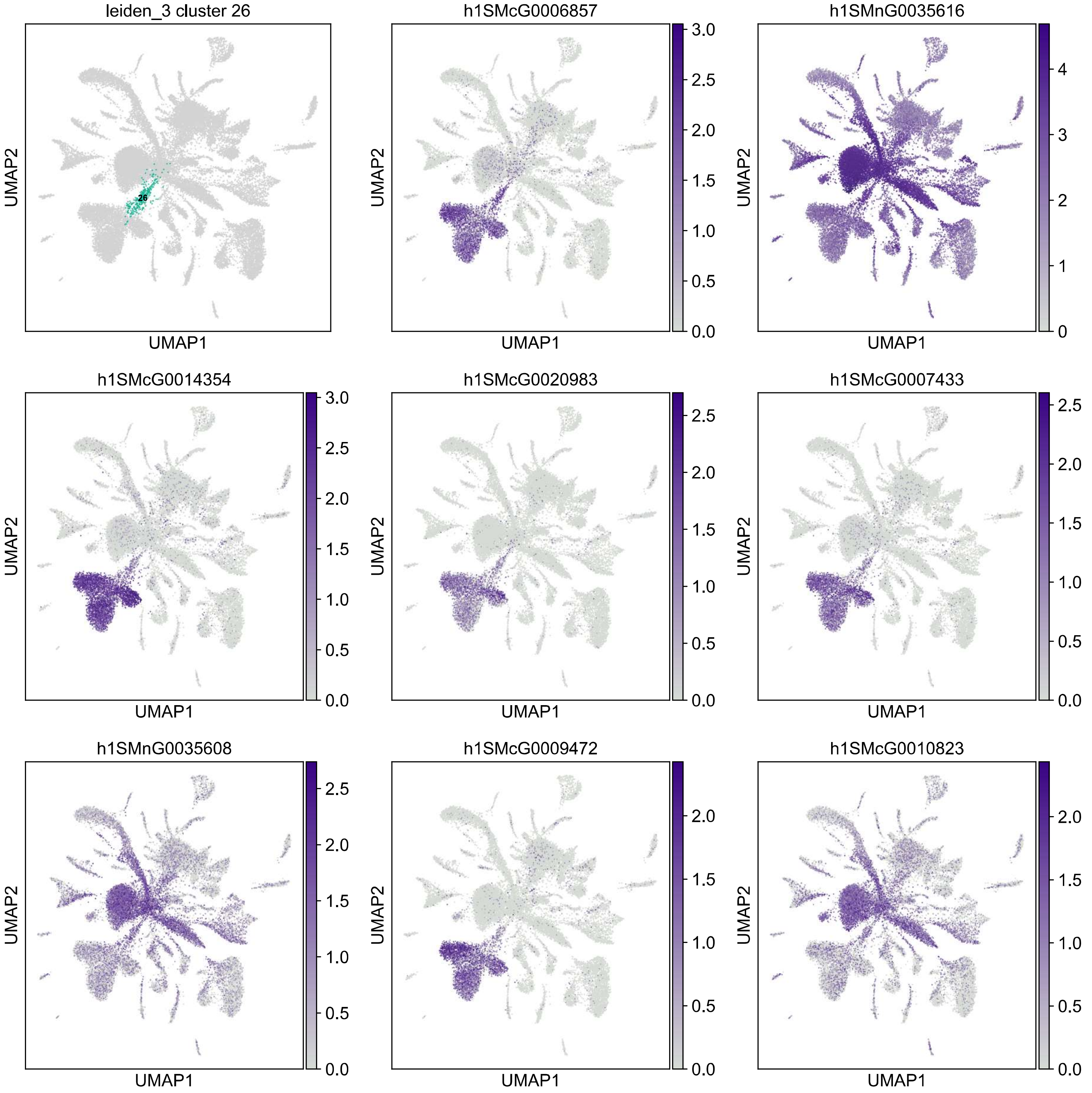

leiden_3 cluster 26
26
h1SMcG0006857
h1SMnG0035616
h1SMcG0014354
h1SMcG0020983
h1SMcG0007433
h1SMnG0035608
h1SMcG0009472
h1SMcG0010823
UMAP1
UMAP2

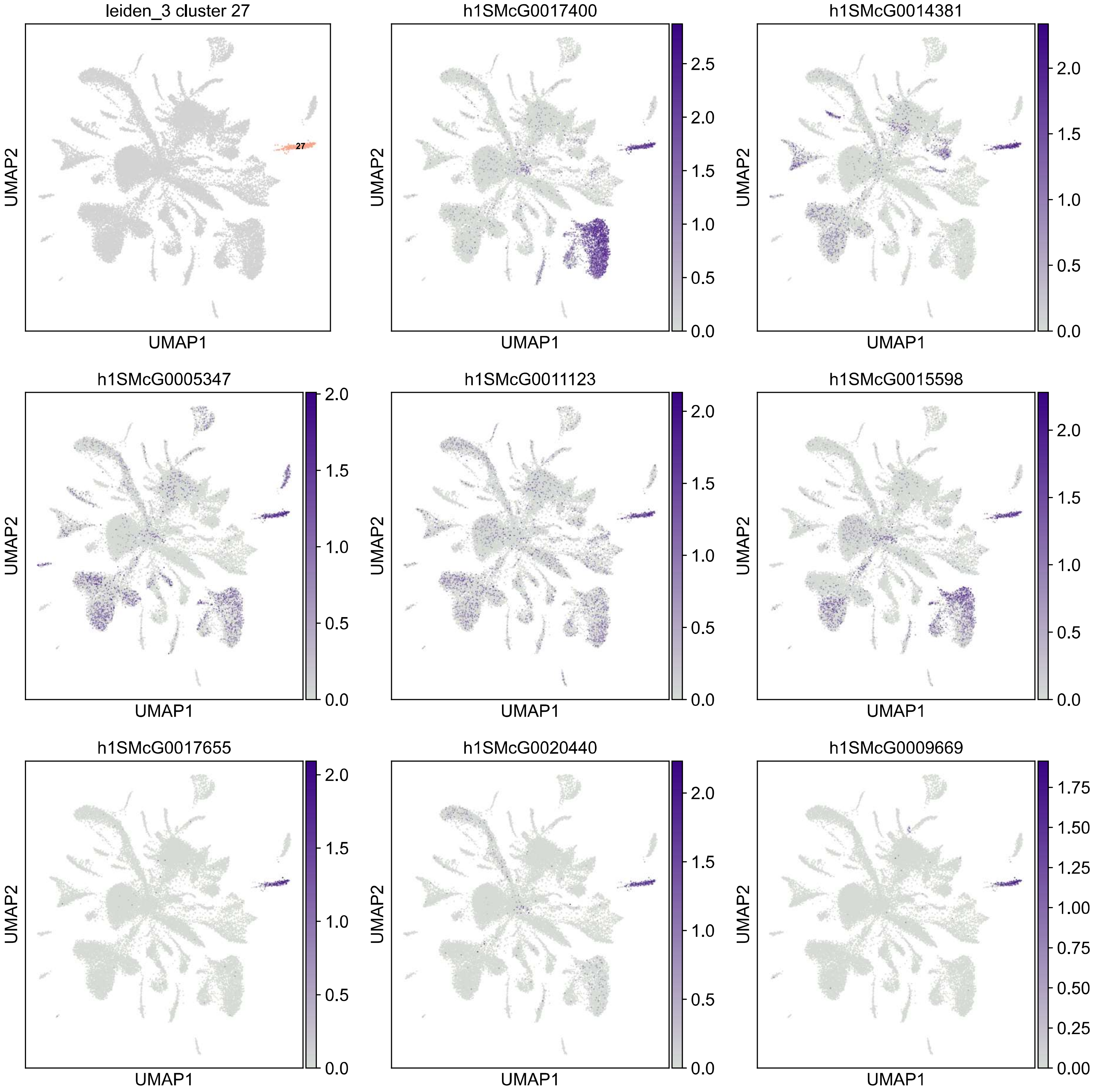

leiden_3 cluster 27
27
h1SMcG0017400
h1SMcG0014381
h1SMcG0005347
h1SMcG0011123
h1SMcG0015598
h1SMcG0017655
h1SMcG0020440
h1SMcG0009669
UMAP1
UMAP2

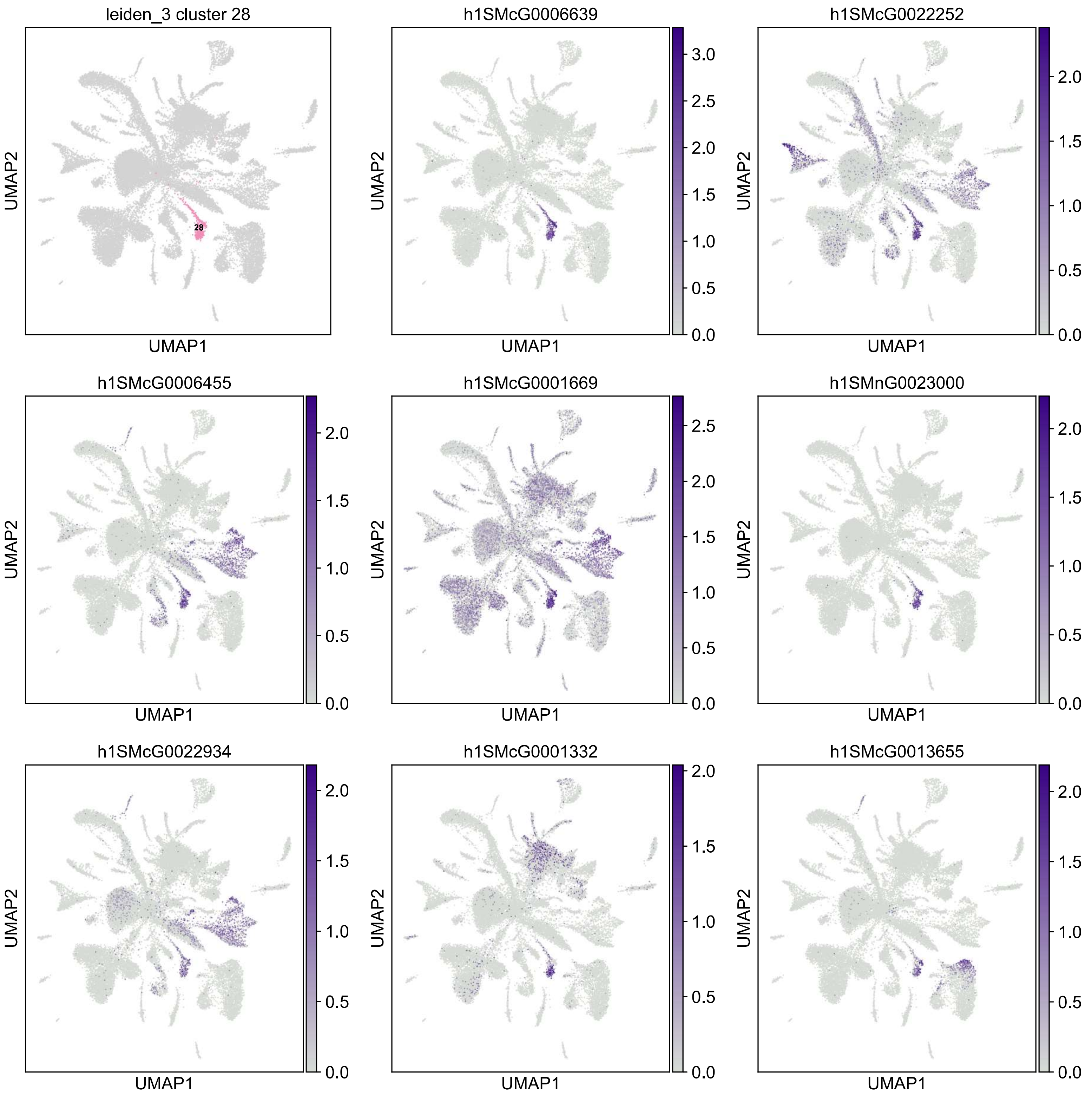
leiden_3 cluster 28
28
h1SMcG0006639
3.0
2.5
2.0
1.5
1.0
0.5
0.0
h1SMcG0022252
2.0
1.5
1.0
0.5
0.0
h1SMcG0006455
2.0
1.5
1.0
0.5
0.0
h1SMcG0001669
2.5
2.0
1.5
1.0
0.5
0.0
h1SMnG0023000
2.0
1.5
1.0
0.5
0.0
h1SMcG0022934
2.0
1.5
1.0
0.5
0.0
h1SMcG0001332
2.0
1.5
1.0
0.5
0.0
h1SMcG0013655
2.0
1.5
1.0
0.5
0.0
UMAP1
UMAP2

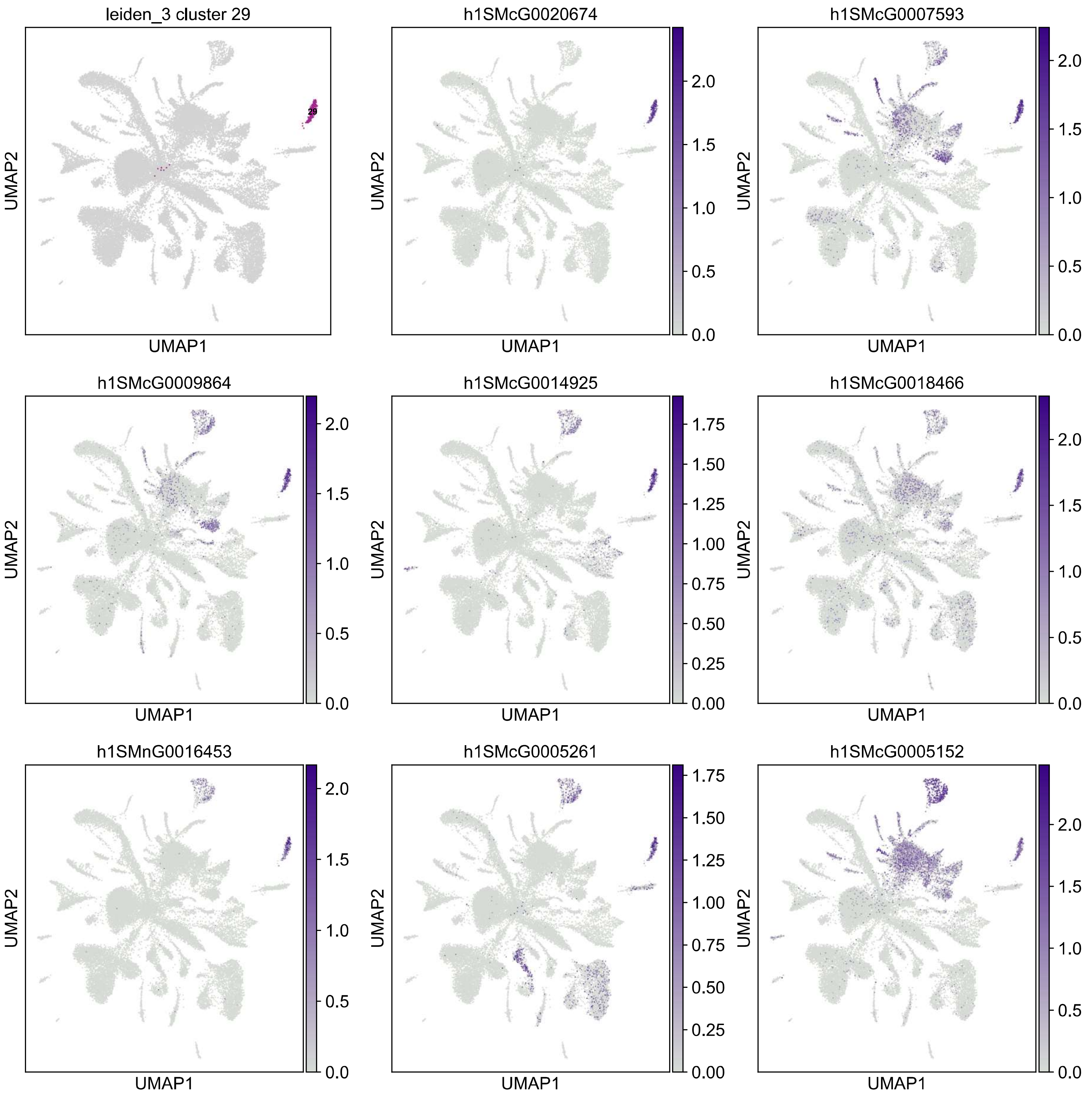
leiden_3 cluster 29
29
h1SMcG0020674
h1SMcG0007593
h1SMcG0009864
h1SMcG0014925
h1SMcG0018466
h1SMnG0016453
h1SMcG0005261
h1SMcG0005152
UMAP1
UMAP2
0.0
0.5
1.0
1.5
2.0
0.00
0.25
0.50
0.75
1.00
1.25
1.50
1.75

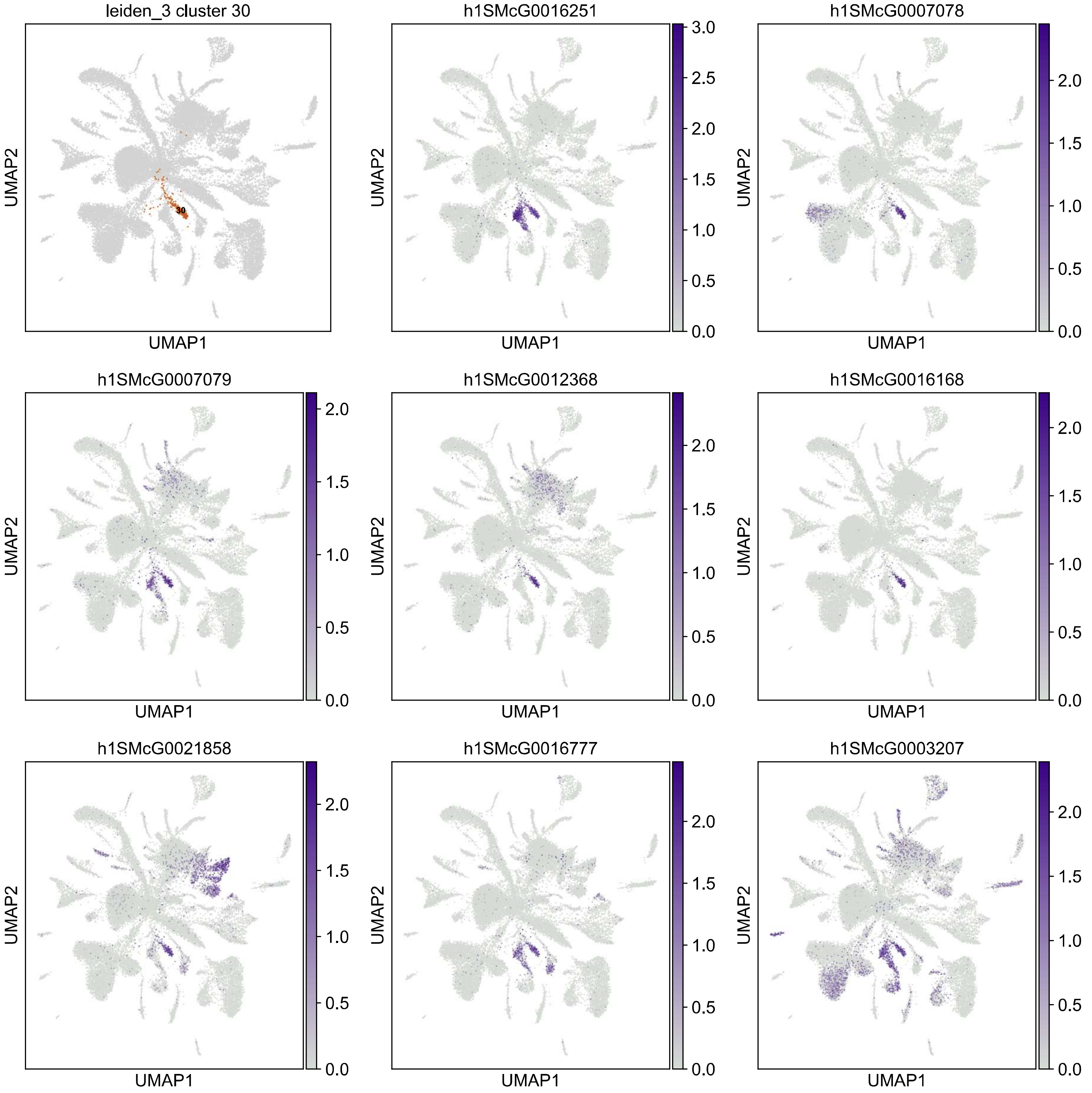
leiden_3 cluster 30
30
h1SMcG0016251
h1SMcG0007078
h1SMcG0007079
h1SMcG0012368
h1SMcG0016168
h1SMcG0021858
h1SMcG0016777
h1SMcG0003207
UMAP1
UMAP2
0.0
0.5
1.0
1.5
2.0
2.5
3.0

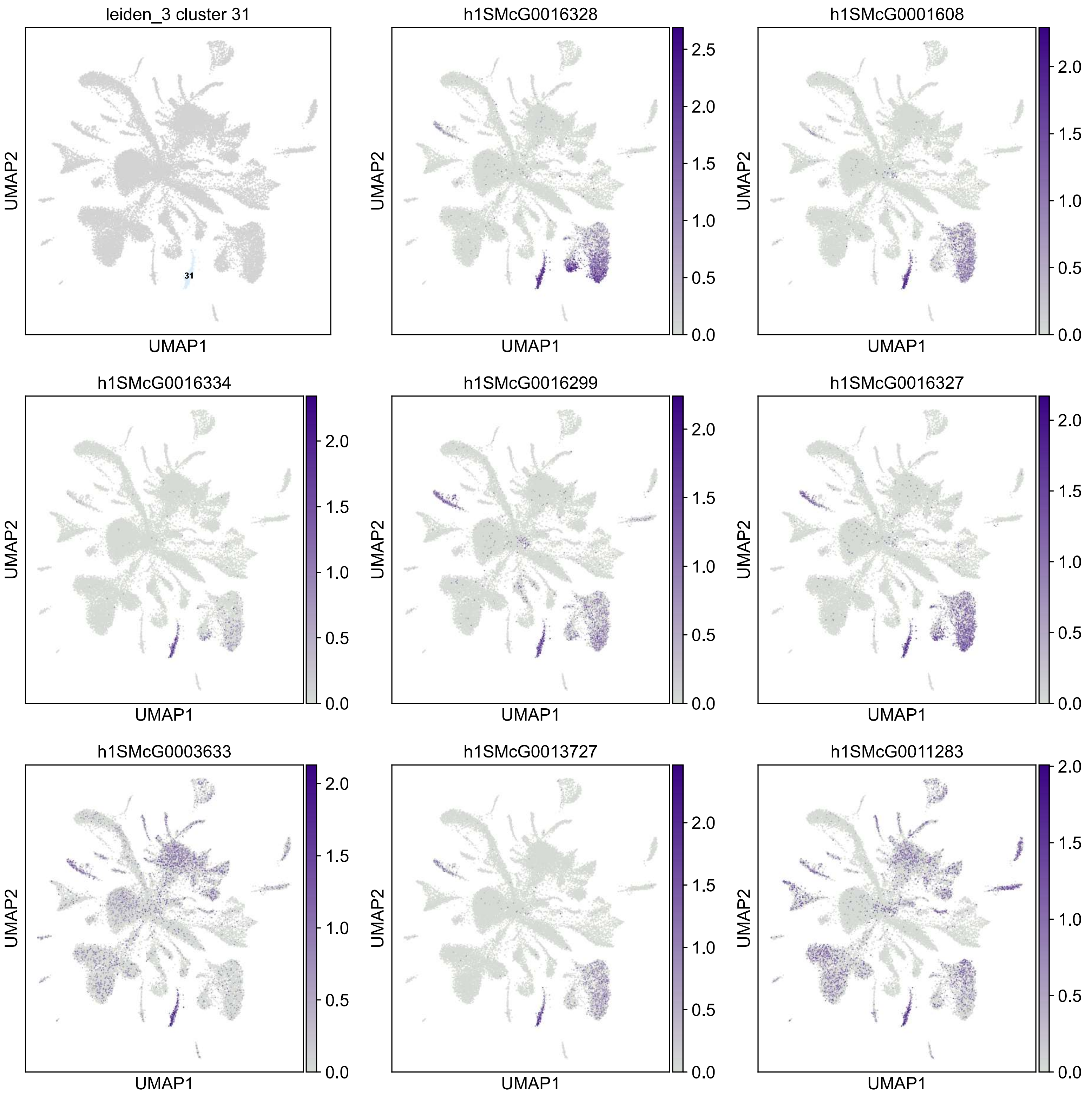
leiden_3 cluster 31
31
h1SMcG0016328
h1SMcG0001608
h1SMcG0016334
h1SMcG0016299
h1SMcG0016327
h1SMcG0003633
h1SMcG0013727
h1SMcG0011283
UMAP1
UMAP2

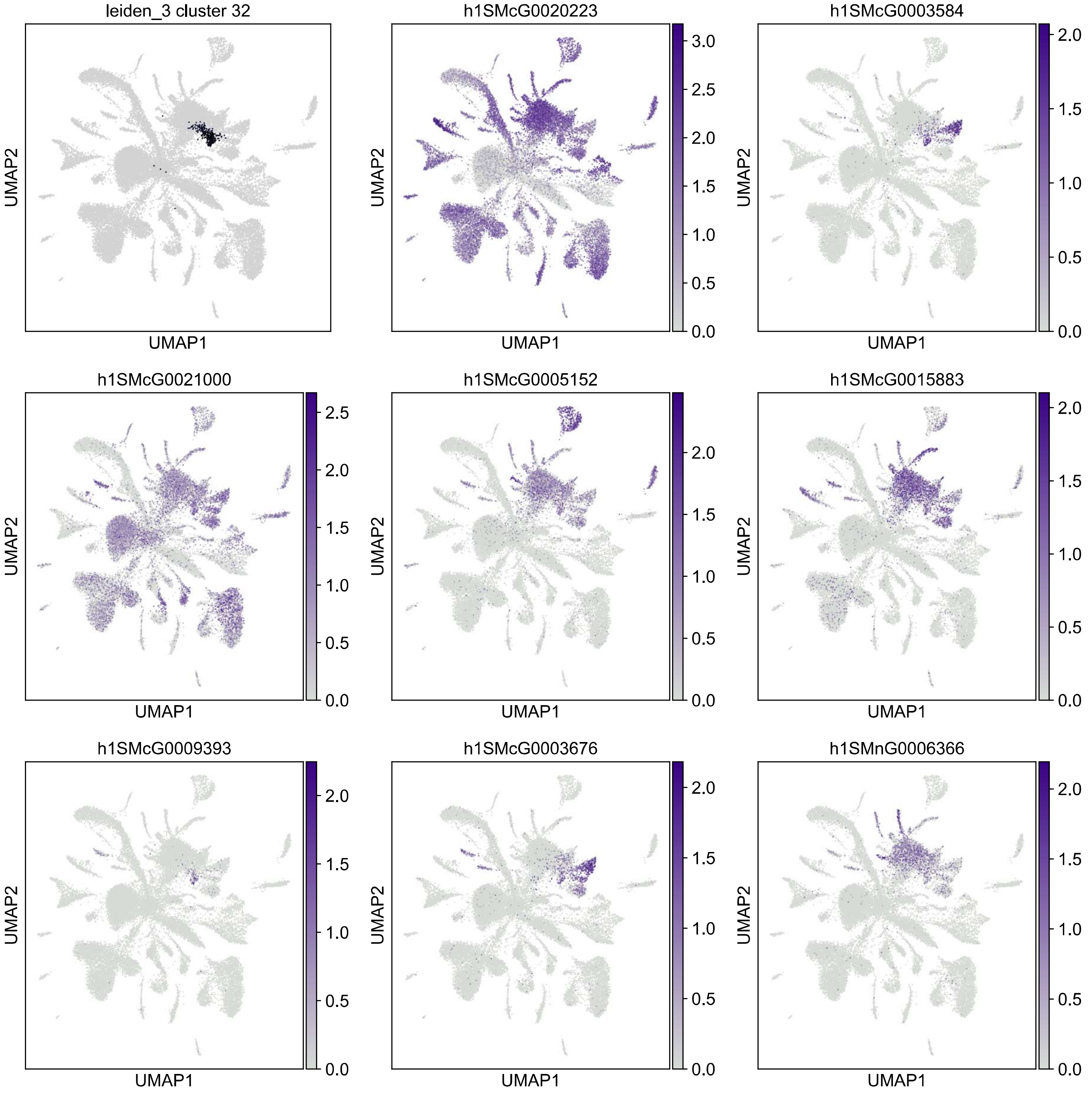

leiden_3 cluster 32
h1SMcG0020223
h1SMcG0003584
h1SMcG0021000
h1SMcG0005152
h1SMcG0015883
h1SMcG0009393
h1SMcG0003676
h1SMnG0006366
UMAP1
UMAP2
0.0
0.5
1.0
1.5
2.0
2.5
3.0

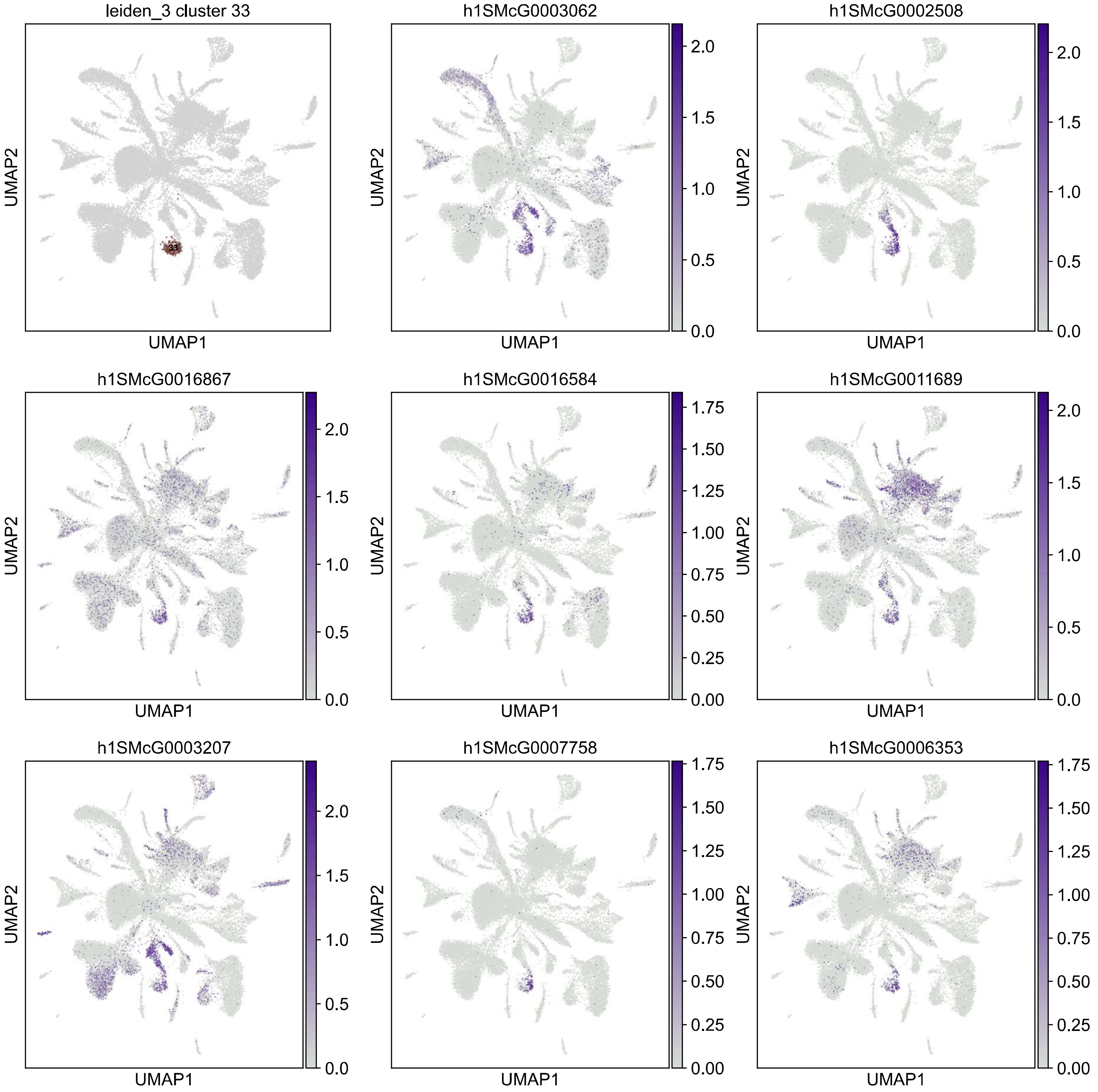

leiden_3 cluster 33
33
h1SMcG0003062
h1SMcG0002508
h1SMcG0016867
h1SMcG0016584
h1SMcG0011689
h1SMcG0003207
h1SMcG0007758
h1SMcG0006353
UMAP1
UMAP2

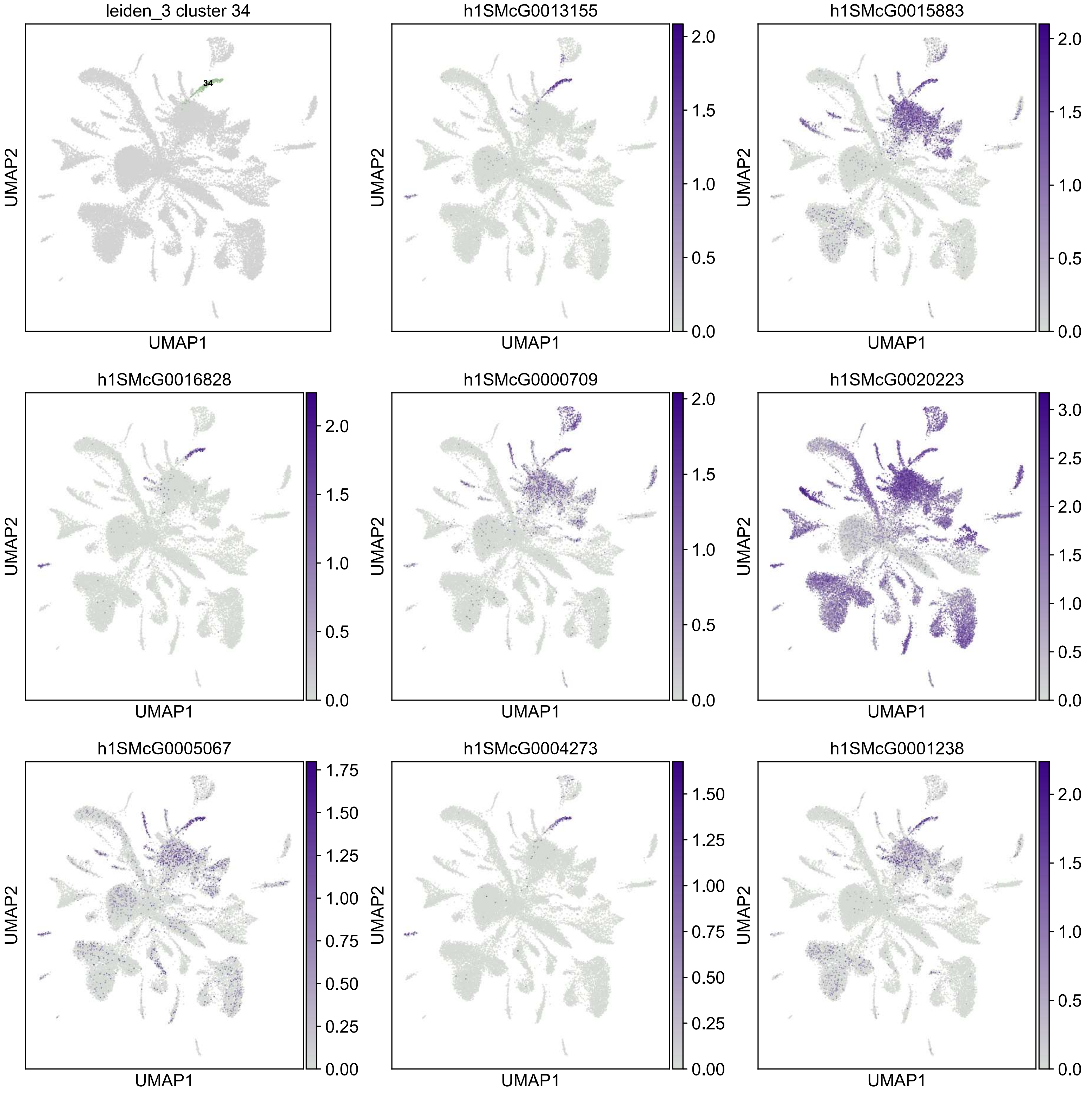

leiden_3 cluster 34
34
h1SMcG0013155
h1SMcG0015883
h1SMcG0016828
h1SMcG0000709
h1SMcG0020223
h1SMcG0005067
h1SMcG0004273
h1SMcG0001238
UMAP1
UMAP2

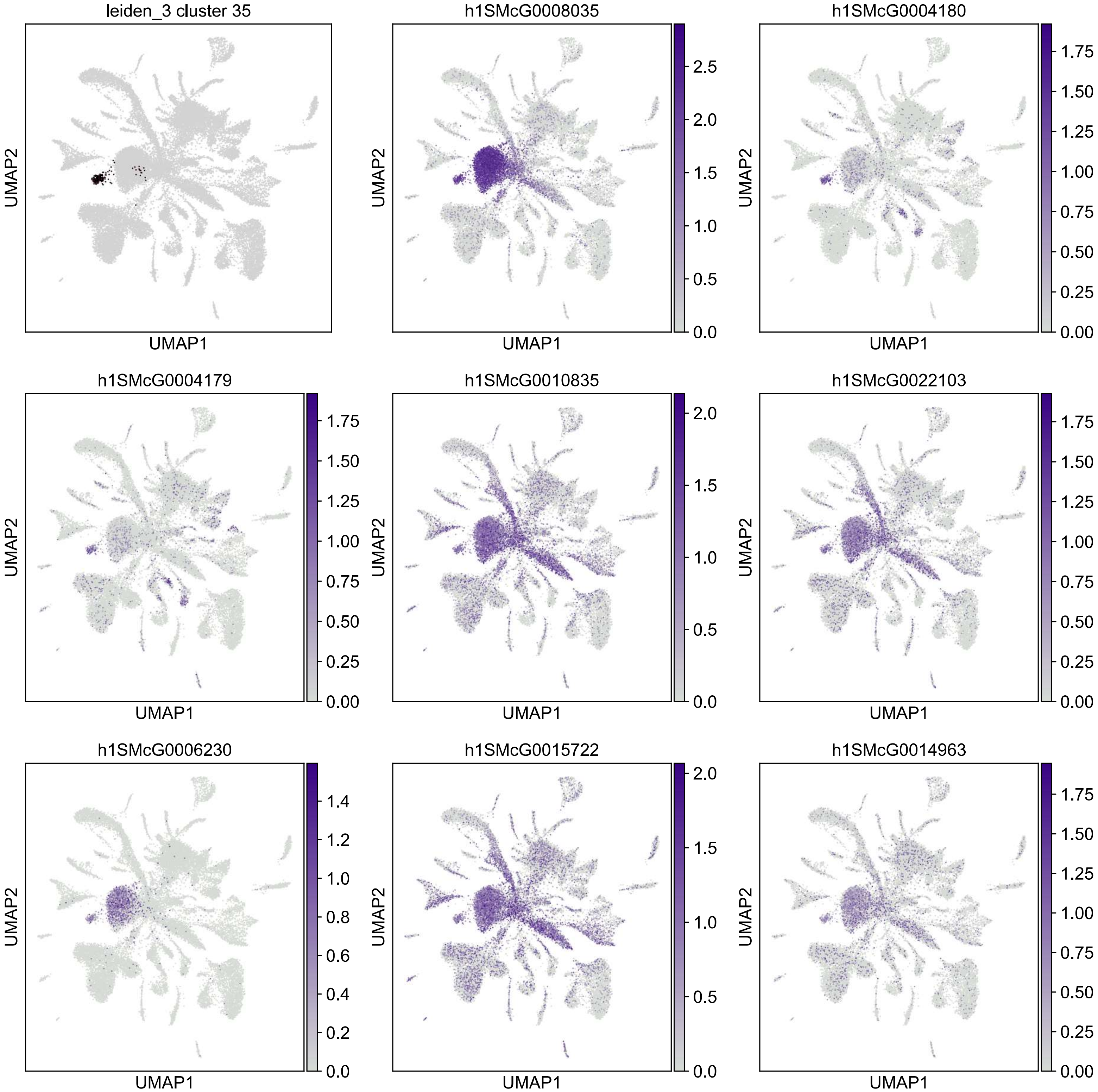
leiden_3 cluster 35
h1SMcG0008035
h1SMcG0004180
h1SMcG0004179
h1SMcG0010835
h1SMcG0022103
h1SMcG0006230
h1SMcG0015722
h1SMcG0014963
UMAP1
UMAP2
0.0 0.5 1.0 1.5 2.0 2.5
0.00 0.25 0.50 0.75 1.00 1.25 1.50 1.75
0.0 0.5 1.0 1.5 2.0
0.0 0.2 0.4 0.6 0.8 1.0 1.2 1.4

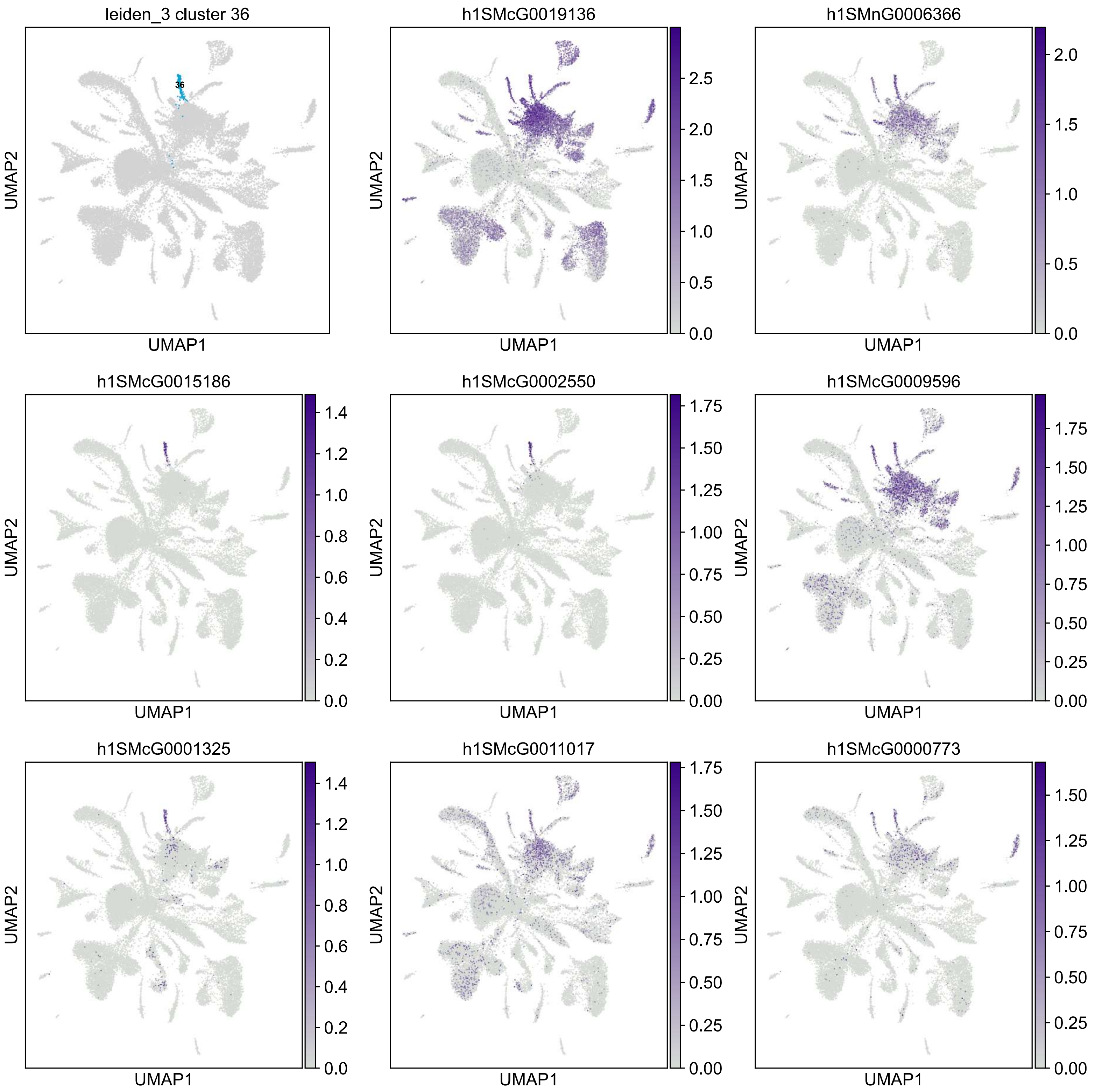
leiden_3 cluster 36
36
h1SMcG0019136
h1SMnG0006366
h1SMcG0015186
h1SMcG0002550
h1SMcG0009596
h1SMcG0001325
h1SMcG0011017
h1SMcG0000773
UMAP1
UMAP2

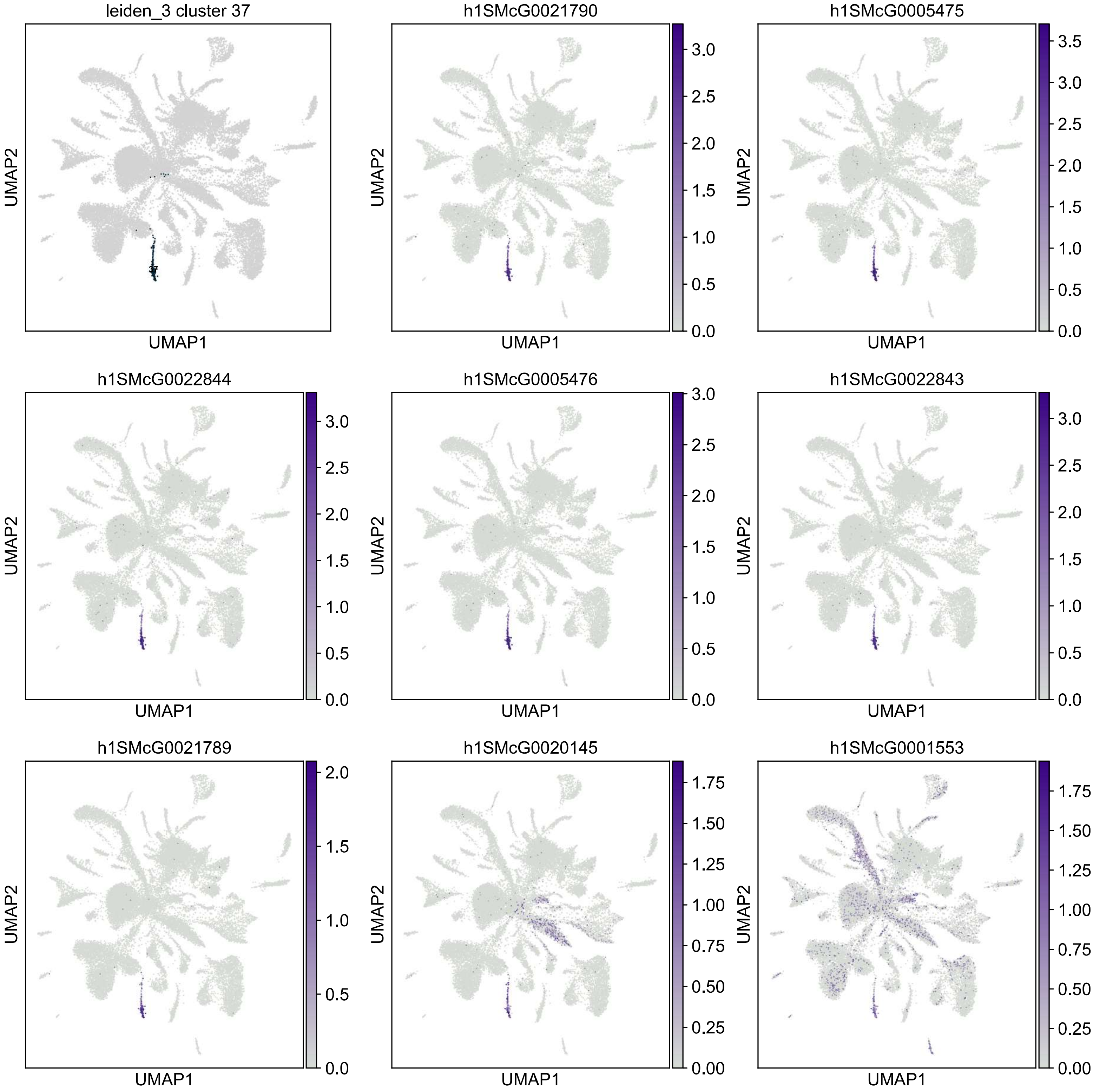

leiden_3 cluster 37
h1SMcG0021790
h1SMcG0005475
h1SMcG0022844
h1SMcG0005476
h1SMcG0022843
h1SMcG0021789
h1SMcG0020145
h1SMcG0001553
UMAP1
UMAP2

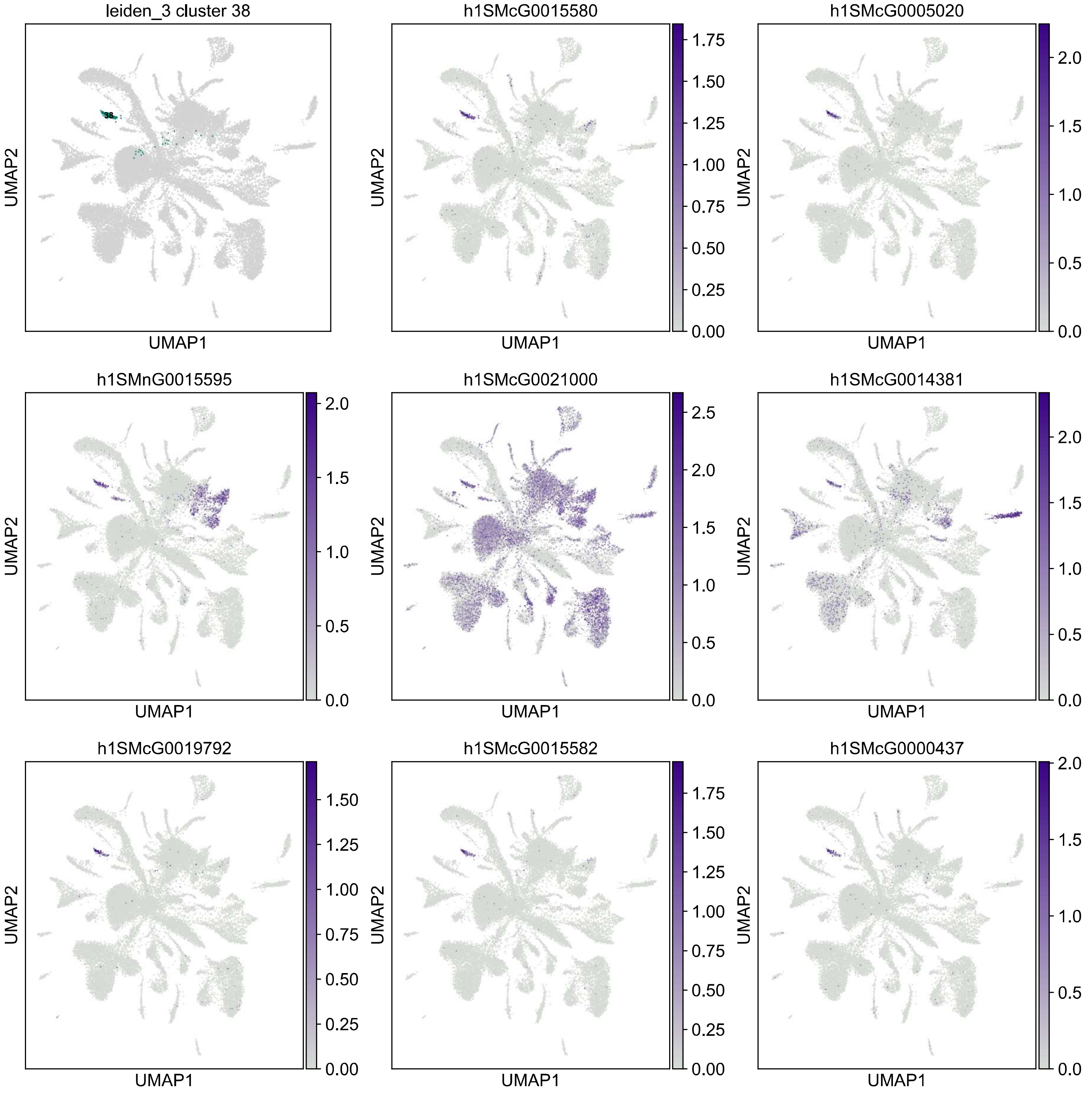
leiden_3 cluster 38
38
h1SMcG0015580
h1SMcG0005020
h1SMnG0015595
h1SMcG0021000
h1SMcG0014381
h1SMcG0019792
h1SMcG0015582
h1SMcG0000437
UMAP1
UMAP2

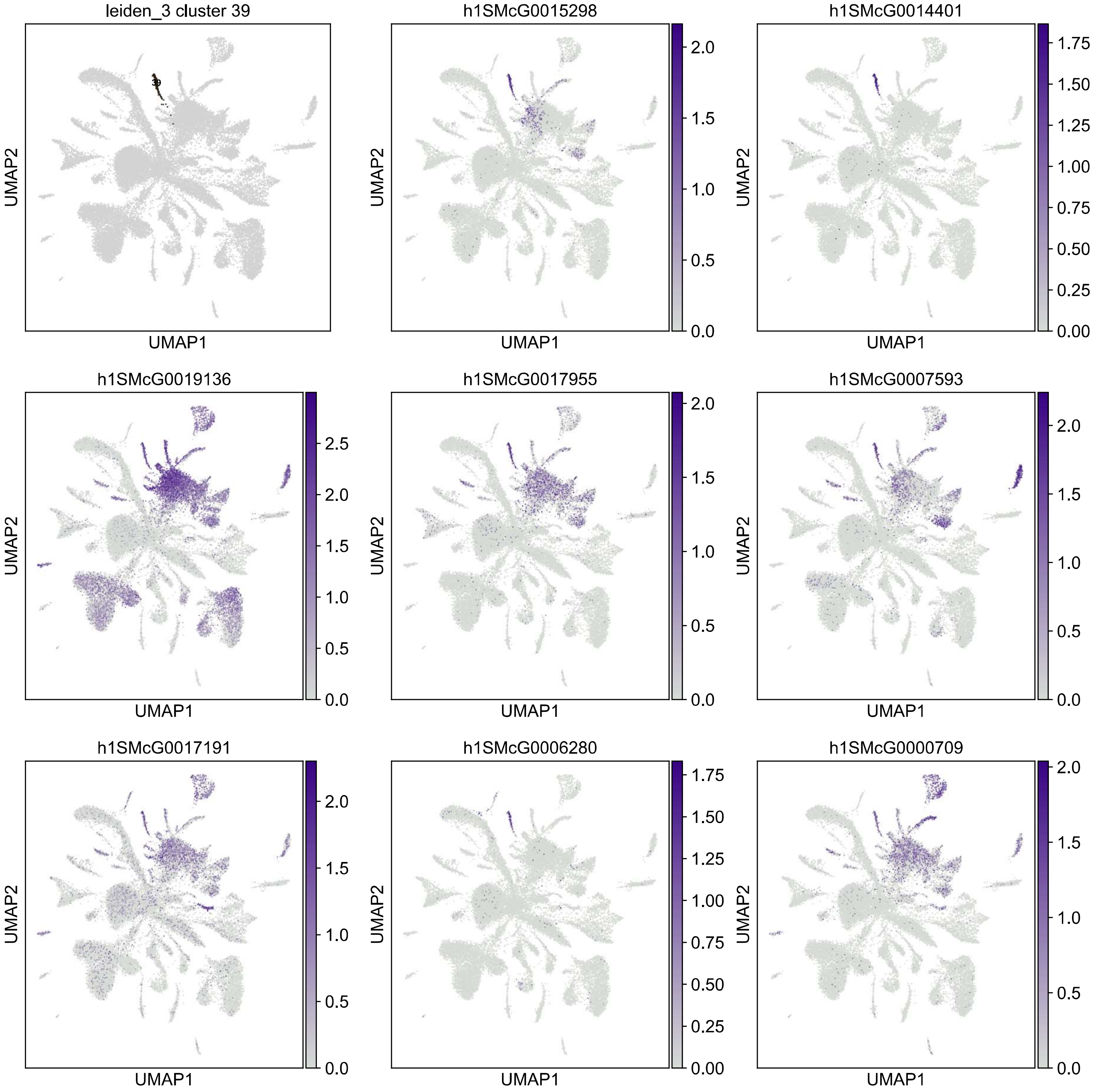
leiden_3 cluster 39
39
h1SMcG0015298
h1SMcG0014401
h1SMcG0019136
h1SMcG0017955
h1SMcG0007593
h1SMcG0017191
h1SMcG0006280
h1SMcG0000709
UMAP1
UMAP2

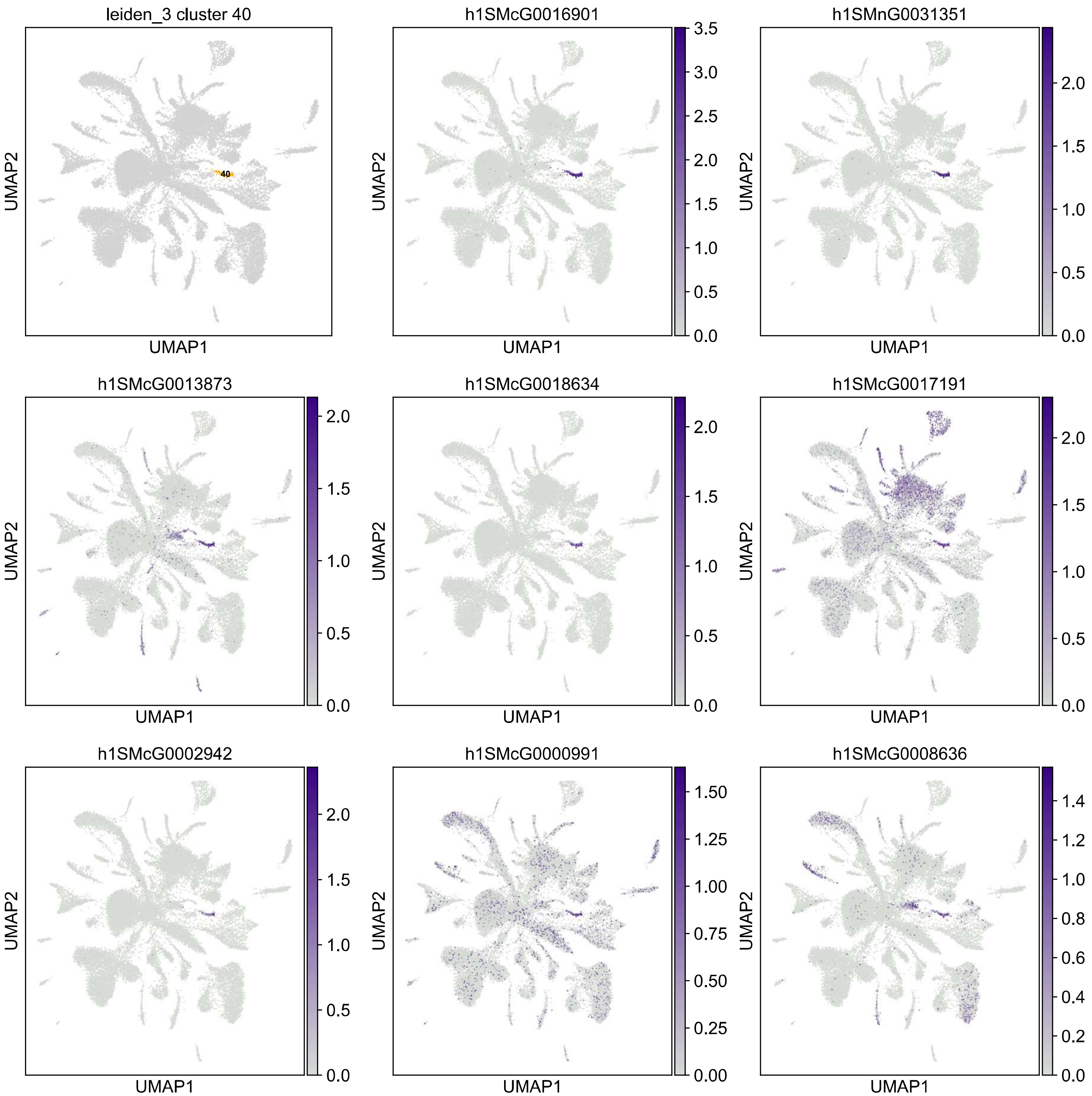

leiden_3 cluster 40
40
h1SMcG0016901
h1SMnG0031351
h1SMcG0013873
h1SMcG0018634
h1SMcG0017191
h1SMcG0002942
h1SMcG0000991
h1SMcG0008636
UMAP1
UMAP2

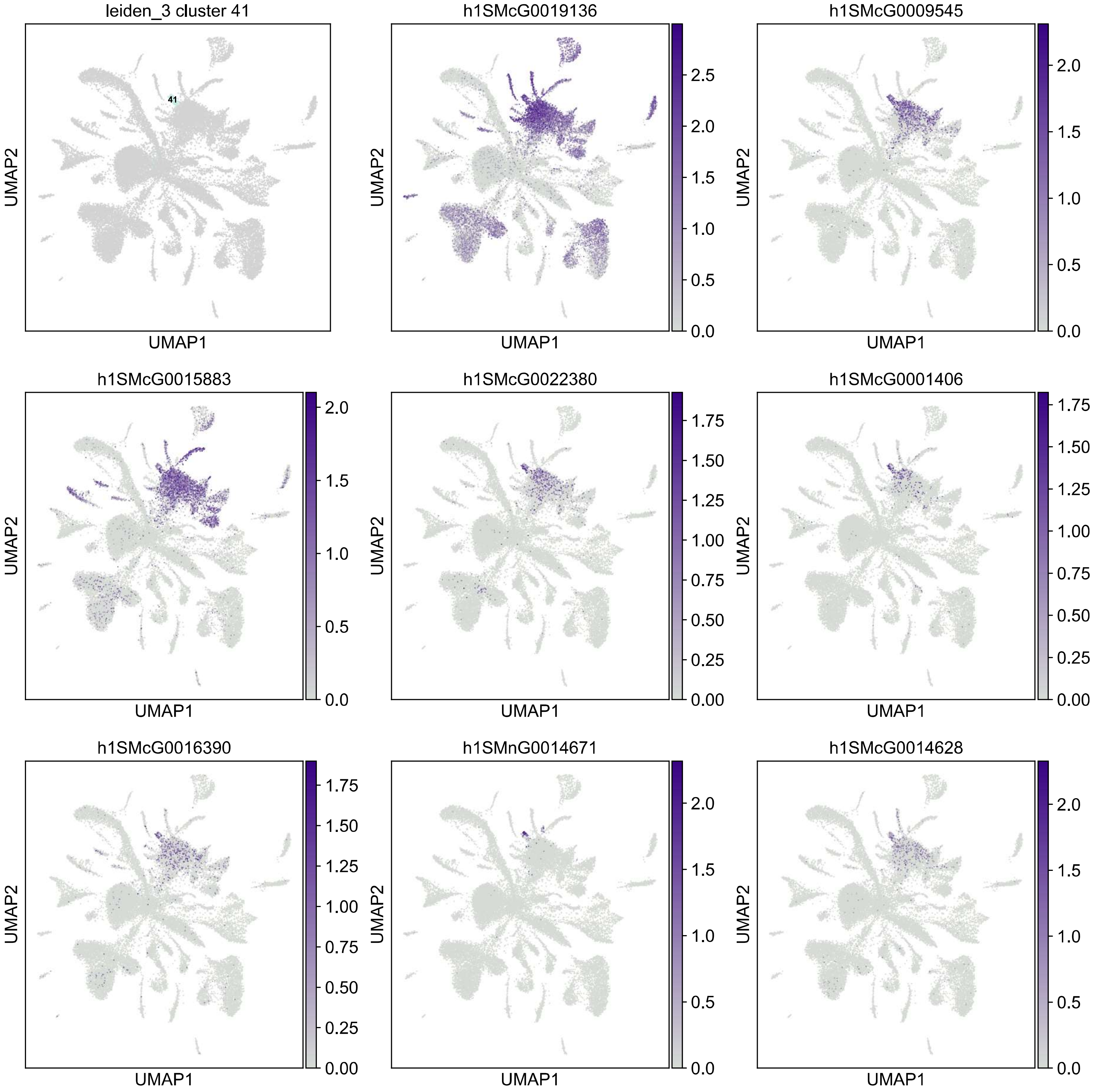
leiden_3 cluster 41
41
h1SMcG0019136
h1SMcG0009545
h1SMcG0015883
h1SMcG0022380
h1SMcG0001406
h1SMcG0016390
h1SMnG0014671
h1SMcG0014628
UMAP1
UMAP2

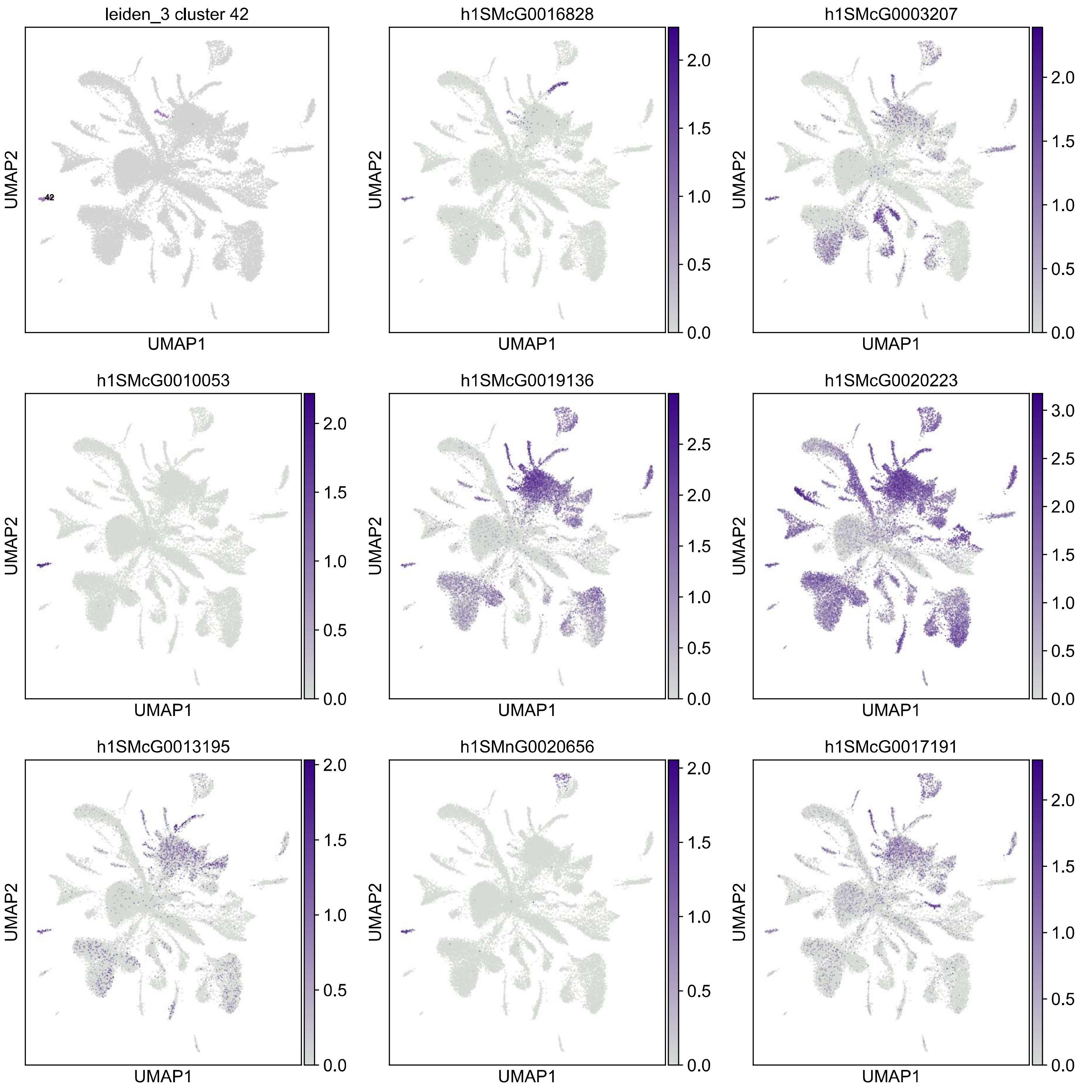

leiden_3 cluster 42
42
h1SMcG0016828
h1SMcG0003207
h1SMcG0010053
h1SMcG0019136
h1SMcG0020223
h1SMcG0013195
h1SMnG0020656
h1SMcG0017191
UMAP1
UMAP2

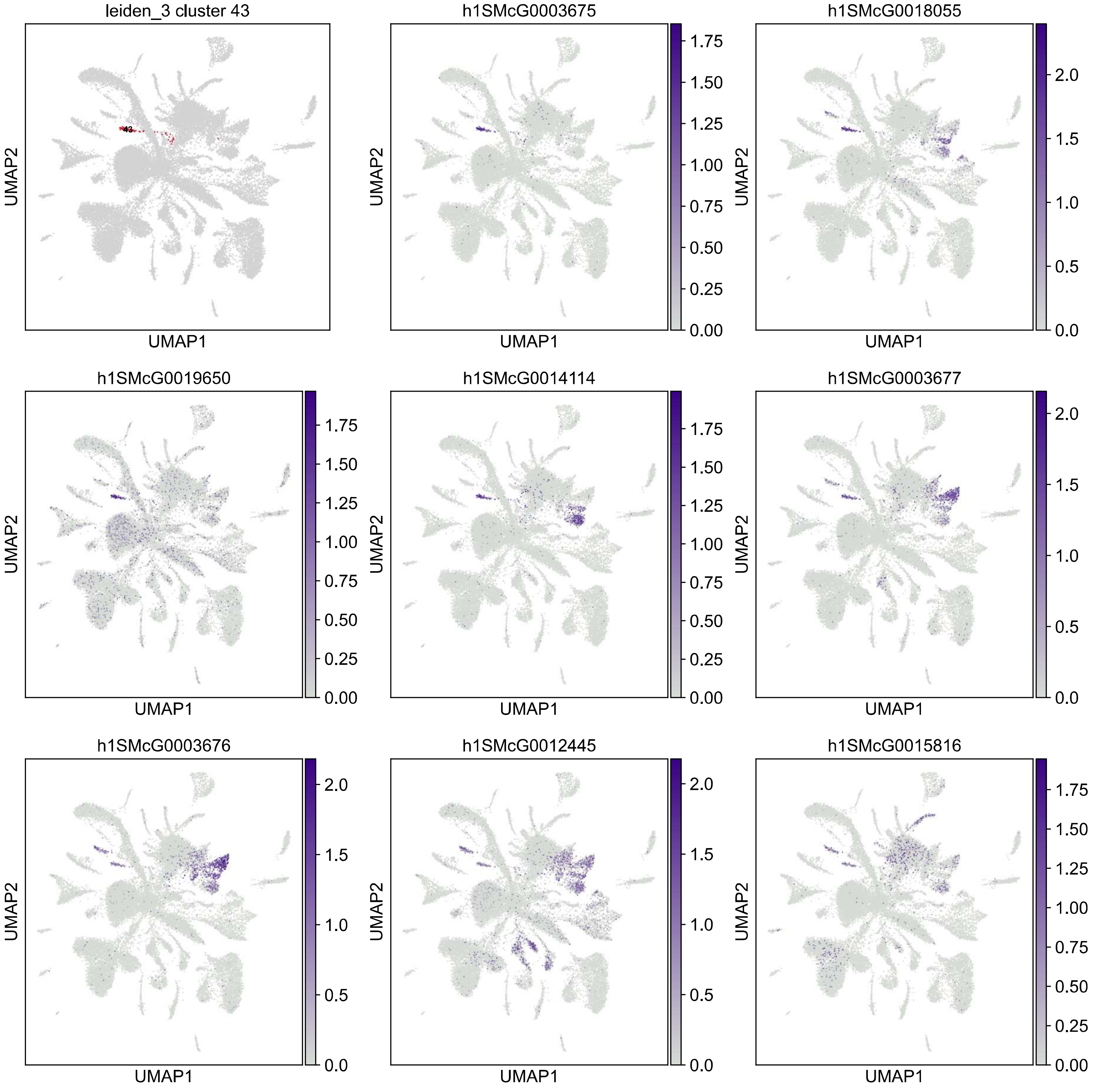
leiden_3 cluster 43
43
h1SMcG0003675
h1SMcG0018055
h1SMcG0019650
h1SMcG0014114
h1SMcG0003677
h1SMcG0003676
h1SMcG0012445
h1SMcG0015816
UMAP1
UMAP2
0.00
0.25
0.50
0.75
1.00
1.25
1.50
1.75
0.0
0.5
1.0
1.5
2.0

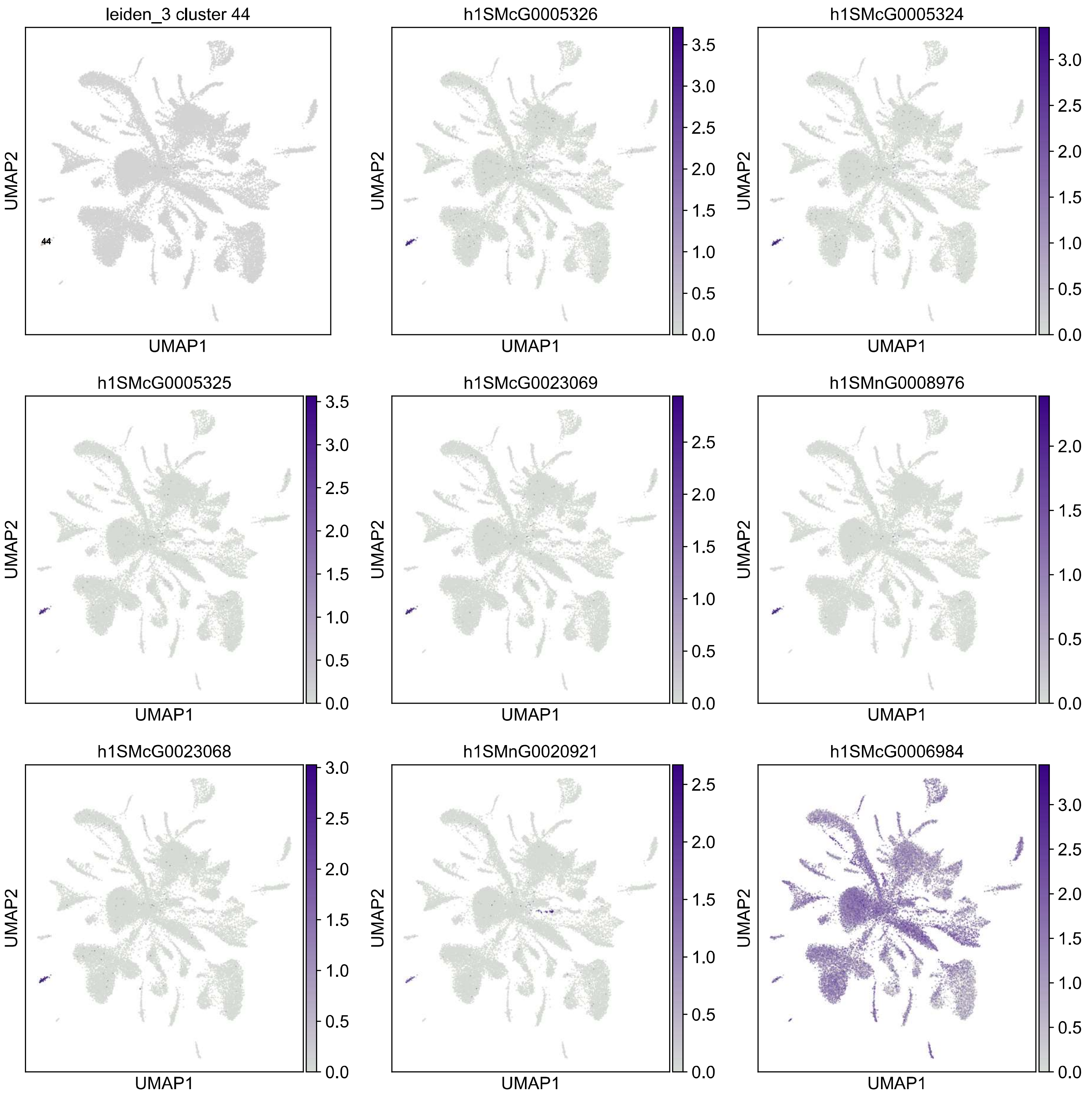

leiden_3 cluster 44
44
h1SMcG0005326
h1SMcG0005324
h1SMcG0005325
h1SMcG0023069
h1SMnG0008976
h1SMcG0023068
h1SMnG0020921
h1SMcG0006984
UMAP1
UMAP2

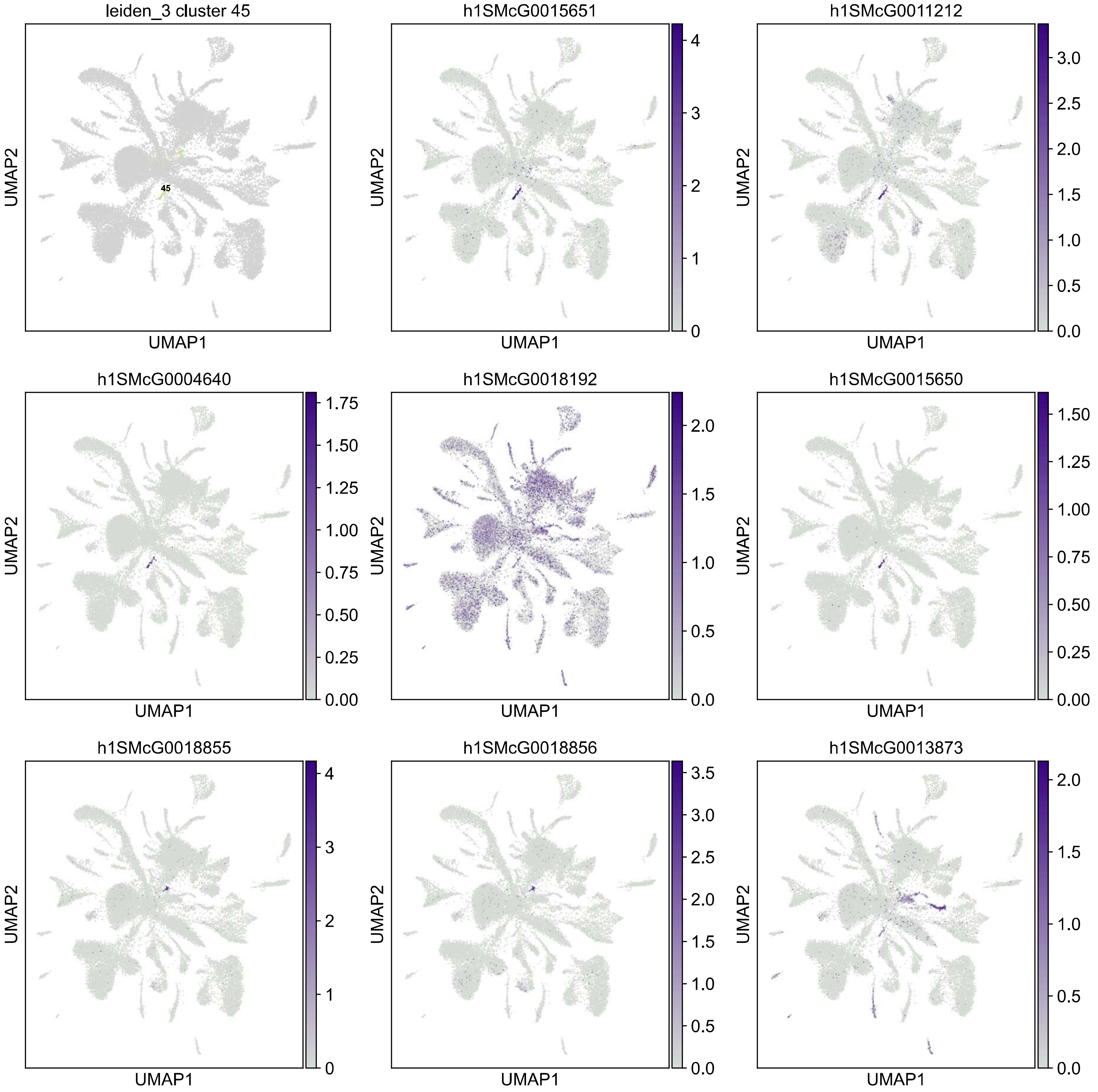

leiden_3 cluster 45
45
h1SMcG0015651
h1SMcG0011212
h1SMcG0004640
h1SMcG0018192
h1SMcG0015650
h1SMcG0018855
h1SMcG0018856
h1SMcG0013873
UMAP1
UMAP2

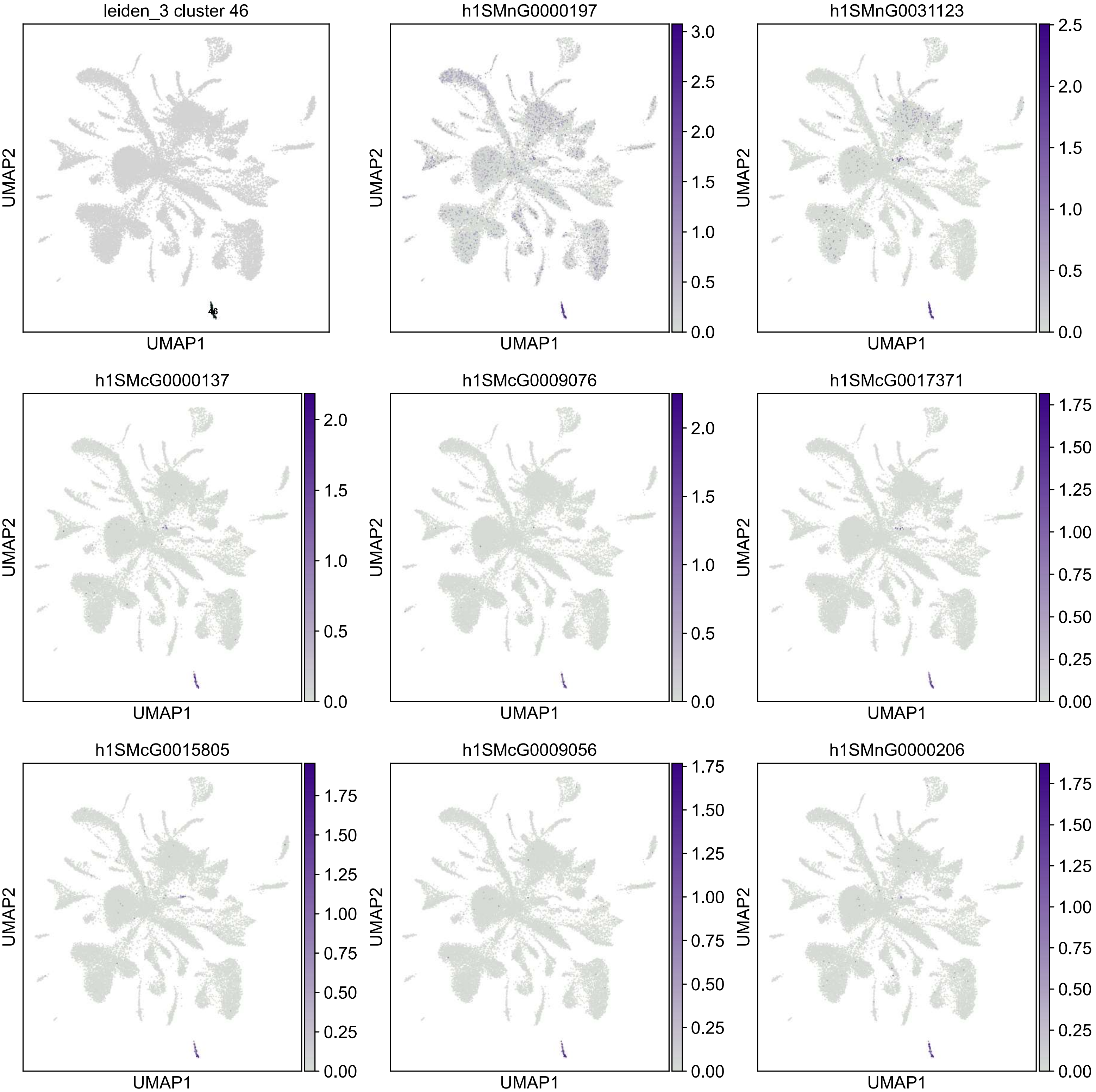

leiden_3 cluster 46
46
h1SMnG0000197
h1SMnG0031123
h1SMcG0000137
h1SMcG0009076
h1SMcG0017371
h1SMcG0015805
h1SMcG0009056
h1SMnG0000206
UMAP1
UMAP2
0.0
0.5
1.0
1.5
2.0
2.5
3.0
0.00
0.25
0.50
0.75
1.00
1.25
1.50
1.75

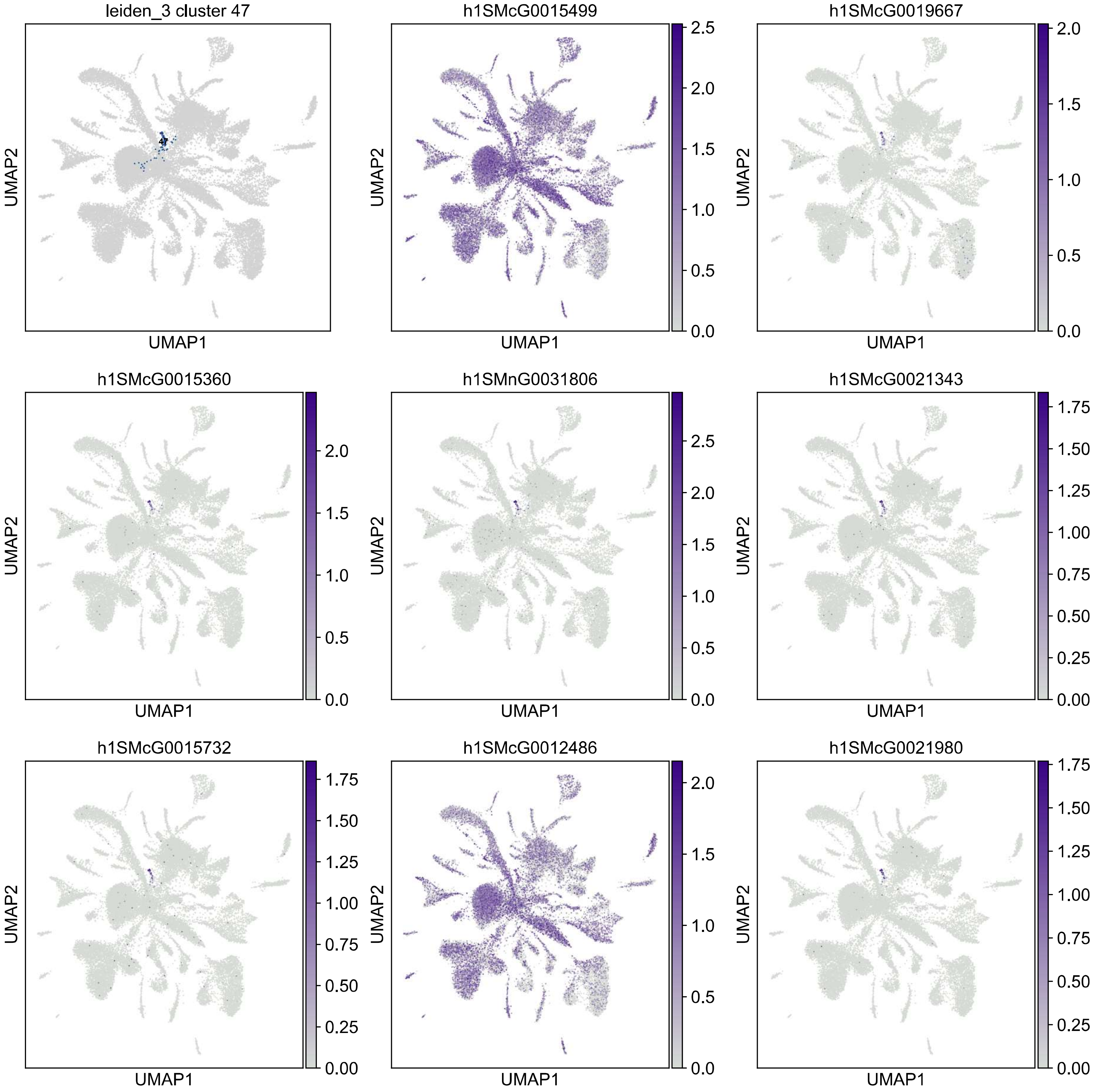

leiden_3 cluster 47
47
h1SMcG0015499
h1SMcG0019667
h1SMcG0015360
h1SMnG0031806
h1SMcG0021343
h1SMcG0015732
h1SMcG0012486
h1SMcG0021980
UMAP1
UMAP2

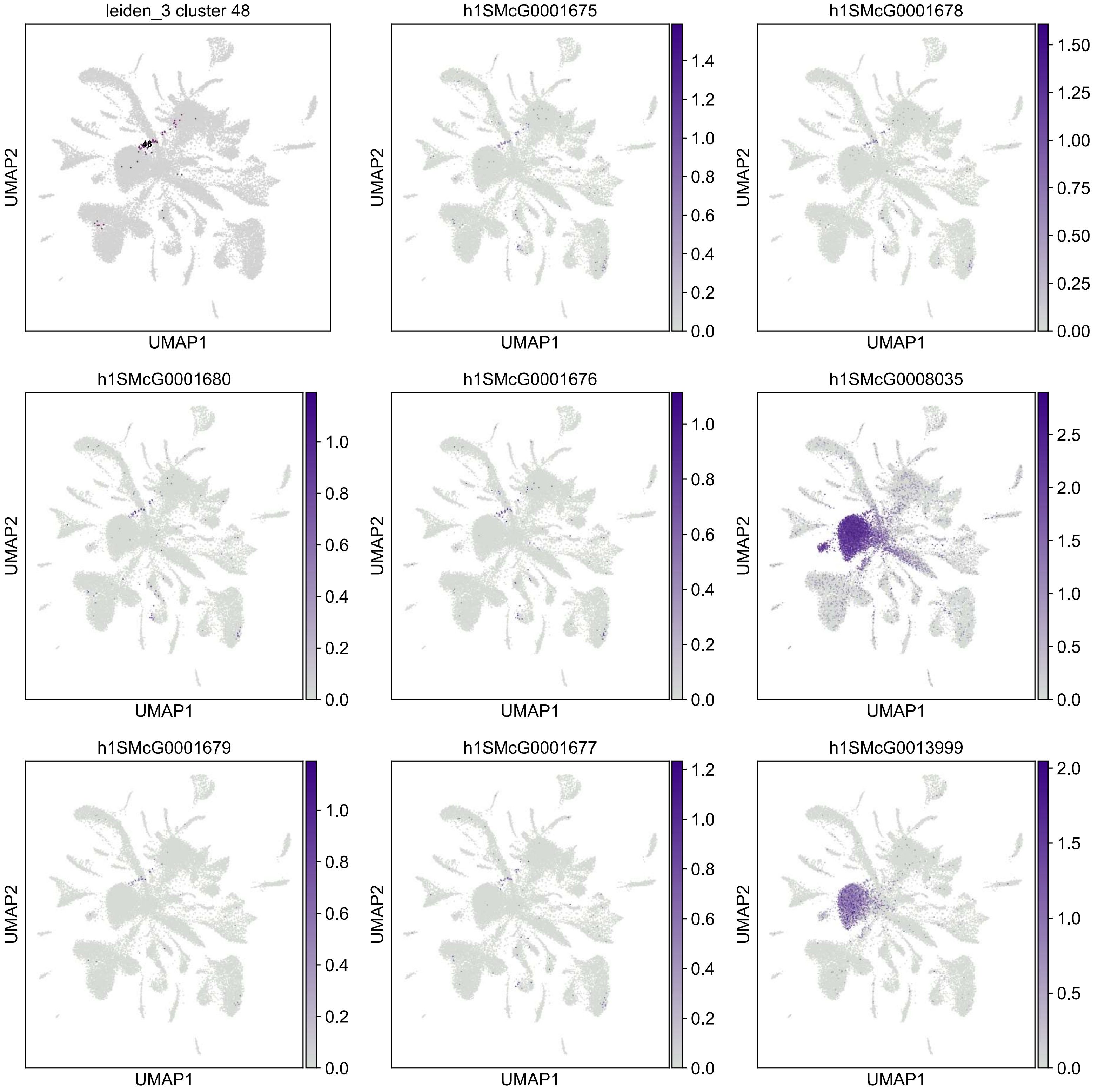

leiden_3 cluster 48
48
h1SMcG0001675
h1SMcG0001678
h1SMcG0001680
h1SMcG0001676
h1SMcG0008035
h1SMcG0001679
h1SMcG0001677
h1SMcG0013999
UMAP1
UMAP2

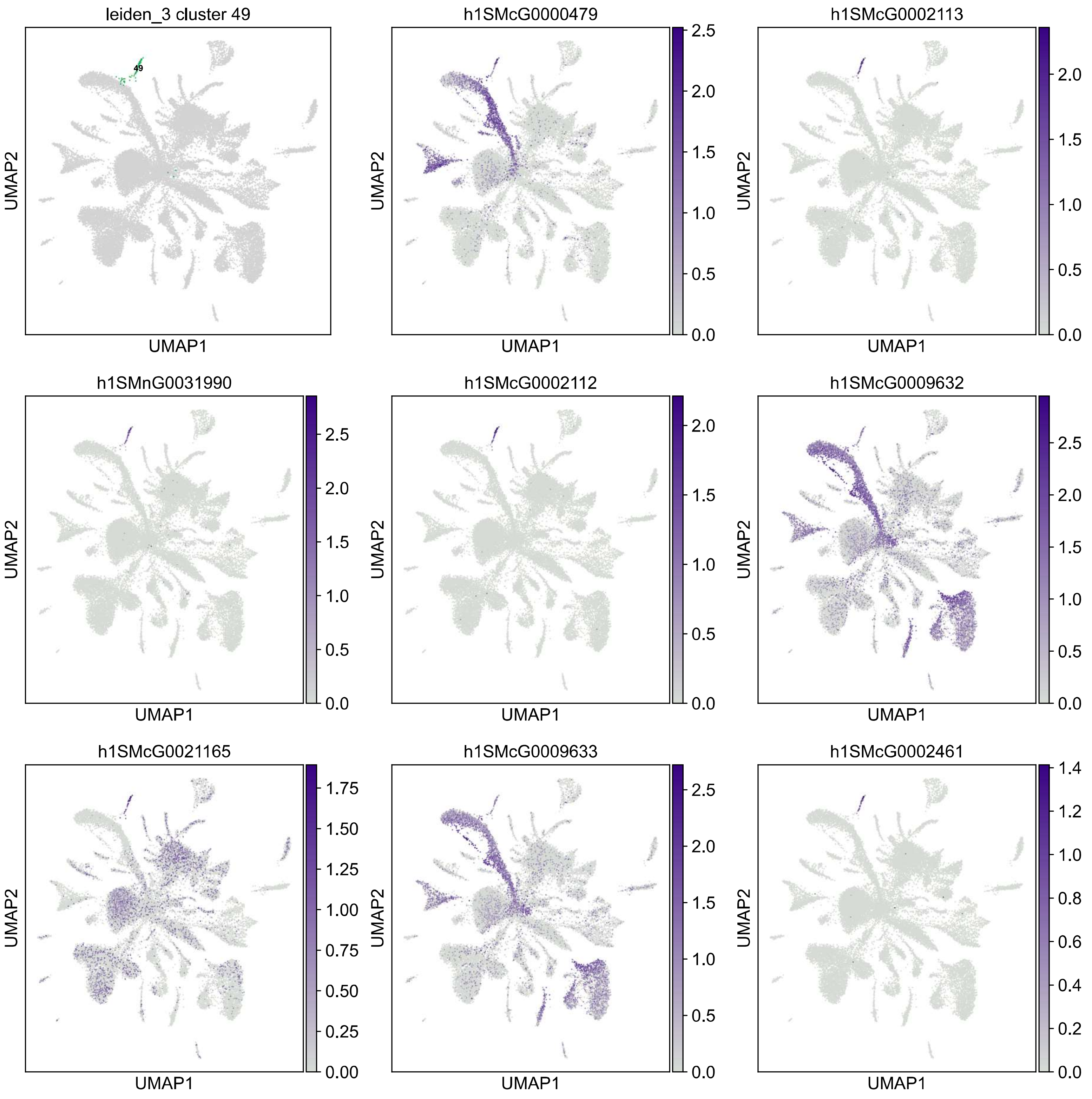
leiden_3 cluster 49
49
h1SMcG0000479
h1SMcG0002113
h1SMnG0031990
h1SMcG0002112
h1SMcG0009632
h1SMcG0021165
h1SMcG0009633
h1SMcG0002461
UMAP1
UMAP2

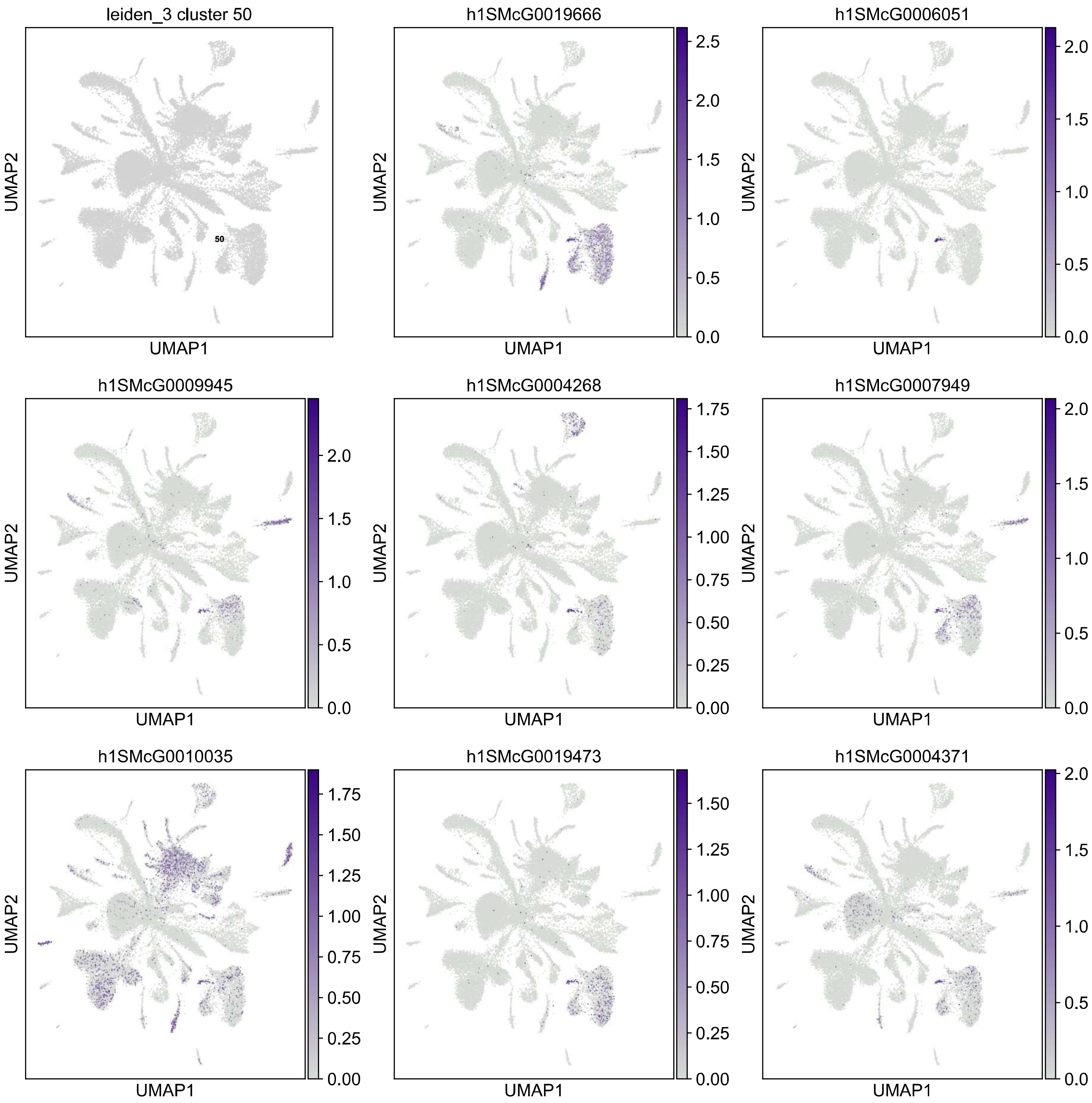
leiden_3 cluster 50
50
h1SMcG0019666
h1SMcG0006051
h1SMcG0009945
h1SMcG0004268
h1SMcG0007949
h1SMcG0010035
h1SMcG0019473
h1SMcG0004371
UMAP1
UMAP2
0.0
0.5
1.0
1.5
2.0
2.5
0.00
0.25
0.50
0.75
1.00
1.25
1.50
1.75

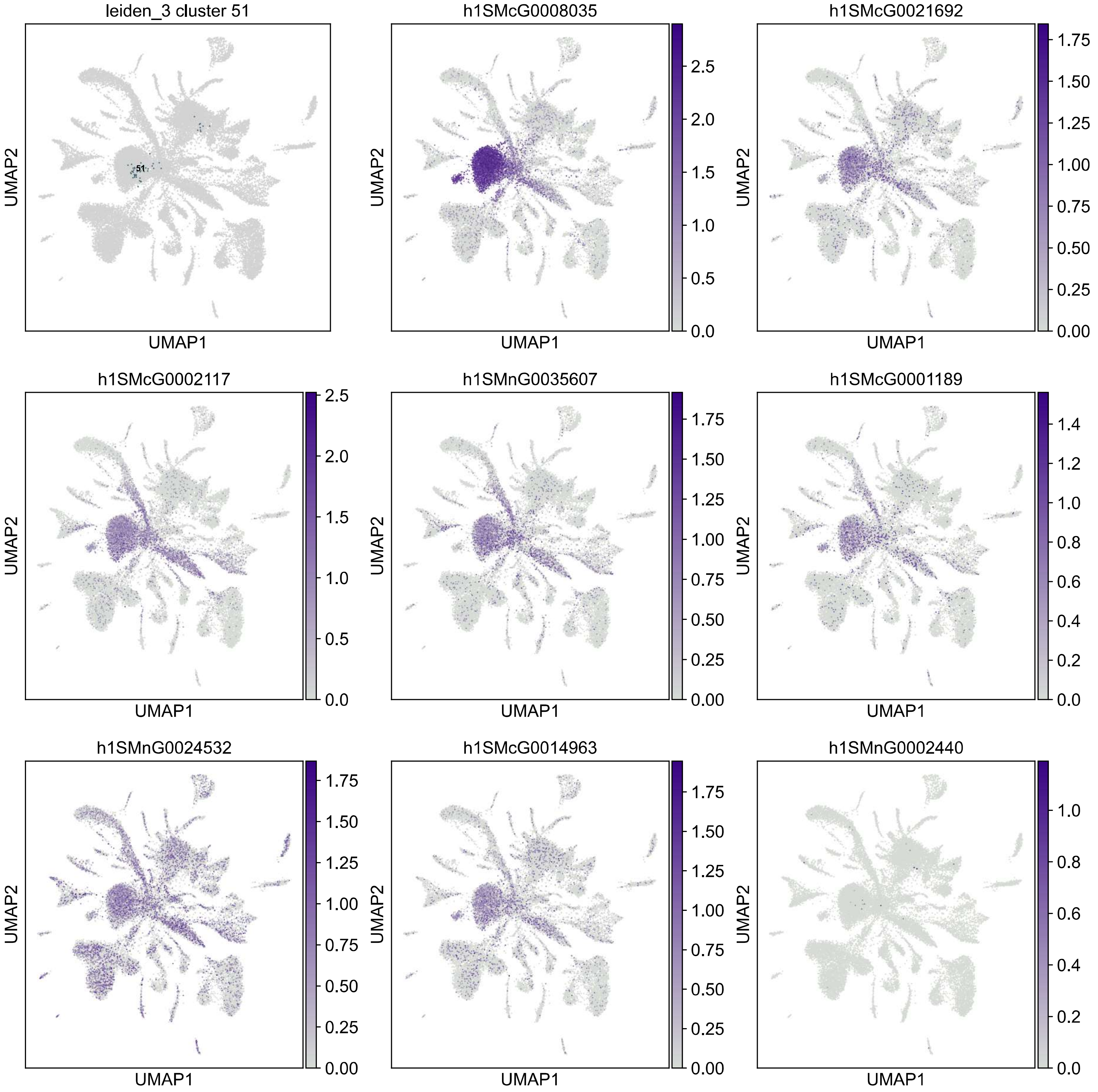

leiden_3 cluster 51
51
h1SMcG0008035
h1SMcG0021692
h1SMcG0002117
h1SMnG0035607
h1SMcG0001189
h1SMnG0024532
h1SMcG0014963
h1SMnG0002440
UMAP1
UMAP2

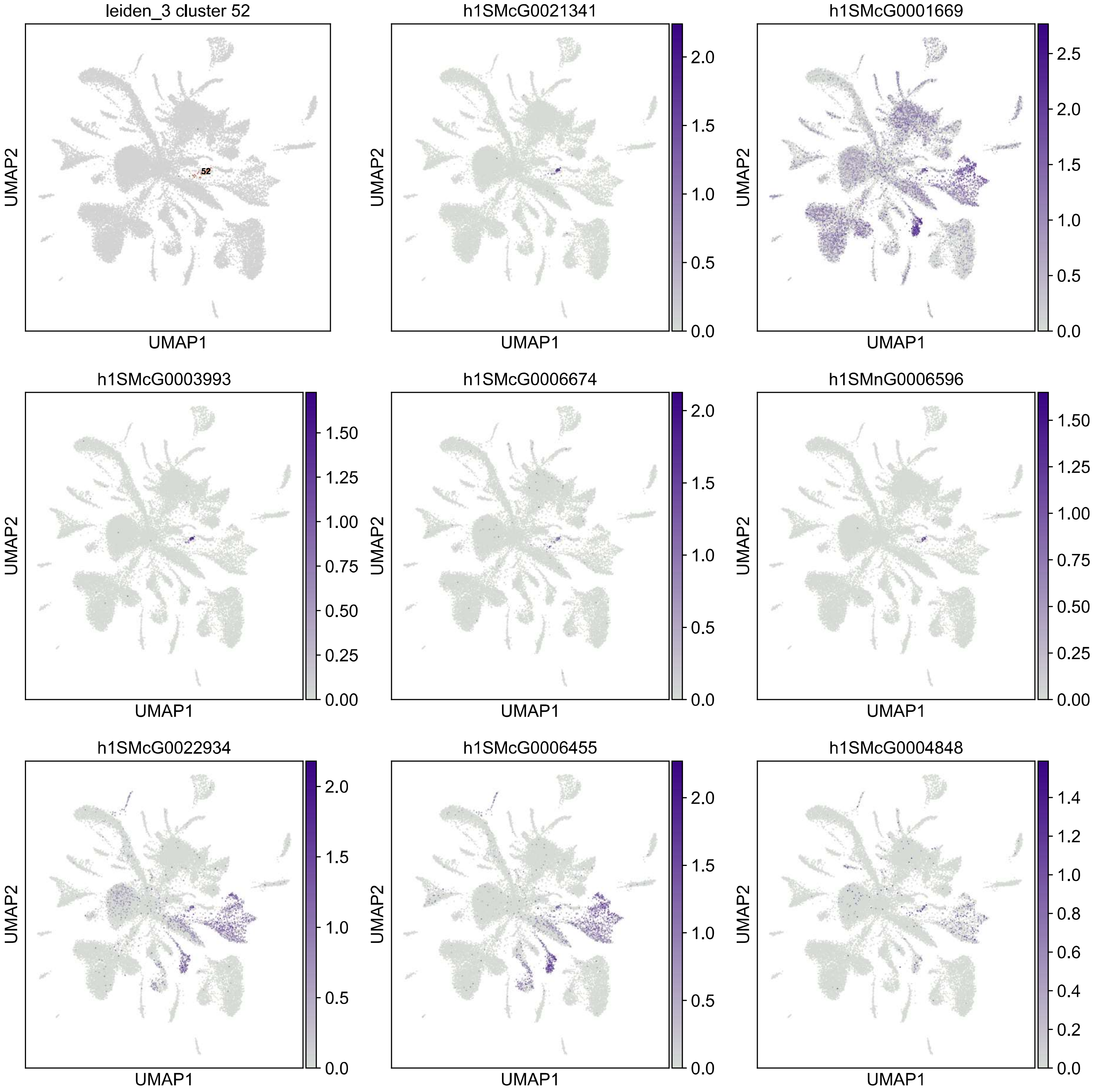
leiden_3 cluster 52
52
h1SMcG0021341
h1SMcG0001669
h1SMcG0003993
h1SMcG0006674
h1SMnG0006596
h1SMcG0022934
h1SMcG0006455
h1SMcG0004848
UMAP1
UMAP2

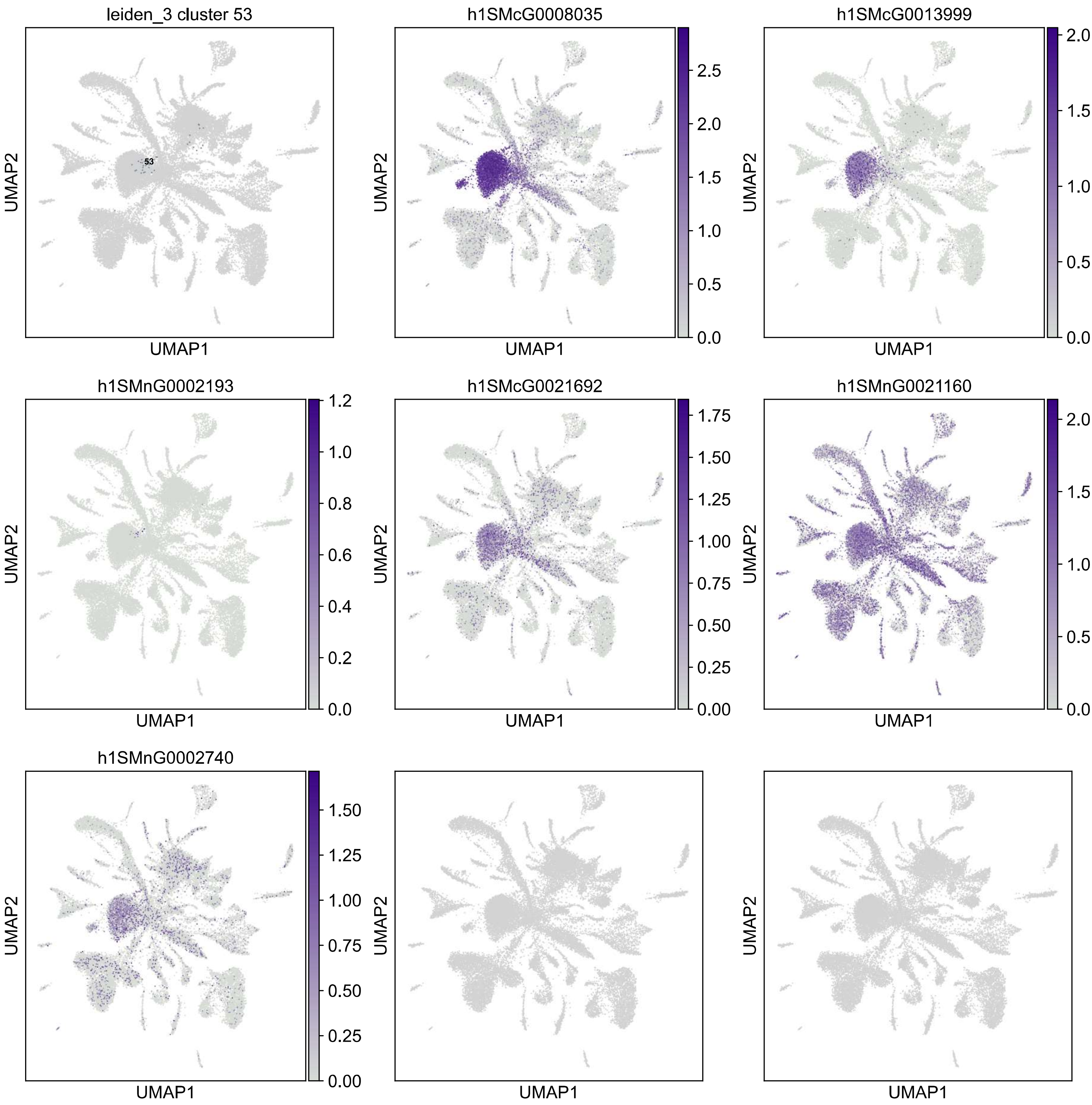

leiden_3 cluster 53
53
h1SMcG0008035
h1SMcG0013999
h1SMnG0002193
h1SMcG0021692
h1SMnG0021160
h1SMnG0002740
UMAP1
UMAP2

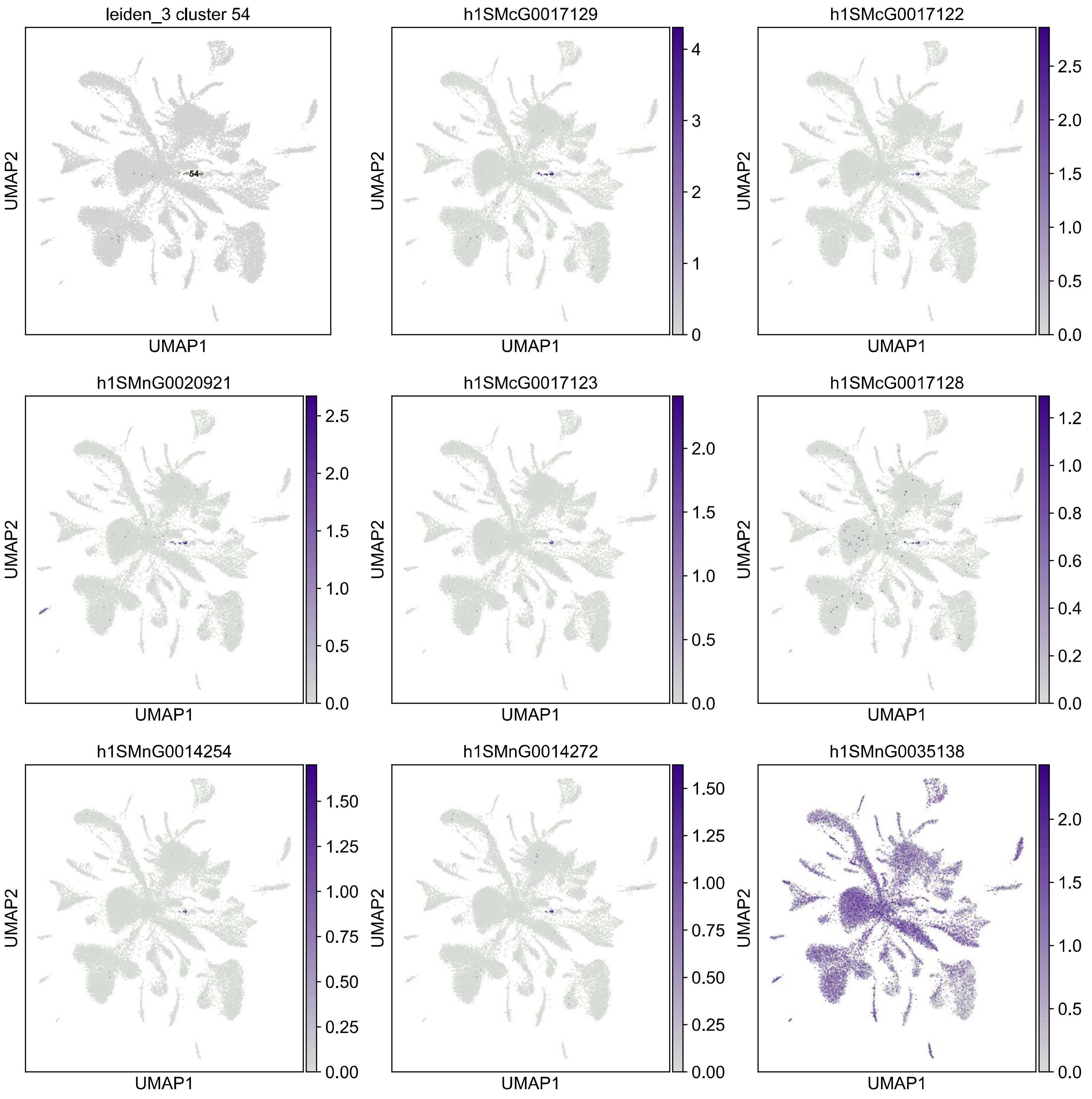

leiden_3 cluster 54
54
UMAP1
UMAP2
h1SMcG0017129
h1SMcG0017122
h1SMnG0020921
h1SMcG0017123
h1SMcG0017128
h1SMnG0014254
h1SMnG0014272
h1SMnG0035138

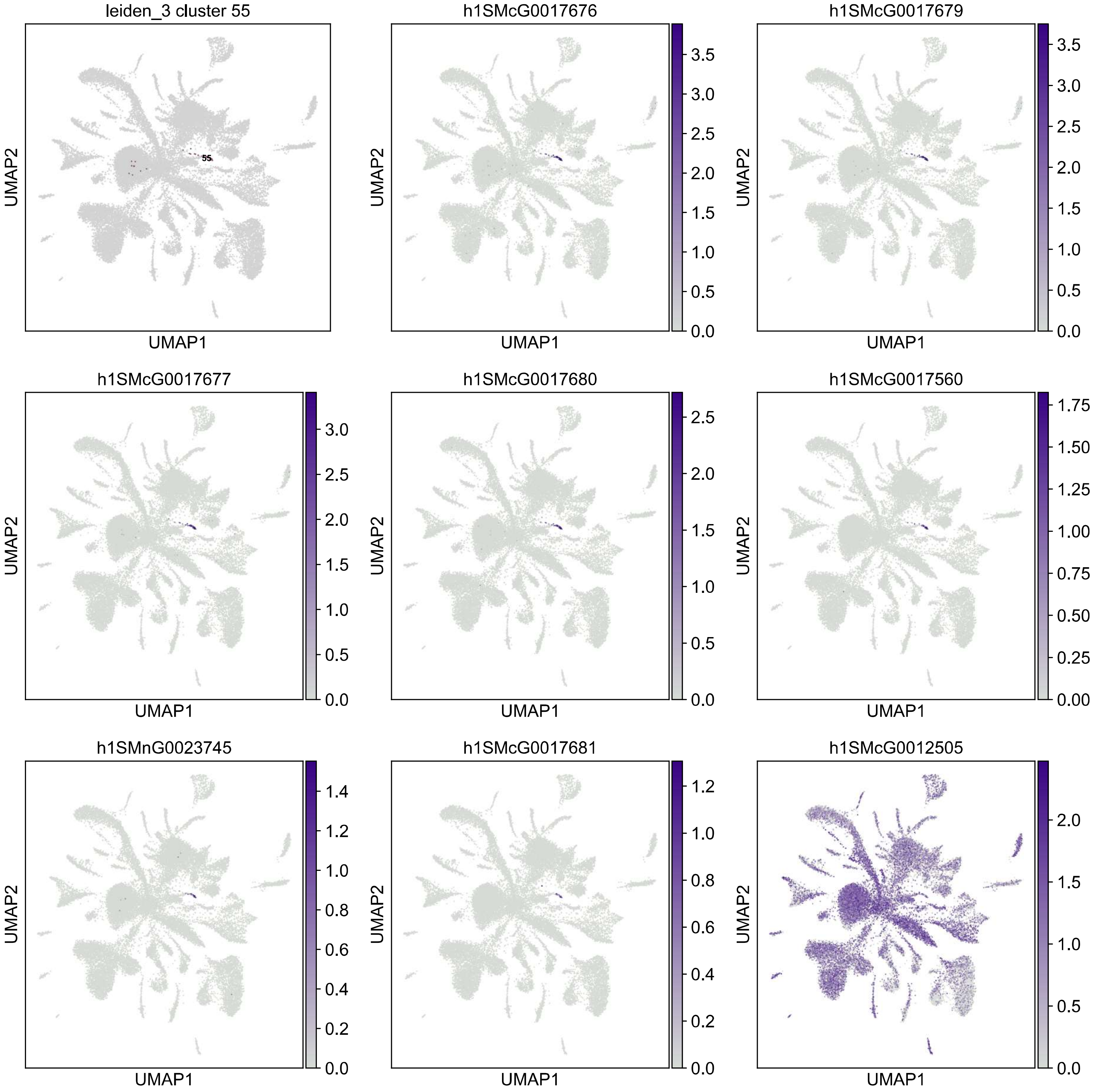

leiden_3 cluster 55
55
h1SMcG0017676
h1SMcG0017679
h1SMcG0017677
h1SMcG0017680
h1SMcG0017560
h1SMnG0023745
h1SMcG0017681
h1SMcG0012505
UMAP1
UMAP2

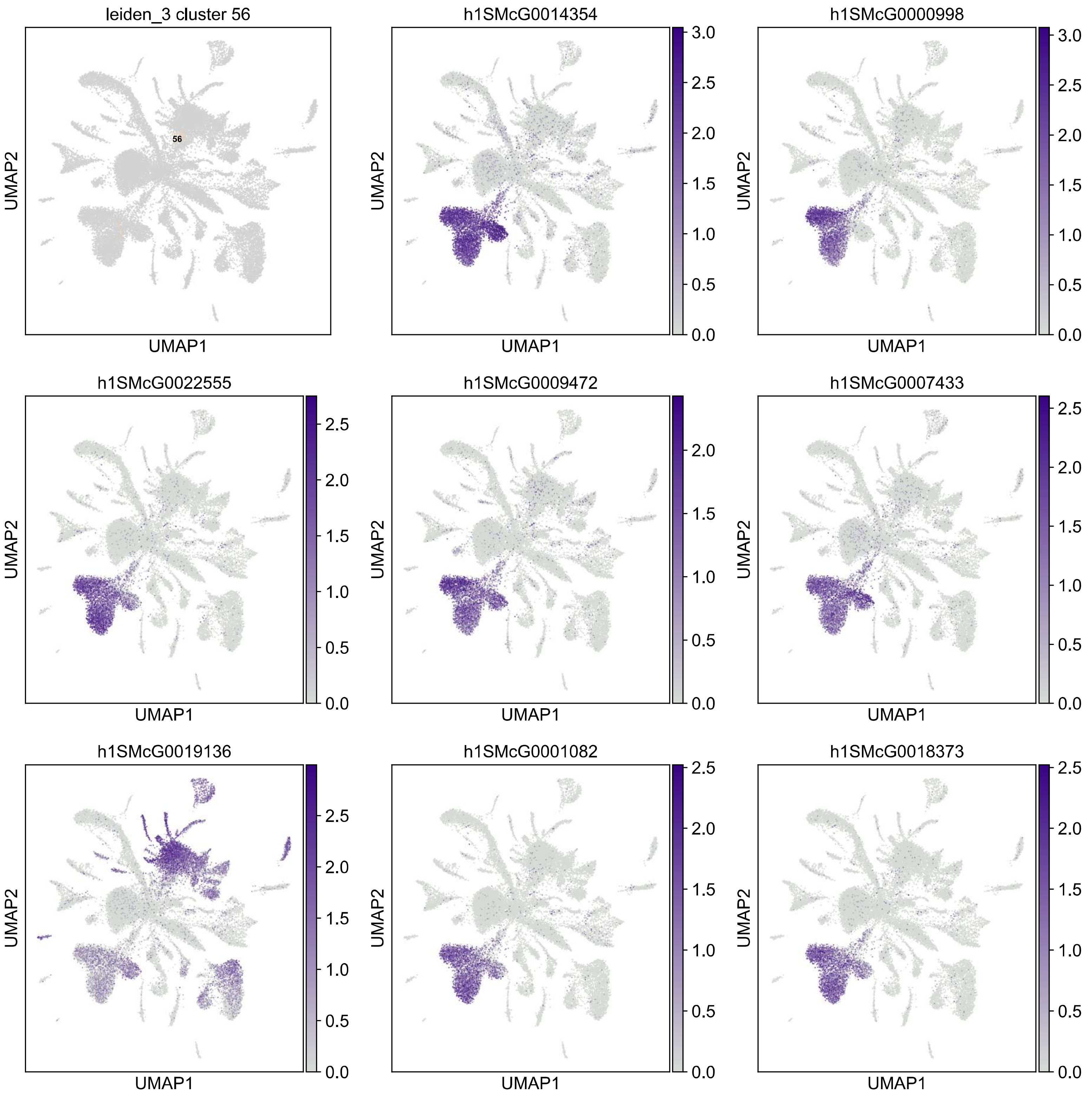

leiden_3 cluster 56
56
h1SMcG0014354
h1SMcG0000998
h1SMcG0022555
h1SMcG0009472
h1SMcG0007433
h1SMcG0019136
h1SMcG0001082
h1SMcG0018373
UMAP1
UMAP2

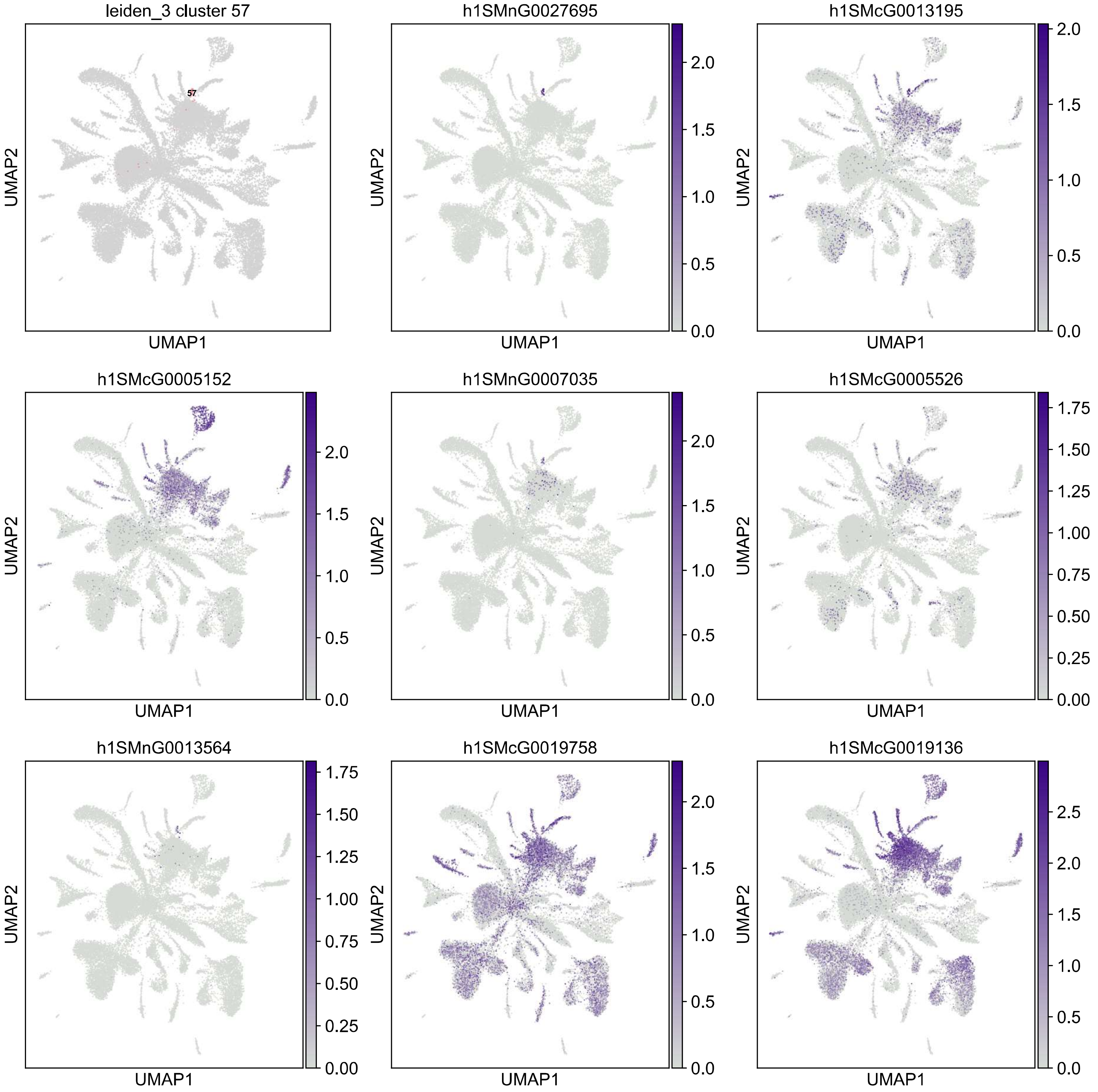
leiden_3 cluster 57
57
h1SMnG0027695
h1SMcG0013195
h1SMcG0005152
h1SMnG0007035
h1SMcG0005526
h1SMnG0013564
h1SMcG0019758
h1SMcG0019136
UMAP1
UMAP2

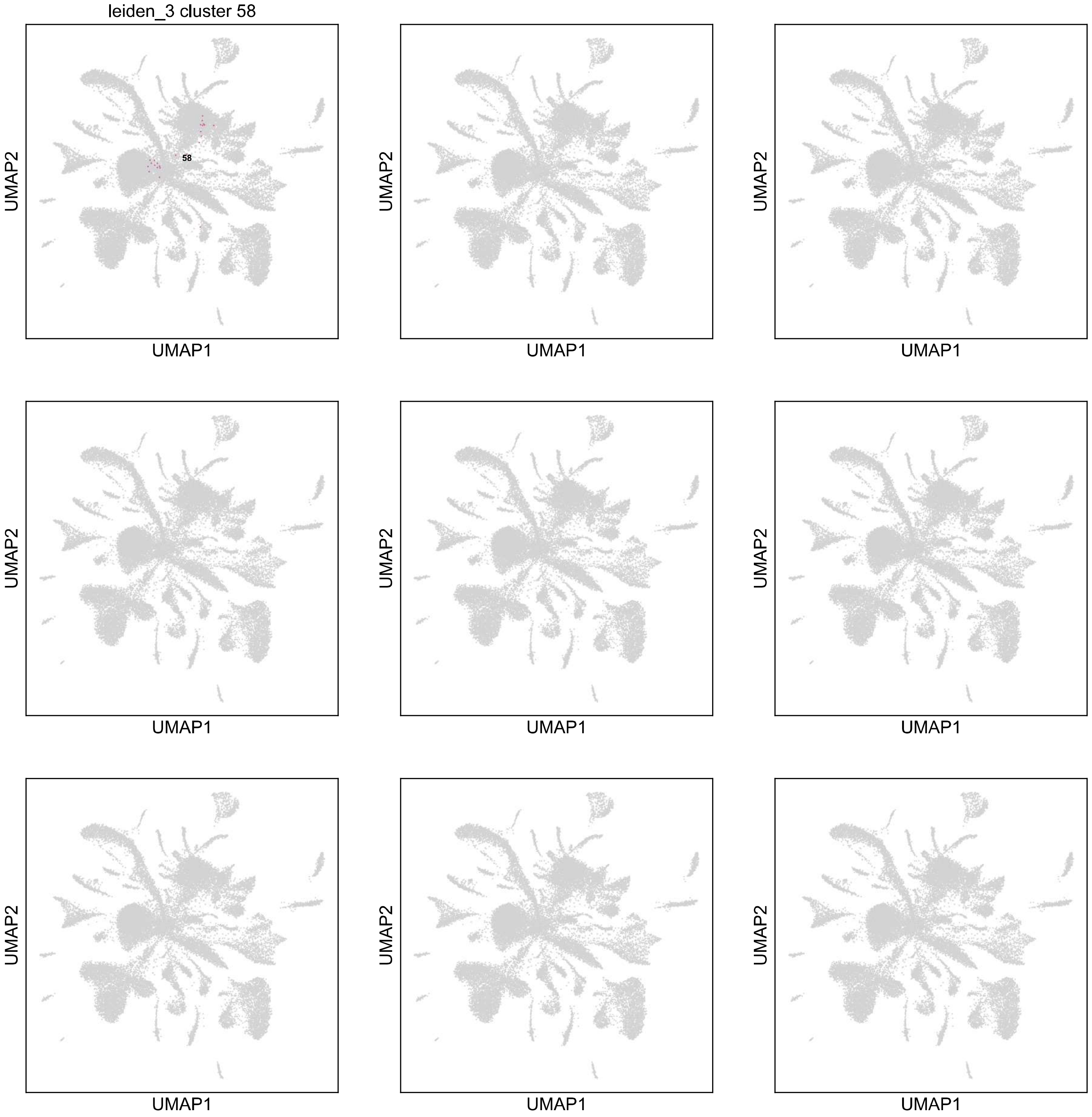
leiden_3 cluster 58
58
UMAP1
UMAP2

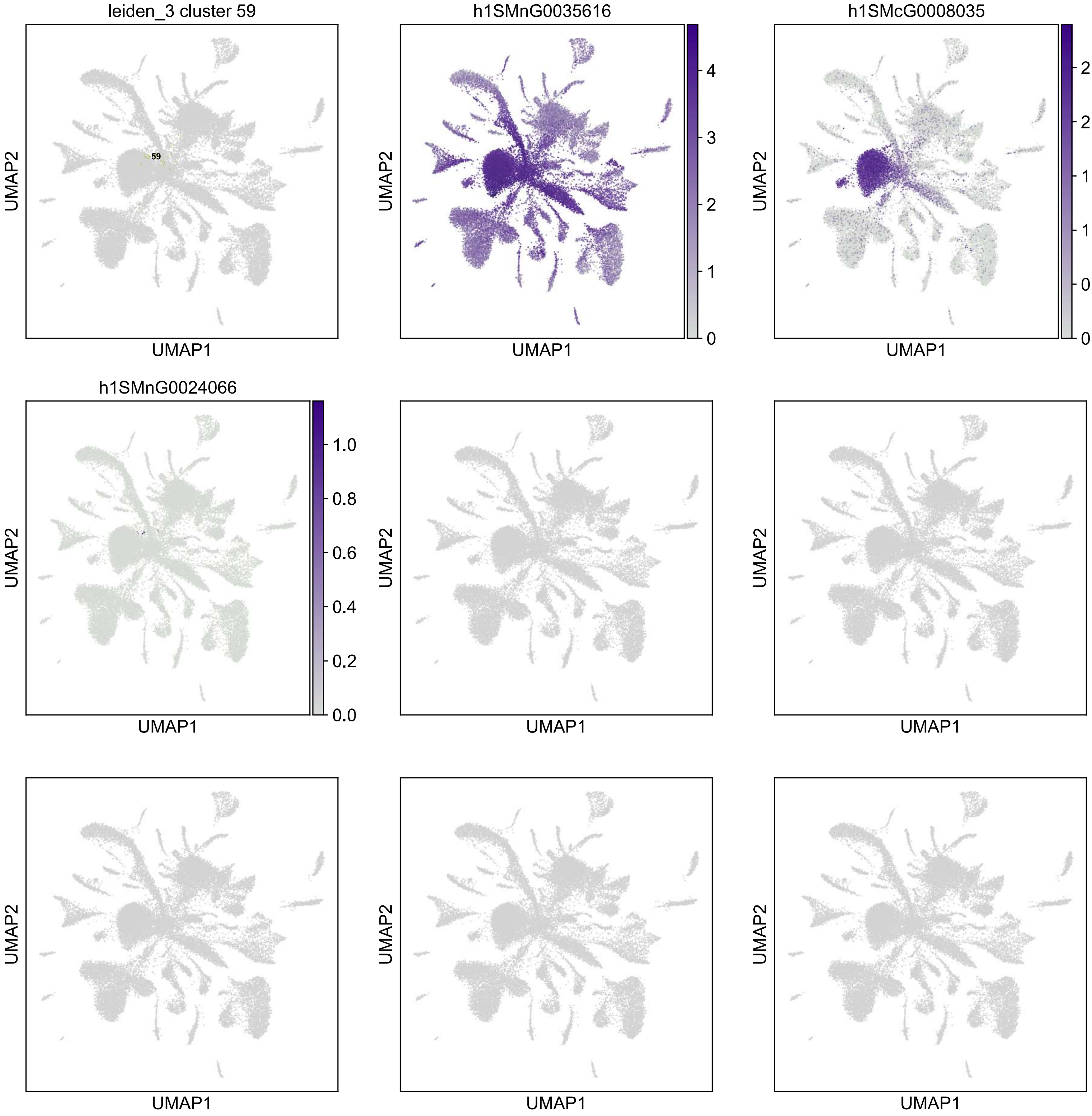
leiden_3 cluster 59
59
h1SMnG0035616
h1SMcG0008035
h1SMnG0024066
UMAP1
UMAP2
0
1
2
3
4
0.0
0.5
1.0
1.5
2.0
2.5
0.0
0.2
0.4
0.6
0.8
1.0

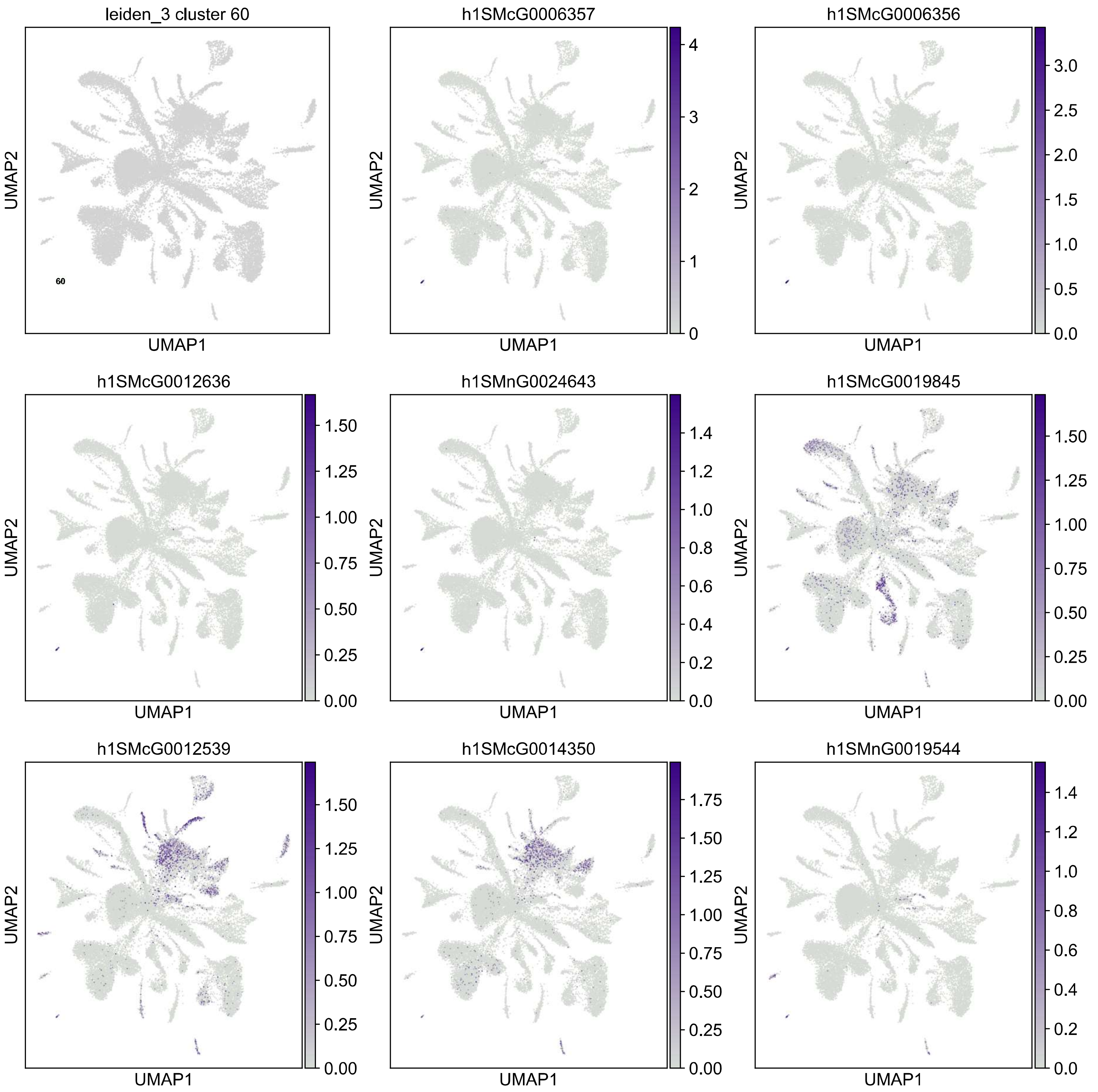

leiden_3 cluster 60
60
h1SMcG0006357
h1SMcG0006356
h1SMcG0012636
h1SMnG0024643
h1SMcG0019845
h1SMcG0012539
h1SMcG0014350
h1SMnG0019544
UMAP1
UMAP2

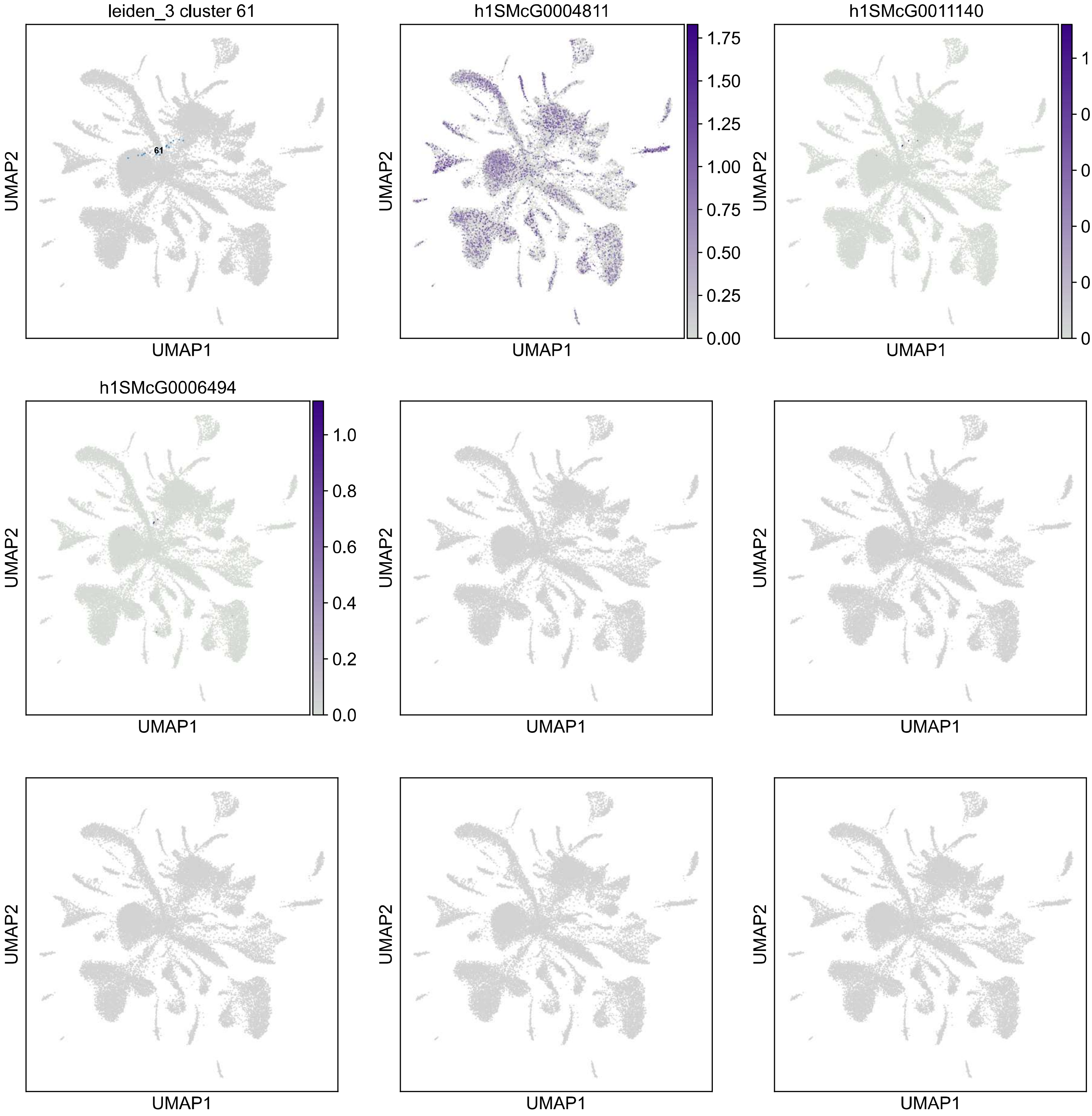

leiden_3 cluster 61
61
h1SMcG0004811
1.75
1.50
1.25
1.00
0.75
0.50
0.25
0.00
h1SMcG0011140
1.0
0.8
0.6
0.4
0.2
0.0
h1SMcG0006494
1.0
0.8
0.6
0.4
0.2
0.0
UMAP1
UMAP2

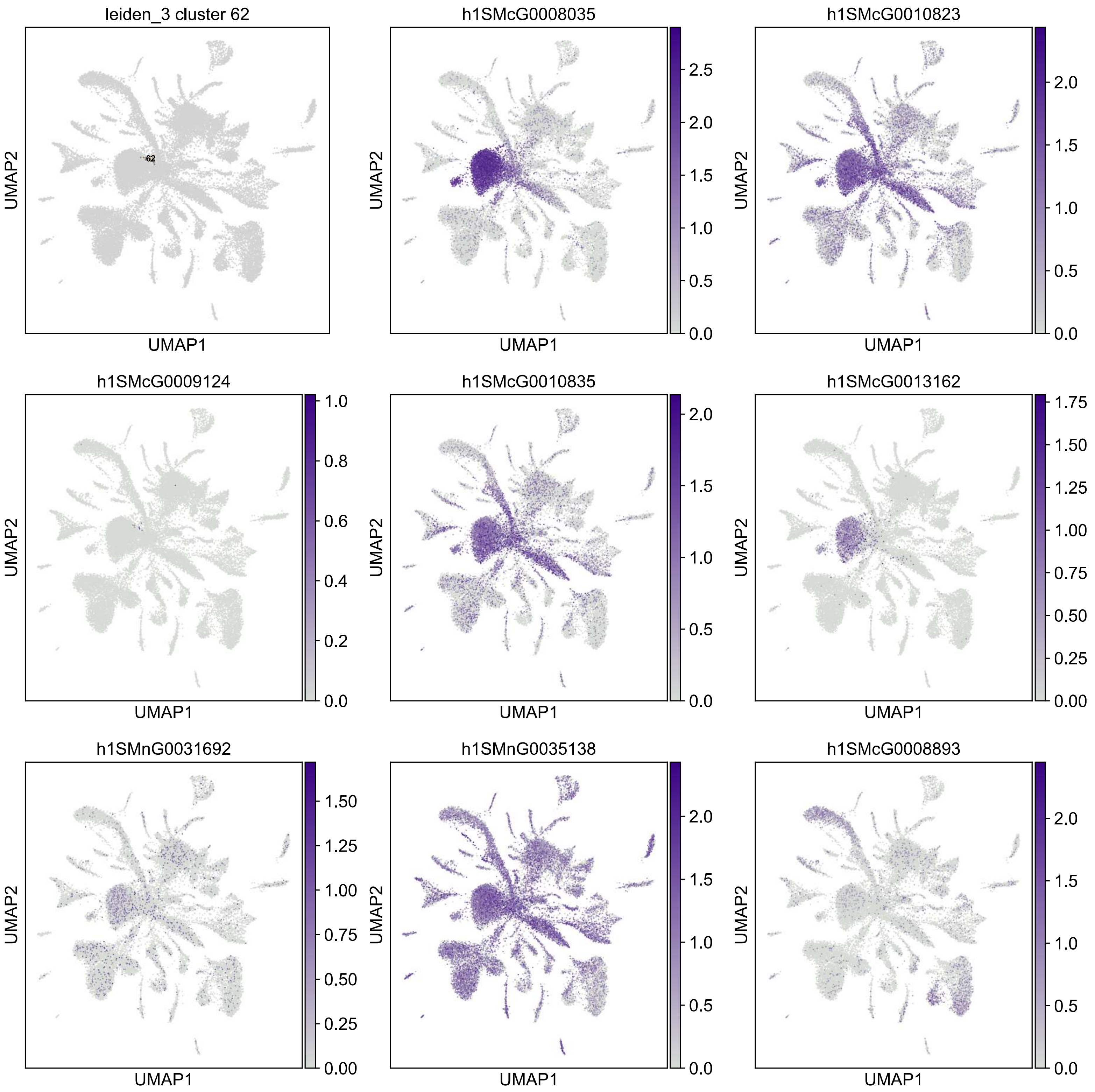
leiden_3 cluster 62
62
h1SMcG0008035
h1SMcG0010823
h1SMcG0009124
h1SMcG0010835
h1SMcG0013162
h1SMnG0031692
h1SMnG0035138
h1SMcG0008893
UMAP1
UMAP2

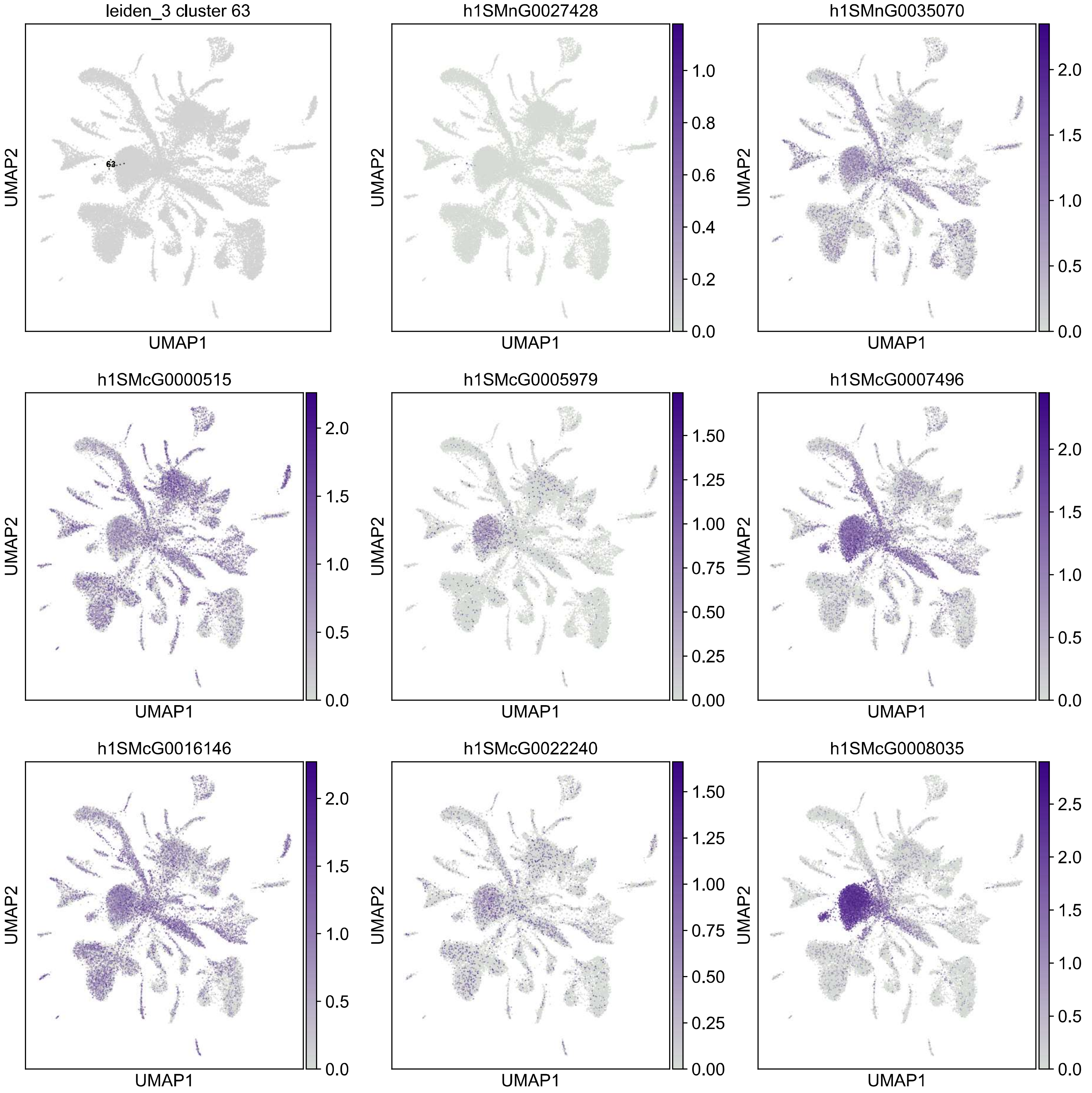
leiden_3 cluster 63
63
h1SMnG0027428
h1SMnG0035070
h1SMcG0000515
h1SMcG0005979
h1SMcG0007496
h1SMcG0016146
h1SMcG0022240
h1SMcG0008035
UMAP1
UMAP2
0.0
0.2
0.4
0.6
0.8
1.0
0.5
1.5
2.0
0.00
0.25
0.50
0.75
1.00
1.25
1.50
2.5

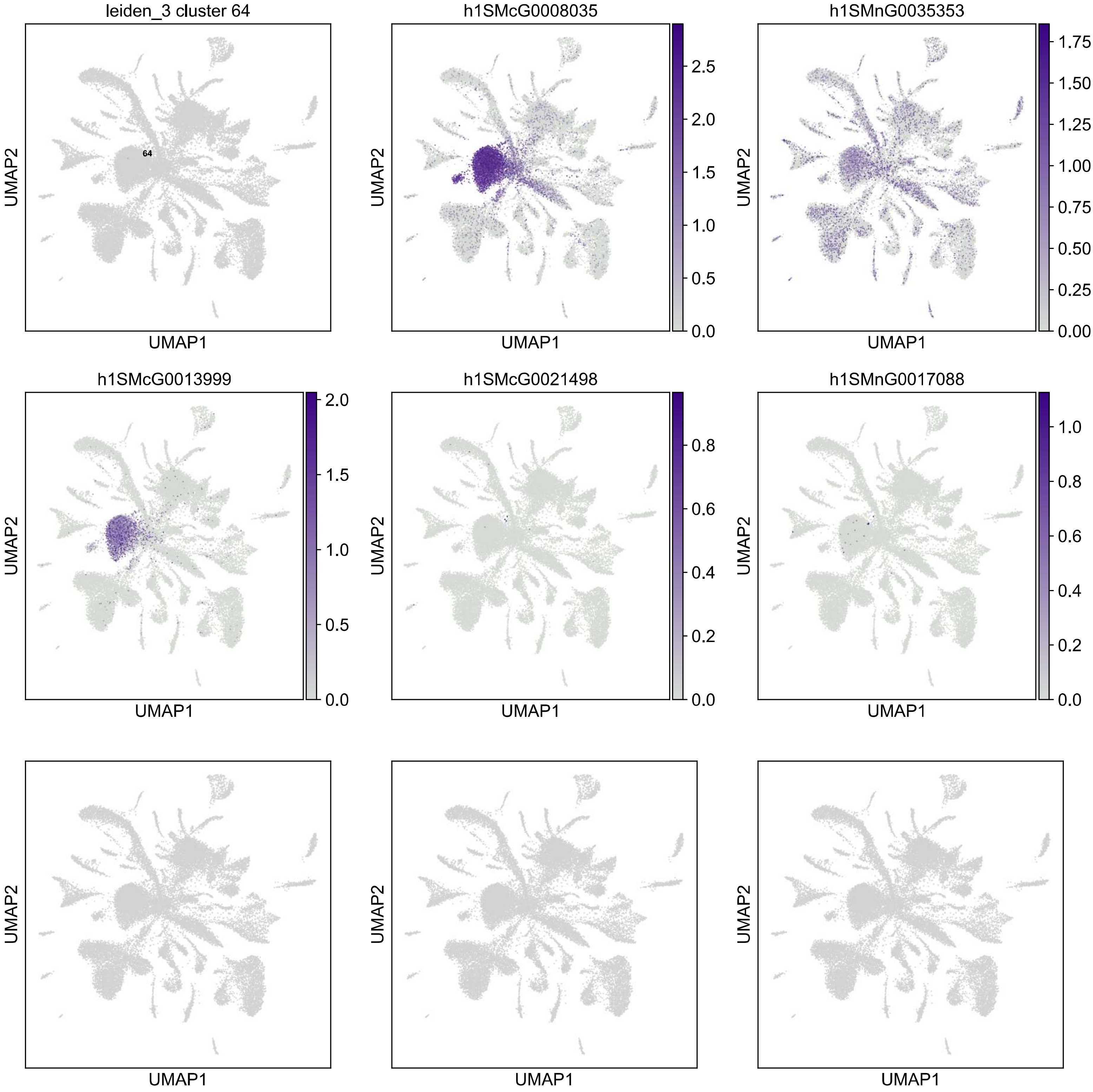

leiden_3 cluster 64
64
h1SMcG0008035
h1SMnG0035353
h1SMcG0013999
h1SMcG0021498
h1SMnG0017088
UMAP1
UMAP2
